# Catch bond models may explain how force amplifies TCR signaling and antigen discrimination

**DOI:** 10.1101/2022.01.17.476694

**Authors:** Hyun-Kyu Choi, Peiwen Cong, Chenghao Ge, Aswin Natarajan, Baoyu Liu, Yong Zhang, Kaitao Li, Muaz Nik Rushdi, Wei Chen, Jizhong Lou, Michelle Krogsgaard, Cheng Zhu

## Abstract

Central to T cell biology, the T cell receptor (TCR) integrates forces in its triggering process upon interaction with peptide-major histocompatibility complex (pMHC)^1-3^. Phenotypically, forces elicit TCR catch-slip bonds with strong pMHCs but slip-only bonds with weak pMHCs^4-10^. While such correlation is commonly observed, the quantitative bond pattern and degree of “catchiness” vary. We developed two models based on the structure, elastic properties, and force-induced conformational changes of the TCR–pMHC-I/II complexes to derive from their bond characteristics more intrinsic parameters that underlie structural mechanisms, predict T cell signaling, and discriminate antigens. Applying the models to 55 datasets of 12 αβTCRs and their mutants interacting with corresponding pMHCs without coreceptor engagement demonstrated the ability for structural and physical parameters to quantitatively integrate and classify a broad range of bond behaviors and biological activities. Comparing to the generic two-state model for catch-slip bond that also fits the data, our models can distinguish class I from class II MHC systems and their best-fit parameters correlate with the TCR/pMHC potency to trigger T cell activation, which the generic model cannot. The models were tested by mutagenesis using structural analysis, bond profile measurement, and functional assay of a MHC and a TCR mutated to alter conformation changes. The extensive comparisons between theory and experiment provided strong validation of the models and testable hypothesis regarding specific conformational changes that control bond profiles, thereby suggesting structural mechanisms for the inner workings of the TCR mechanosensing machinery and plausible explanation of why and how force may amplify TCR signaling and antigen discrimination.

## INTRODUCTION

Antigen recognition via TCR–pMHC interactions is essential for T cell activation, differentiation, proliferation, and function^11^. Mechanical forces applied to αβTCR via engaged pMHC substantially increase antigen sensitivity and amplify antigen discrimination^4-10,12^. As a fundamental force-elicited characteristic, strong cognate pMHCs form catch-slip bonds with TCR where bond lifetimes increase with force until reaching a peak, and decrease as force increases further, whereas weak agonist and antagonist pMHCs form slip-only bonds with TCR where bond lifetimes decrease monotonically with increasing force^4-10,13^. However, the mechanism underlying the correlation between the force-lifetime pattern and the ability for force on TCR to induce T cell signaling remains unclear.

An intuitive hypothesis is that catch bonds prolong interactions, which allow the process of CD3 signal initiation to proceed sufficient number of phosphorylation steps to the threshold for downstream signal propagation, as proposed by the kinetic proofreading model^14^. However, this hypothesis faces two challenges upon scrutiny of multiple TCR–pMHC systems. First, since bond lifetime *vs* force profiles are monotonically decreasing for slip-only bonds but bell-shaped for catch-slip bonds, the type of bonds that would last longer depends on the force range, which may be catch-slip bonds in one force regime but slip-only bonds in another force regime. Second, TCR–pMHC interactions exhibiting catch-slip bonds often have longest lifetimes around 10-20 pN (Ref. ^4-10,13^) and it has been reported that upon engaging pMHC, T cells would exert 12-19 pN endogenous forces on the TCR in a signaling-dependent fashion^15^. However, the relevance of this force range to T cell signaling remains incompletely understood. More perplexingly, some signal-inducing pMHCs form catch-slip bonds with TCRs but exhibit shorter lifetime than other pMHCs that do not induce signaling by, and form slip-only bonds with, the same TCRs even in the optimal force range^16^. These observations prompt the questions of what mechanism underlies the association of TCR–pMHC bond type with the T cell signaling capacity and what impact the 10-20 pN force range has on TCR mechanotransduction. To answer these questions requires an in-depth analysis of the multiple datasets with mathematical models, which was lacking.

Slip and catch bonds refer to two opposite effects of physical force on biomolecular interactions: increasing or decreasing their off-rate of dissociation, respectively^17,18^. Because force tends to be disruptive and destabilizing, slip bonds are intuitive, whereas catch bonds are counter-intuitive. Since excessive force can rupture even covalent bonds^19^, continued force increase will eventually overpower any catch bond, turning it to a slip bond after an “optimal” force where the off-rate is minimal^4-10,18^. Slip bond is commonly modeled by the Bell equation^20^, which assumes the off-rate *k* of a molecular bond dissociating along a single pathway in a one-dimensional (1D), single-well energy landscape to be an exponential function of force, 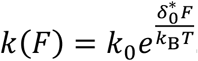. Here, *k*_0_ is the transition rate at zero force, *F* is tensile force, 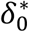 is the force-free distance from the bound state at the bottom of the energy well to the top of the energy barrier known as the “transition state”, *k*_B_ is the Boltzmann constant, and *T* absolute temperature^20^. Several models have been developed to account for catch-slip bond behavior. Most introduced two dissociation pathways and/or two bound states in a two-dimensional (2D) energy landscape that is tilted by force^21^ (Fig. 1a). One noticeable exception is that of Guo et al. where dissociation is modeled to start from a single bound state along a single pathway through a 1D energy landscape based on the physical process of peeling a polymer strand with force until the transition state is reached^22^. A distinct advantage of this model is its ability to relate the force-induced tilting of the energy landscape to the force-induced conformational change of the molecular complex (which all other models lack), thereby connecting parameters of the abstract energy landscape to the structural-elastic properties of the interacting molecules in question.

**Figure 1.**
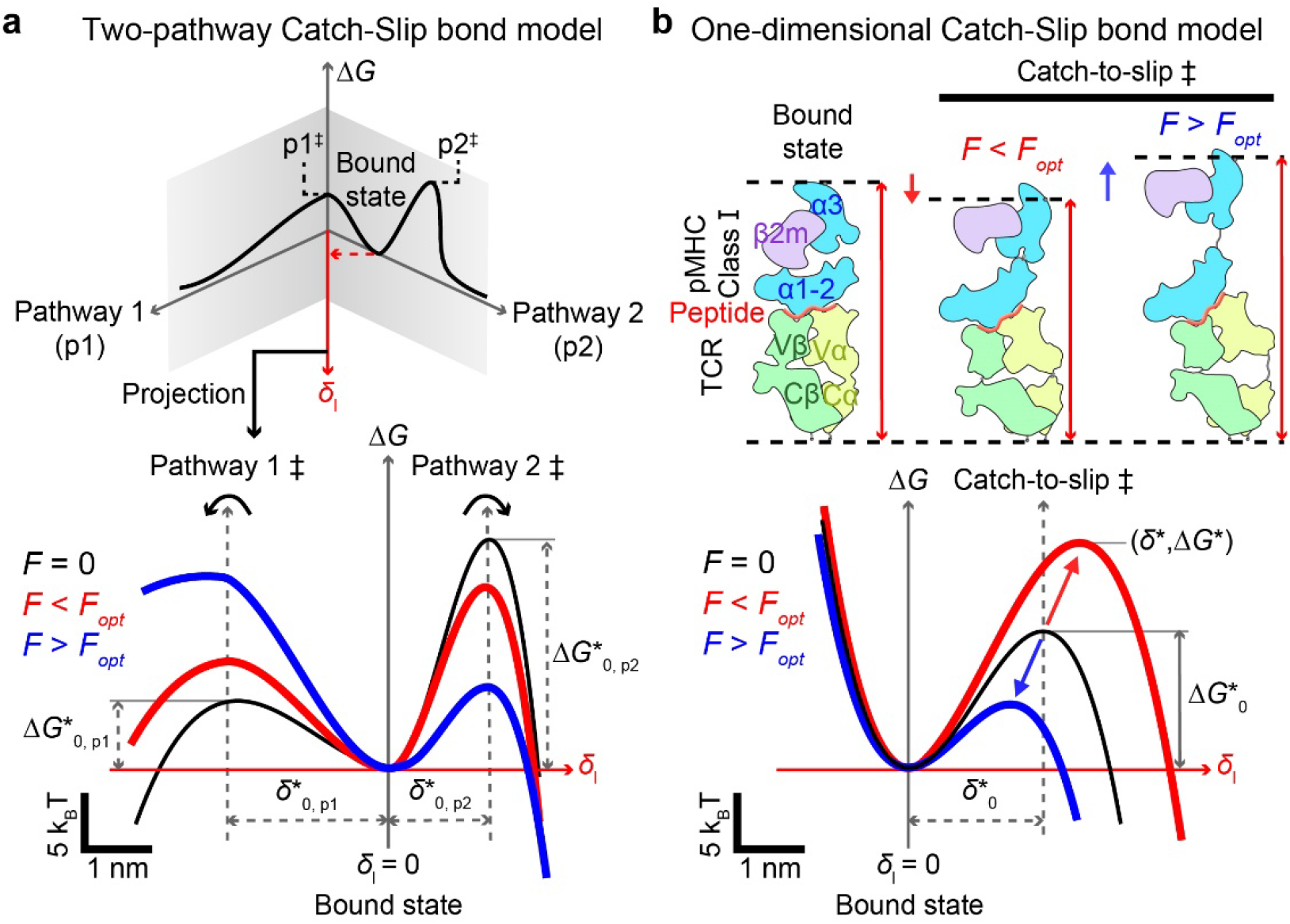
Comparison between the two-pathway model and the TCR–pMHC-I model for catch bond. **a** *Upper*: The 2D energy landscape of the two-pathway model where the bond is trapped in the bound-state energy well by two energy barriers that resist dissociation along two pathways, p1 and p2, with two distinct transition states, p1^‡^ and p2^‡^. The application of force projects the energy landscape towards a dissociation pathway along force (*δ*_1_). *Lower*: Force also tilts the energy-landscape, raising the energy barrier of the first pathway and lowering the energy barrier of the second pathway on the energy landscape projection (*red* and *blue*) relative to their positions in the absence of force (*black*). **b** *Lower*: The proposed 1D energy landscape of TCR–pMHC-I model with a single transition state *δ*^*^. Below an optimal value (*F*_opt_), force raises the energy barrier (*red*) relative to the zero-force conformation (*black*) by contraction of flexible regions due to entropic fluctuation. Above *F*_opt_, force lowers the energy barrier (*blue*) by stretching the molecular complex. Together, these two mechanisms give rise to a catch-slip bond. *Upper*: Schematics of the TCR–pMHC-I structure (*left*) and its conformational changes that correspond to low (*middle*) and high (*right*) forces.

Besides binding properties of the TCR–pMHC complex, its structural features and conformational changes have been suggested to be important for TCR triggering. For example, TCR–pMHC docking orientation has been correlated to its ability to trigger T cell signaling^16,23,24^. Partial unfolding or allosteric regulation of either the TCR and/or MHC molecules has been inferred from mechanical experiments and steered molecular dynamics (SMD) simulations of pulling single TCR–pMHC bonds^6,9,25,26^. Whereas the extent of these conformational changes has been correlated to the strength of TCR–pMHC catch bonds^6,9,25^, the two phenomena have not been integrated into a mathematical model to explore their potential connection.

Here we developed two such models, one for each MHC class, to describe both αβTCR catch-slip and slip-only bonds. The model development follows Kramers’ kinetic rate theory and uses polymer physics models to construct a 1D energy landscape for single-state, single-pathway dissociation that incorporates the structures, elastic properties, and force-induced conformational changes of the TCR–pMHC-I/II complexes at the sub-molecular level, which includes domain stretching, hinge rotation, and molecular extension. Incorporating the force-induced conformational changes into the energy landscape formulation allows force to shift the energy barrier up in low forces and down in high forces, thereby giving rise to catch-slop bonds along a single dissociation pathway in a 1D energy landscape (Fig. 1b). We applied our models and a published model^27^ to analyze 49 TCR–pMHC bond lifetime *vs* force datasets published to date in 9 papers by four laboratories measured using two different techniques^4-6,8,9,13,16,28,29^, plus 6 new datasets generated in this study, including 12 TCRs and their mutants expressed on the cell membrane or coated on beads interacting with corresponding panels of both classes of pMHCs without coreceptor engagement. This analysis demonstrated the superior power of our models’ structural and physical parameters to quantitatively integrate and classify a broad range of catch-slip and slip-only bond behaviors as well as their corresponding biological activities. Our models were rigorously validated by extensively comparing theory with experiment, testing the model assumptions and predictions, and using mutagenesis to alter specific conformational changes in the TCR–pMHC structure under force to modulate the catch bond profiles. By constructing the energy landscape underlying our models and investigating its properties, we obtained mechanistic insights into the inner workings of the TCR–pMHC mechanosensory machinery. By examining the correlation of the model parameters with the biological activities of a large number of TCR–pMHC-I/II systems, we explained how force-elicited catch bond may amplify TCR signaling and antigen discrimination.

## RESULTS

### Model development

#### Model goal

Kramers’ kinetic rate theory treats bond dissociation as state transition in a 1D energy landscape Δ*G*^*^(*δ*_1_) from a free-energy well (bound state) over a barrier (transition state) along the dissociation coordinate *δ*_1_^30^. Following Guo et al.^22^ to adapt the linear-cubic model of Dudko et al.^31^ but allow force *F* to tilt the original energy landscape by an amount of −*δ*_1_*γ*(*F*), the energy landscape takes the form of

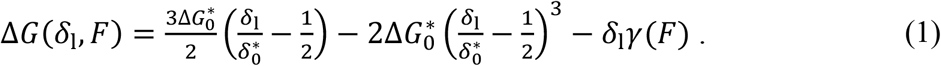

where 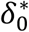 and 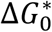 are the differences in dissociation coordinates and free-energy levels, respectively, between the transition state and bound state of the original force-free energy landscape. The corresponding force-dependent kinetic rate is

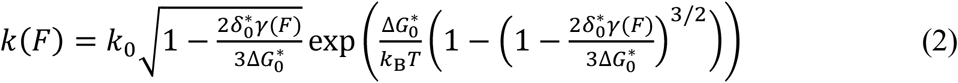

where *k*_0_ is the dissociation rate at zero force^22^. Letting *γ*∼*F* recovers from Eq. **2** the Dudko-Hummer-Szabo (DHS) model^31^, and further assuming 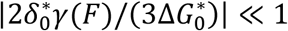 reduces it to the Bell model^20^. The condition for *k* to be able to model catch-slip bond is the derivative *k*^′^(*F*_0_) = 0 where *F*_0_ > 0. This translates to two conditions: the barrier height 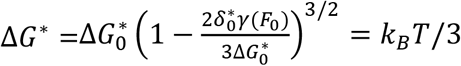 or *γ*^′^(*F*_0_) = 0. The first condition requires the energy change −*δ*_1_*γ* induced by *F* to lower the energy barrier height to *k*B*T*/3 located at 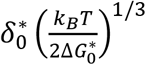. The second condition requires *γ* to be a biphasic function of *F*. This excludes the Bell model and the DHS model because both of their *γ* functions depend on *F* monotonically. Our goal is to construct a biphasic *γ*(*F*) with appropriate structural-elastic dependency to account for TCR– pMHC conformational changes from the bound state to the transition state, as analyzed by the biomembrane force probe (BFP) and optical tweezers (OT) experiments where a tensile force is applied to its two ends to modulate bond dissociation^4-10,13^.

#### Key model assumptions

Following the reasoning of Guo et al.^22^, 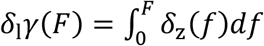 where the integrand *δ*_z_(*f*) = *z*(*f*) − *z*_0_(*f*) is the projection on the force direction of the change induced by force *f* of the TCR–pMHC extension at the transition state relative to its extension at the bound state. For *γ* to depend on *F* biphasically as required for describing catch-slip bonds, *δ*_z_ should be a biphasic function of *f* as discussed later. Therefore, dissociation occurs because the system moves in the energy landscape along the dissociation coordinate *δ*_1_ from the bound state to the transition state by a distance 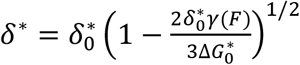 (Ref. ^22^). We assume that the differential contour length along the force transmission path across the TCR–pMHC structure (*i*.*e*., summing up all contour lengths of various domains connected at nodes of force action, as depicted in red lines in Fig. 2a, for MHC-I) at the transition state *l* and bound state *l*_0_ can serve as a dissociation coordinate, i.e., *δ*_1_ = *l* − *l*_0_. *δ*_z_ is the projection of *δ*_1_ on the z axis – the direction of the pulling force (Fig. 2a). When only contour lengths are considered, 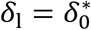, which serves as a criterion for finding the best-fit parameters (Fig. 2a, *left* and Supplementary Model Derivations, Eq. S7).

**Figure 2.**
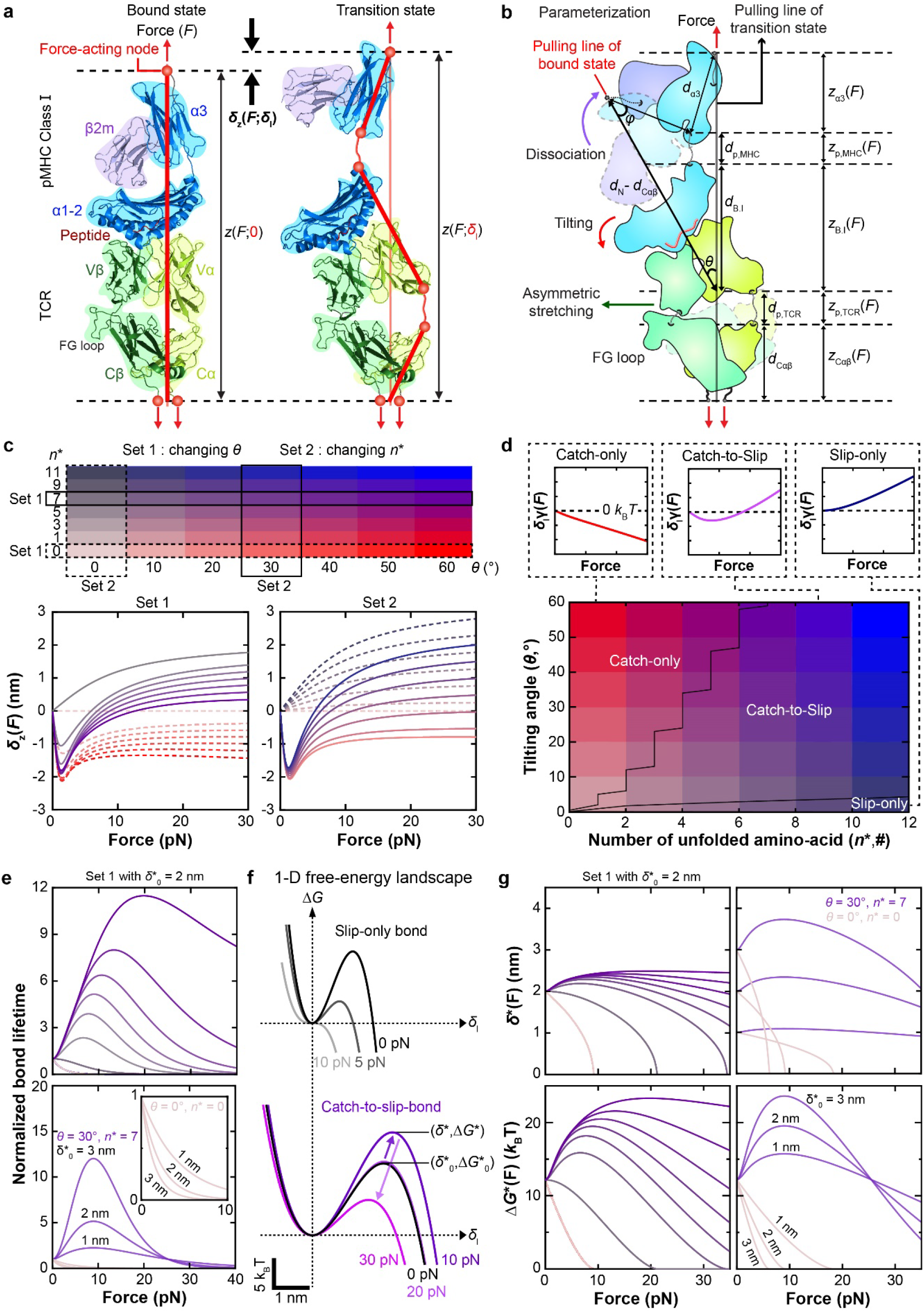
Structure, mechanics, and characteristics of the TCR–pMHC-I catch bond model. **a** Force-induced conformational changes of a TCR–pMHC-I complex as it traverses from the bound state (*left*) to the transition state (*right*). The diagrams of the 2C TCR α (*yellow*) β (*green*) subunits and the DEVA peptide (*red*) bound to the H2-K^b^ (various domains indicated) are based on snapshots from SMD simulations performed on the complex structure (2CKB) at the initial time (bound state) and a later time (transition state)^9^. The force-transmission path is shown as red lines connecting the force-acting nodes. **b** Various contributions to the total extension projected on the force axis: rotation of the α_3_-β_2m_ domains about the MHC C-terminus upon dissociation of the β_2m_–α_1_α_2_ interdomain bond (*z*_α3_), relative rotation between α_3_ and α_1_α_2_ about their stretched interdomain hinge (*z*_p,MHC_), tilting of the MHC α_1_α_2_ complexed with TCR Vαβ (*z*_B.I_), rotation about and extension of the Vα–Cα interdomain hinge (*z*_p,TCR_), and extension of the Cαβ and rotation about their C-termini (*z*_Cαβ_). Two α_3_-β_2m_ structures are shown: before (light colors) and after (dark colors) β_2m_ dissociation from α_1_α_2_., with two parameters describing their contributions to the total extension: *d*_*α*3_ = the distance between the α3 C- and N-termini excluding the α_1_α_2_–α3 hinge and *θ* = the angle between the normal direction of the TCR–pMHC bonding interface at the bound state (cf. (**a**), *left*) and the tilted direction at the transition state (cf. (**a**), *right*). **c** Extension change *vs* force curves (*lower*) for the color-matched *n*^*^ and *θ* values (*upper*). The left panel (set 1 in *upper* table) shows the effect of changing *θ* with (*n*^*^ = 7) and without (*n*^*^ = 0) partial unfolding. The right panel (set 2) shows the effect of changing *n*^*^ with (*θ* = 30°) and without (*θ* = 0°) tilting. **d** *n*^*^-*θ* phase diagram showing three parameter domains: slip-only, catch-slip, and catch-only respectively colored by red-purple, purple-blue, and blue-black. Upper insets indicate corresponding energy change *δ*_1_*γ vs* force curves for each bond type. **e** Theoretical normalized bond lifetime *vs* force curves for indicated parameters. The upper and lower panels show the respective effects of changing *θ* and 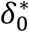 from the set 1 parameters defined in (**c**). **f** Energy landscapes expressed as families Δ*G vs δ*_*l*_ curves for a range of forces for slip-only (*upper*) and catch-slip (*lower*) bonds. The bound state is located at the origin Δ*G* = 0 and *δ*_*l*_ = 0. The transition state has an energy of Δ*G*^*^ located at *δ*^*^ when *F* > 0 and 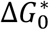 located at 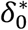 when *F* = 0.**g** Plots of transition state location *δ*^*^ (*upper*) and height of energy barrier Δ*G*^*^ (*lower*) *vs* force *F* for changing *θ* while keeping 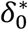 constant (*left*) or changing 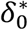 while keeping *θ* constant (*right*) for the indicated values from parameter table in (**c**).

Suggested by single-molecule OT^6^, BFP^9^ and magnetic tweezers^9^ experiments as well as steered MD (SMD) simulations^9^, we assume that force-induced TCR–pMHC dissociation is accompanied by conformational changes in the TCR, MHC, or both. Specifically, we assume that at the bound state, force induces elastic extension of the TCR–pMHC structure as a whole (Fig. 2a, *left*); but as the system moves toward the transition state for dissociation, conformational changes may occur, which may include disruption of intramolecular interfaces, hinge rotation, and partial unfolding of interdomain joints (Fig. 2a, *right*). To include appropriate details of these proposed conformational changes at the sub-molecular level into the expression of *δ*_z_, we model the TCR–pMHC structure as a system of semi-rigid bodies representing the whole complex as well as various globular domains connected by semi-flexible polymers that allow extension and hinge rotation under mechanical loads (Fig. 2a, *right*). Specifically, we assume that, as the system moves along the dissociation pathway from the bound state toward the transition state, force may induce disruption of the MHC α_1_α_2_–β_2m_ interdomain bond, thereby shifting the mechanical load originally borne by this bond to the α_1_α_2_–α_3_ joint to induce its partial unfolding, as observed in SMD simulations^9^. As such, the MHC α_3_ domain would change its length and rotate about its C-terminus (Fig. 2b). Since the TCRβ subunit has also been proposed to undergo FG-loop-regulated conformational change^6,25,26^, we assume that disruption of the α_1_α_2_–β_2m_ joint would also result in tilting of the TCR–pMHC bonding interface and shifting of the mechanical load from the TCR Vβ–Cβ joint to the Vα–Cα joint, leading to partial unfolding of the Vα–Cα joint (Fig. 2b). This increased stretching of the Vα–Cα joint relative to the Vβ–Cβ joint is assumed to result from strengthening of the Vβ–Cβ joint by the FG-loop^12^. At the transition state, therefore, we treat the MHC α_3_ domain (*d*_α3_), the MHC α_1_α_2_ domains bound to the TCR Vαβ domains (*d*_B.I_), and the TCR Cαβ domains (*d*_Cαβ_) as three semi-rigid bodies connected by two unfolded peptide chains of the MHC α_1_α_2_–α_3_ joint (*d*_p,MHC_) and the TCR Vα–Cα joint (*d*_p,TCR_) (Fig. 2b, *right*). At the bound state, neither disruption of intramolecular bonds nor partial unfolding of interdomain joints occurs, as mentioned earlier, allowing the whole TCR–pMHC ectodomain (ECD) complex to be modeled as one semi-rigid body (*d*_N_).

#### Force-induced energy change

To derive an expression for the last term on the right-hand side of Eq. 1, we model the semi-rigid bodies *d*_i_ (*i* = N, α_3_, B.I., and Cαβ to respectively denote the whole TCR–pMHC ECD structure as well as its indicated domains) as three-dimensional freely-jointed chains (FJC) and employ polymer physics to obtain their force-extended length *d*_i_(*f*) from their force-free length *d*_i,c_ (Ref. ^32^) (Supplementary Model Derivations, Eq. S1).

The assumed partial unfolding of the α_1_α_2_–α_3_ joint and the Vα–Cα joint are based on suggestions from single-molecule OT^6^, BFP^9^ and magnetic tweezers^9^ experiments as well as SMD simulations^9^. We model these unstructured polypeptides as extensible worm-like chains (eWLC) and employ polymer physics to obtain their force-induced extension *d*_p,i_(*f*) (*i* = MHC and TCR) from their force-free, folded state, which has zero length^33^ (Supplementary Model Derivations, Eq. S3).

Upon projecting the various force-induced extensions described above onto the force axis, we obtain *z* components of five contributions to the TCR–pMHC length increase at the transition state: extension of the MHC α_3_ domain (*z*_*α*3_), unfolding of the MHC α_1_α_2_–α_3_ interdomain joint (*z*_p,MHC_), extension of bonding interface that includes the MHC α_1_α_2_ domains bound to the TCR Vαβ domains (*z*_B.I_), unfolding of the Vα–Cα joint (*z*_p,TCR_), and extension of the TCR Cαβ domains (*z*_Cαβ_) (Fig. 2b). Finally, we obtain:

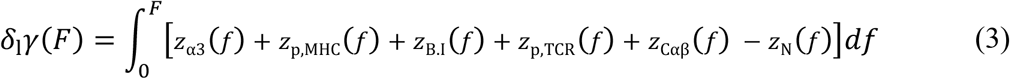

### Model characterization

#### Model constants and parameters

The FJC model constants for the 1^st^, 3^rd^, 5^th^, and 6^th^ terms in the integrand on the right-hand side of Eq. 3 include the force-free lengths *d*_i,c_ and the elastic modulus of the folded globular domains *E*_c_, all of which are available from the literature. The 2^nd^ and 4^th^ terms are proportional the respective numbers of amino acids in the polypeptides of the partially unfolded MHC α_1_α_2_–α_3_ joint (*n*_p,MHC_) and TCR Vα–Cα joint (*n*_p,TCR_), which can be combined as the product of the total unfolded amino acid number *n*^*^ = *n*_p,MHC_ + *n*_p,TCR_, the average contour length per unfolded amino acid *l*_c_, and the extension per unit contour length *z*_u,p_(*f*). The eWLC model constants for *z*_u,p_(*f*) include the average persistence length per unfolded amino acid *l*_p_ and the elastic modulus of polypeptides *E*_p_ (Supplementary Table 1).

After applying model constraints and the approximating 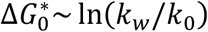 where *k*_*w*_ ∼10^6^ s^-1^ is known as the prefactor (Supplementary Model Derivations), the model parameters are reduced to five: three structural parameters (*d*_α3_,*θ, n*^*^) and two biophysical parameters 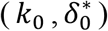, for describing dissociation of TCR–pMHC-I bonds. We will determine these parameters by comparing the model predictions with experimental measurements, and by doing so, illustrate the ability of our model to use a relatively low number of parameters to capture the coarse-grained structure and conformational changes at the sub-molecular level during TCR–pMHC-I dissociation.

#### Model features and properties

To explore the general features and properties of the model, we plotted *δ*_z_ vs *F* for two *n*^*^ values and a range of *θ* values as well as two *θ* values and a range of *n*^*^ values (Fig. 2c). Conceptually, force-heightened energy barrier generates catch bonds and force-lowered energy barrier produces slip bonds (Fig. 2d, *top left* and *right* panels). Since −*δ*_1_*γ* represents the energy input by force *F* to the original energy landscape, a biphasic *δ*_1_*γ* is required to create catch-slip bonds (Fig. 2d, *top middle* panel); correspondingly, *δ*_z_ is required to have a root at positive *F* where catch bond transitions to slip bond (Fig. 2c). The parameter domains capable of generating catch, catch-slip, and slip bonds are mapped on an *n*^*^-*θ* phase diagram (Fig. 2d), showing that our model can describe catch-slop bond if and only if *n*^*^ > 0, *θ* > 0, and *δ*_1_*γ*(∞) > 0 (Fig. 2d, *top middle* panel). In other words, catch-slip bonds require partial unfolding of the MHC α_1_α_2_–α_3_ and/or TCR Vα–Cα joints and tilting of the TCR–pMHC bonding interface, a prediction consistent with previous results of SMD simulations and single-molecule experiments^9^.

For single-bond dissociation from a single bound state along a single pathway, the reciprocal dissociation rate should be equal to the average bond lifetime. Regardless of the bond type, the reciprocal zero-force off-rate controls the *y*-intercept of the bond lifetime *vs* force curves. We plotted the theoretical bond lifetime (normalized by its zero-force value) *k*_0_/*k vs* force *F* for a range of *n*^*^, *θ*, and 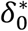 to examine how the model parameters control the bond lifetime *vs* force profile (Fig. 2e). Consistent with Fig. 2c-d, only if *n*^*^ > 0 and *θ* > 0 can our model describe catch-slip bond. Increasing the tilting angle *θ* results in more pronounced catch-slip bonds with longer lifetimes that peak at higher forces (Fig. 2e, *upper panel*). By comparison, increasing 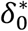 changes the level of slip-only bonds if *n*^*^ = 0 and *θ* = 0, but prolongs lifetime of catch-slip bonds (until cross-over at a higher force) without changing the force where lifetime peaks if *n*^*^ > 0 and *θ* > 0 (Fig. 2e, *lower panel*).

To understand physically how our model describes catch-slip bonds, we plotted the energy landscape using Eq. 1 (Fig. 2f). Setting *θ* = 0 in the upper panel generates a family of Δ*G vs δ*_1_ curves where the energy barrier is suppressed monotonically with increasing force, indicating a slip-only bond (Fig. 2f, *upper panel*). By comparison, setting *θ* > 0 results in a family of Δ*G vs δ*_1_ curves where the energy barrier height is initially raised, then lowered, by increasing force, indicating a catch-slip bond (Fig. 2f, *lower panel*, also see Fig. 1b). We also examine how the transition state location (Fig. 2g, *upper row*) and energy barrier height (Fig. 2g, *lower row*) change with force for a range of *θ* and 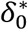 values that give rise to slip-only bonds and catch-slip bonds. Noticeably, at fixed *θ* values, both rates by which the transition state location and the energy barrier height change with force are accelerated by increasing 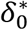 (Fig. 2g, *right column*), suggesting that this parameter can be used as a measure for force sensitivity. Interestingly, increasing *θ* slows the decrease in both the transition state location and energy barrier height with force at higher values, suggesting that the tilting angle controls the range at which force sensitivity can last (Fig. 2g, *left column*).

### Model validation

#### Model’s capability to fit data

To test the validity of our model, we used it to analyze 9 class I-restricted TCRs and their mutants (MT) either expressed on primary T cells or hybridomas with CD3s, or purified ECD without CD3s, which form catch-slip bonds and slip-only bonds with their respective specific peptides presented by wild-type (WT) or MT MHCs, consisting of 42 datasets published by 4 labs and a new dataset. We re-analyzed a TCR system published by the Zhu lab: the murine OT1 TCR expressed on either primary naïve CD8^+^ T cells or CD4^+^CD8^+^ thymocytes, or soluble TCRαβ ECD, which interacted with various peptides presented by a MT MHC (H2-K^b^α3A2) that abolished CD8 co-engagement^4,8^ (9 datasets, Fig. 3a and Supplementary Fig. 1a). We also re-analyzed two TCR systems published by the Zhu lab and Chen lab: WT or 2 MT murine 2C TCRs expressed on either primary naïve CD8^+^ T cells, CD4^+^CD8^+^ thymocytes or CD8^-^ hybridoma cells, or soluble TCRαβ ECD, which interacted with various peptides presented by H2-K^b^α3A2 (for CD8^+^ primary T cells, CD4^+^CD8^+^ thymocytes, or soluble TCR) or H2-K^b^ (for CD8^-^ hybridoma cells) without or with two point mutations specifically designed to alter bond profile, or by H2-L^d^(m31), a different MHC allele from H2-K^b^ (Ref. ^9^) (19 datasets, Fig. 3b, Supplementary Fig. 1b-d), and WT or 3 MT human 1G4 TCRs expressed on hybridoma cells, which interacted with the melanoma peptide NY-ESO-1 bound to HLA-A2 (Ref. ^9^) (4 datasets, Supplementary Fig. 1e). Furthermore, we re-analyzed five TCR systems published by the Evavold lab: the murine P14 TCR expressed on primary naïve CD8^+^ T cells, which interacted with various peptides presented by H2-D^b^ with a D227K point mutation to abrogate CD8 binding^29^ (3 datasets, Fig. 3f, Supplementary Fig. 1f), and four mouse TCRs expressed on hybridomas interacted with NP_366_ bound to the D227K mutant of H-2D^b^ to prevent CD8 binding^16^ (4 datasets, Supplementary Fig. 1g). Moreover, we fitted our model to a TCR system published by the Lang lab: the soluble mouse N15 TCRαβ interacting with VSV and two MT peptides bound to H2-K^b^ (Ref. ^6^) (3 datasets, Supplementary Fig. 1h). In addition, we performed a new experiment specifically designed to test our model prediction that destabilizing the α_1_α_2_–β_2_m interdomain bond of H2-K^b^ would amplify TCR–pMHC catch bond (see Fig. 2a), which measured 2C TCR interaction with the same peptide (R4) presented by the WT H2-K^b^α3A2 instead of the hybrid H2-K^b^α3A2 that swaps the mouse β_2_m with the human β_2_m (see Fig. 7a-c below) (1 dataset, Fig. 3b and Supplementary 1b). Gratifyingly, the theoretical reciprocal force-dependent off-rate 1/*k*(*F*) fits all 43 experimental bond lifetime *vs* force curves well (Fig. 3a-b, Supplementary Fig. 1, and Supplementary Tables 2 and 3), demonstrating our model’s capability to describe a wide range of data.

**Figure 3.**
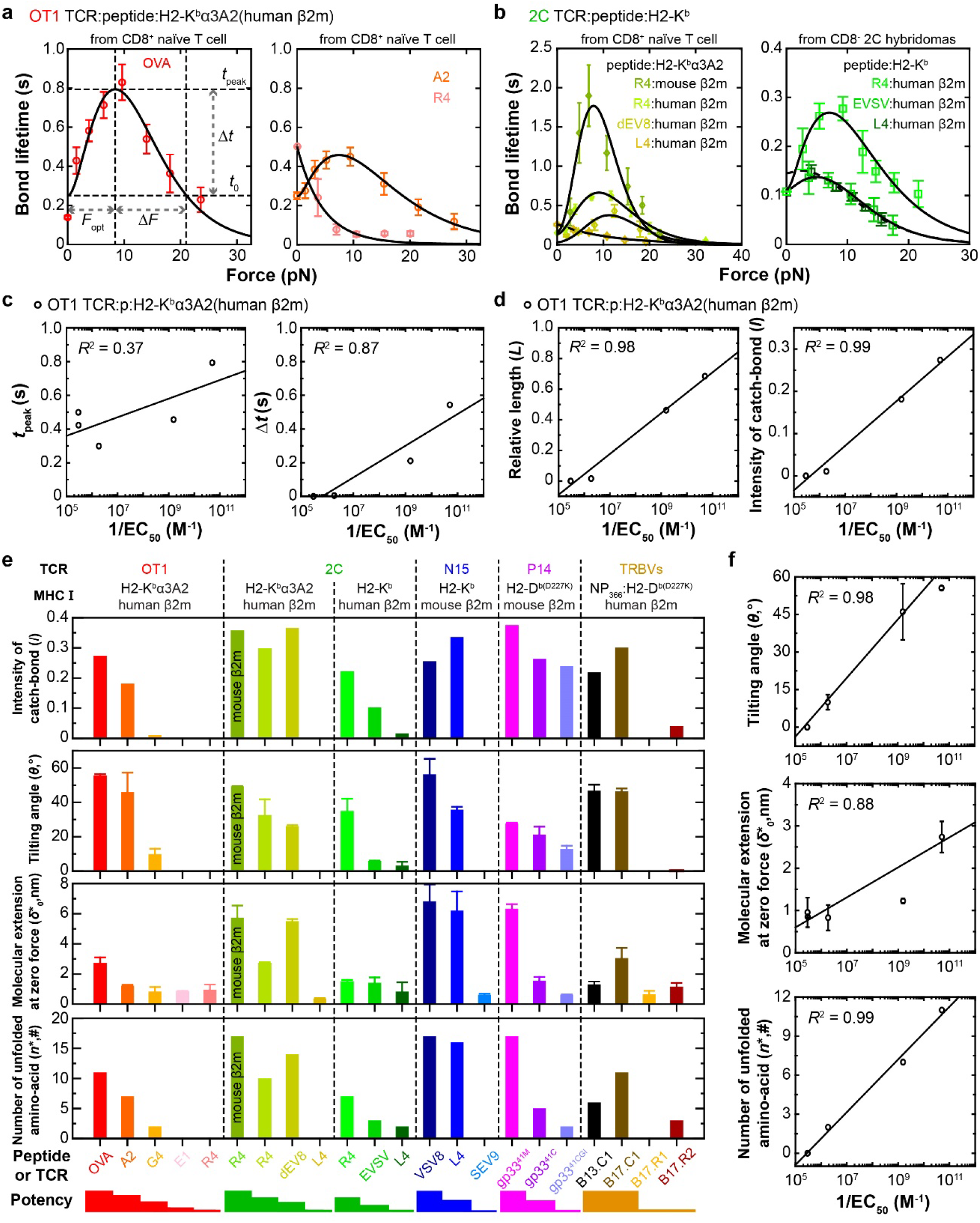
TCR bond type characterization and correlation with pMHC-I biological activity. **a**-**b** Fitting of theoretical 1/*k*(*F*) curves to experimental bond lifetime *vs* force data (*points*, mean ± sem, re-analyzed from Refs. ^4,9^) of OT1 (**a**) or 2C (**b**) TCR expressed on CD8^+^ naïve T cells interacting with indicated p:H2-K^b^α3A2 ((**a**) and (**b**) *left*) or on CD8^-^ 2C hybridomas interacting with indicated p:H2-K^b^ ((**b**) *right*). Several metrics are defined to characterize the force-lifetime curve as indicated in the left panel of (**a**): *F*_opt_ is the “optimal force” where lifetime peaks (*t*_peak_), Δ*t* is the lifetime increase from the zero-force value *t*_0_ to *t*_peak_, and Δ*F* is the force increase from *F*_opt_ to the force where lifetime returns to *t*_0_. **c**-**d** Two dimensional metrics, *t*_peak_ and Δ*t* (**c**), and two dimensionless metrics, *L* = Δ*t*/*t*_peak_ and *I* = *L*/(1 + *B*) where *B* = Δ*F*/*F*_opt_ (**d**), are plotted *vs* the logarithm of the reciprocal peptide concentration required to stimulate half-maximal T cell proliferation (1/EC_50_) and fitted by a linear function. **e** Intensity of catch bond (*I*, 1^st^ row), best-fit model parameters *θ* (the tilted angle of the bonding interface, 2^nd^ row), 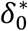 (the width of zero-force free-energy well, 3^rd^ row), and *n*^*^ (the number of unfolded amino acids, 4^th^ row) derived from the fitted force-lifetime curves of OT1, 2C TCR on primary T cells, 2C TCR on hybridomas, purified N15 TCRαβ, P14 TCR on primary T cells, or TRBV TCRs (B13.C1/B17.C1 and B17.R1/B17.R2) expressed on hybridomas interacting with their corresponding pMHCs are plotted according to the ranked-order of peptide potencies (*bottom*). **f** best-fit model parameters *θ* (the tilted angle of the bonding interface, 1^st^ row), 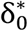 (the width of zero-force free-energy well, 2^nd^ row), and *n*^*^ (the number of unfolded amino acids, 3^rd^ row) are plotted *vs* the logarithm of the reciprocal peptide concentration required to stimulate half-maximal T cell proliferation (1/EC_50_) and fitted by a linear function. All error bars are fitting error (Supplementary Table 3).

#### Characterization of force-lifetime relationship

Previous work reported qualitative correlations between the TCR bond type, *i*.*e*., catch-slip bond *vs* slip-only bond, with the biological activity of the peptide to induce T cell activation, *i*.*e*., pMHC potency^4,7-10^. To reduce data representation and extract more information quantitatively from the bond lifetime *vs* force data, we defined several metrics from their model fit for each TCR–pMHC system and examined their correlation with T cell activation induced by a given interaction, using the OT1 system as an example because the quantitative ligand potency data are available^4,34^. We measured the peak bond lifetime, *t*_peak_, and the change, Δ*t*, from *t*_peak_ to the force-free bond lifetime, *t*_0_ = 1/*k*_0_ (Fig. 3a, *left panel*). We found the relative metric Δ*t* to be more suitable for comparison across different TCR systems, and to better correlate with ligand potency, than the absolute counterpart *t*_peak_ (Fig. 3c). Although the force where catch-slip bond lifetime peaks, *F*_opt_, occurs in a narrow range (10-20 pN), the force increment, Δ*F*, from *F*_opt_ to the level where bond lifetime returns to *t*_0_ defines the force span of a catch-slip bond (Fig. 3a, *left panel*). Both scaled parameters, *L* = Δ*t*/*t*_peak_ (relative length of lifetime) (Fig. 3d, *left panel*), and to a lesser extent *B* = Δ*F*/*F*_opt_ (relative breadth of lifetime) (Supplementary Fig. 2a), correlate with ligand potency well. We define a scaled parameter, *I* = *L*/(1 + *B*), which is the area ratio of two rectangles: Δ*t* × *F*_opt_ over *t*_peak_ × *F*_opt_ + Δ*F*. Remarkably, this combined parameter, which we term the catch bond intensity or catchiness, correlates best with the ligand potency across different TCR systems (Fig. 3d, *right panel* and 3e, 1^st^ *row*), supporting its usefulness as a metric of reduced data representation for a bond profile.

#### Model parameters’ correlation to ligand potency

It seems reasonable to test the validity of our model by examining the possible correlation of (or the lack thereof) the model parameters with features of the biological system, *e*.*g*., the ligand potency. The rationale is that if its parameters are capable of capturing and predicting such biological features, then the model would be more meaningful and useful than merely a curve fitting tool. Therefore, we plotted the tilted angle of the bonding interface *θ*, the number of the unfolded amino acids *n*^*^, and the width of the zero-force free-energy well 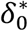 that best-fit the force-lifetime curves of OT1, 2C, P14, N15, and TRBV TCRs interacting with their corresponding panels of pMHCs (Fig. 3e). Gratifyingly, we observed good correlation between each model parameter and the peptide potency for all 21 published datasets of TCR–pMHC-I catch-slip bonds and slip-only bonds measured by four independent laboratories in five papers^4,6,9,16,29^. Moreover, model parameter and the peptide potency for OT1 quantitatively showed positive correlation with linear fitting (Fig. 3f).

In a previous study, we mutated residues in the 2C or 1G4 TCR and/or their corresponding pMHCs to alter bond profiles as predicted by SMD simulations, which was confirmed by BFP experiment^9^. We therefore fitted our model to the force-lifetime curves of these mutant TCR–pMHC interactions to evaluate the model parameters, 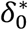, *θ*, and *n*^*^ (Fig. 4a, Supplementary Fig. 2b-c). In the absence of other functional data, we took an indirect approach to examine their correlations with the catchiness *I* of these bond lifetime *vs* force curve (Fig. 4b) since *I* and all three model parameters correlate with the peptide potency (Fig. 3e and f). Results are exemplified by the 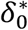, *θ*, and *n*^*^ *vs I* plots, which are graphed together with the data without TCR and MHC mutations that already showed functional correlates. For the WT OT1, 2C, P14, N15, and TRBV TCRs interacting with their corresponding panels of pMHCs, the best-fit model parameters 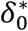 (Fig. 4c), *θ* (Supplementary Fig. 2d), and *n*^*^ (Supplementary Fig. 2e), correlate with the peptide potency predictor *I* (blue-open symbols). Remarkably, for the 2C and 1G4 TCRs specifically mutated to alter bond profiles with the corresponding WT or MT MHCs presenting the same agonist peptide, their best-fit 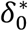, and to a lesser extent, *θ* and *n*^*^, also correlate well with *I* (Fig. 4c and Supplementary Fig. 2d-e, green*-*closed symbols). Interestingly, 1/*k*_0_ shows no correlation with *I* (Supplementary Fig. 2*f*), consistent with reports that zero-force bond lifetime does not correspond to ligand potency in these cases^4,9,34^.

**Figure 4.**
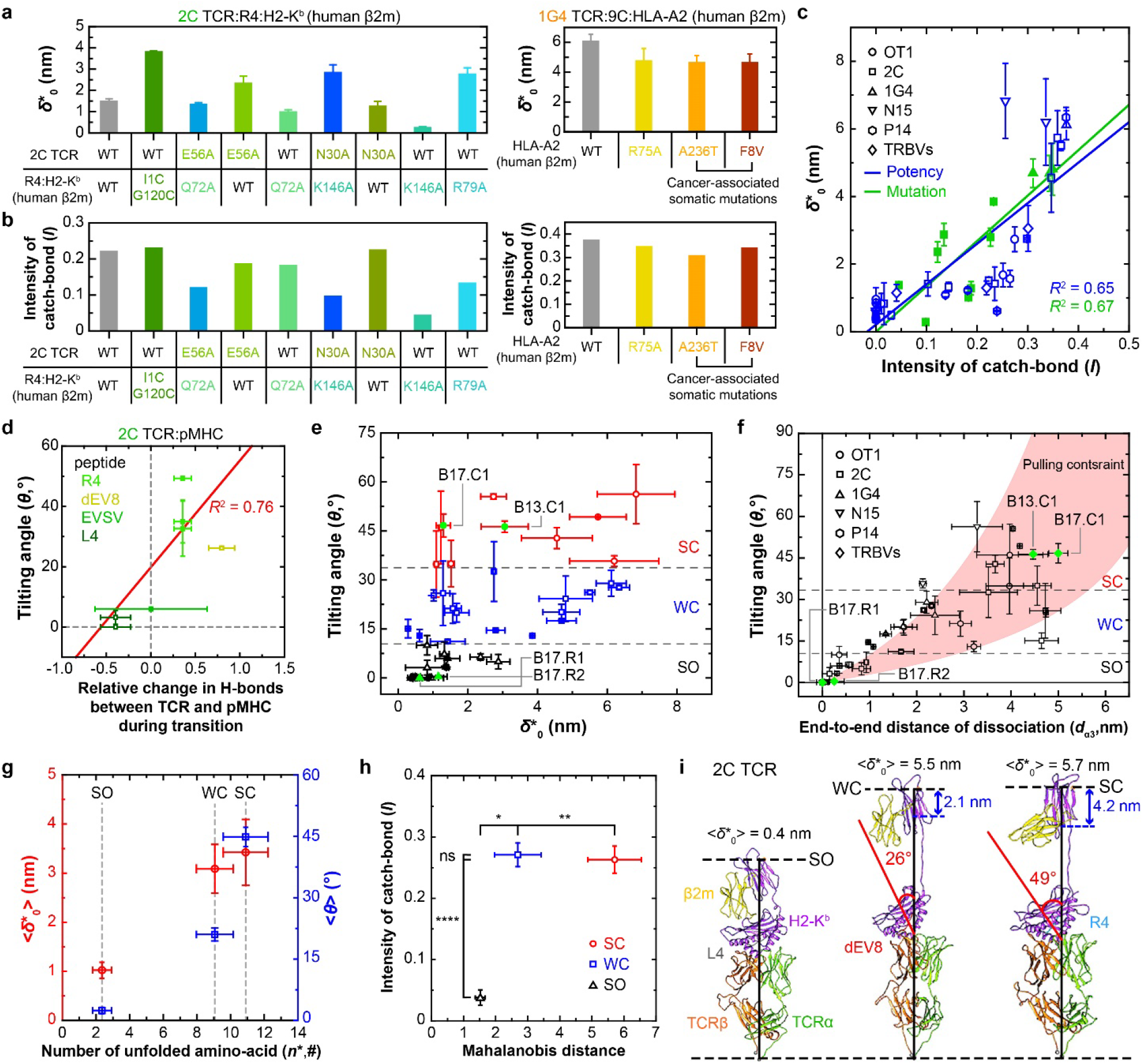
Properties and biological relevance of class I model parameters. **a**-**b** The width of zero-force energy well 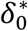 (**a**) and the intensity of catch bond *I* (**b**) calculated from WT or mutant 2C TCRs (*left*) and WT 1G4 TCR (*right*) interacting with their corresponding WT or MT pMHCs. The MT 2C TCRs and H2-K^b^s were designed to destabilize the TCR–pMHC interaction. The MT p:HLA-A2s were designed to either destabilize the TCR–pMHC interaction (R75A) or stabilize the MHC intramolecular interaction (A236T and F8V). **c** Data from **Fig. 3e** (3^rd^ row) are re-graphed as 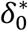 *vs I* plot to show their correlation (*blue*). Additional 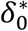 *vs I* data from MT TCRs and/or MT pMHCs without functional data also show strong correlation (*green*). Different TCR systems are indicated by different symbols. The two datasets were separately fitted by two straight lines with the goodness-of-fit indicated by *R*^2^. **d** Tilting angle of the bonding interface (*θ*) *vs* normalized net gain of hydrogen bonds at the interface between 2C TCR and the indicated pMHCs is plotted (*points*) and fitted (*line*) (error bars in *x*- and *y*-axes represent uncertainties propagated from errors in Supplementary Fig. 3 and fitting errors, respectively). **e** Clustering analysis reveals three clusters in the 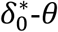 phase diagram: slip-only (SO, *black*), weak catch-slip (WC, *blue*), and strong catch-slip (SC, *red*) bonds. **f** Tilting angle (*θ*) *vs* end-to-end distance of dissociated α3 domain (*d*_*α*3_). The three types of bonds, SO, WC, and SC, are also clustered in this phase diagram, which are separated by the dotted lines that predicted from the pulling constraints of the model. The two pairs of TRBV TCRs are indicated in e and f by green dots. **g** The average molecular extensions at zero force (⟨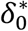⟩, *left ordinate*) and the average rotation angle (⟨*θ*⟩, *right ordinate*) (mean ± sem) are plotted *vs* the total number of unfolded amino acids (*n*^*^, *abscissa*) to show three clusters. Each bond type is indicated by a dotted line (*n* = 10, 16, and 17 for numbers of data in the SO, WC, and SC groups, respectively; individual data of each cluster are shown in (**e**-**f**)). **h** Catch bond intensity *vs* Mahalanobis distance plot, again showing three clusters. Principal component analysis was used to find principal axes. Mahalanobis distances for each cluster were calculated using common principal axes from total dataset (numbers of data are the same as (**g**)). **i** Structural models illustrating the conformations of three bond types according to their model parameters based on the previous SMD simulation of the 2C TCR system^9^. Two structural parameters (*θ, red*; *d*_*α*3_, *blue*) are indicated to show the differences between bond types. Unless otherwise described, all errors shown in (**a-f**) are fitting errors (Supplementary Table 3).

It is worth pointing out that the above results not only support our model’s validity but they also suggest that our model is more than a mere analytical framework to organize experimental data. Rather, the model parameters may be used to distinguish antigen recognition efficacy with force-amplified discriminative power. For example, the correlations of *θ* and 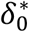 with peptide potency (Figs. 3f and 4c) indicate that the more potent the peptide, the higher the force sensitivity of its TCR–pMHC interaction, and the narrower the force range over which the TCR–pMHC interaction is sensitive to force (Fig. 2g).

#### Comparison between coarse-grained and all-atom models

Bonding interface tilting has been observed to be associated with changes in the number of hydrogen bonds bridging the TCR and pMHC molecules as they were pulled to unbind in SMD simulations^9^. Therefore, we investigated whether, and if so, how well the tilting angle would correlate with the change of hydrogen bonds between TCR and pMHC. Remarkably, *θ* was found to be proportional to the net change in the total number of hydrogen bonds at the bonding interface (Fig. 4d and Supplementary Fig. 3). This finding is intuitive and supports the validity of our coarse-grained model because it is able to recapitulate the results of all-atom SMD simulations^9^.

#### Classification of bond types by clustering analysis on phase diagrams

In Fig. 2e-g we have explored the model parameter space to identify regions that correspond to slip-only bonds and catch-slip bonds. Here we examined whether, and if so, how parameters that best-fit different experimental bond types map onto different regions of the parameter space. Since the model has four parameters, *θ*, 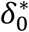, *d*_*α*3_, and *n*^*^ (*k*_0_ is not considered because if its lack of correlation with catch bond intensity), we analyzed their clustering and projected their values in the 4D parameter space onto three phase diagrams spanning the 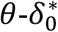 (Fig. 4e), *θ*-*d*_*α*3_ (Figs. 4f and Supplementary Fig. 4), and 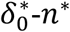 (Fig. 4g) 2D space. Clustering analysis of the model parameters that best-fit 43 TCR–pMHC bond lifetime *vs* force curves (Supplementary Fig. 5) shows three distinct clusters in the 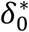 *vs θ* and *θ vs d*_*α*3_ plots as well as 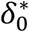 and *θ vs n*^*^ plots (Fig. 4e-g), which classify the TCR–pMHC interactions into slip-only (SO), weak catch-slip (WC) and strong catch-slip (SC) bonds, which correspond to weak, intermediate, and strong potencies for pathogenic peptides and their variants. Whereas transition in bond type from SO to WC and SC requires monotonical increase in *θ* and *n*^*^ (Fig. 4f, g), the corresponding change in 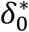 is non-monotonic (Fig. 4e-g). SO bonds show small *n*^*^, 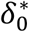, and *θ* values. WC and SC bonds observed from experiments are best-fitted by similar *n*^*^ (9 for WC and 11 for SC) but oppositely ranked 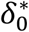 and *θ* values. To change from WC to SC bonds requires a slight increase in 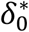 (from 3 to 3.5 nm) and a large increase in *θ* (from 20 to 45 °) (Fig. 4g). We also performed principal component analysis and calculated the Mahalanobis distances of the principal axes for the three bond types^35^, which are statistically separated in the catch bond intensity *vs* Mahalanobis distance plot (Fig. 4h). Interestingly, WC and SC bonds show distinct conformational changes despite their similar *I* values measured from the force-lifetime curves. The corresponding structural features of these three types of bonds are depicted in Fig 4i, which have been observed in our previous SMD studies^9^. Of note, model parameters visualized by SMD simulations are usually larger than their best-fit values, which may have two explanations: First, to enable dissociation to be observed in affordable computational times, much higher forces were used in simulations than experiments to accelerate the biophysical processes, which likely induced much larger conformational changes. Second, our model describes the average conformational change during the entire dissociation process, which is smaller than the maximum conformational changes likely to occur right before unbinding and to be captured by SMD.

#### Our structure-based model is superior to the generic two-pathway model

It seems that other published catch bond models should also be able to fit the experimental force-lifetime profiles analyzed here, given their relatively simple shapes. As an example, we examined the two-pathway model below^27^:

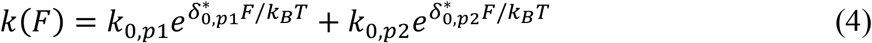

where *k*_0,*p*1_ and *k*_0,*p*2_ are the respective zero-force off-rates of the first and second pathway, 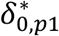 and 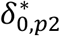 are the respective distances from the bound state to the transition states along the first and second pathways (Fig. 1a).Here the off-rate for each pathway takes the form of the Bell model, but the catch pathway parameter 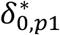 has a negative value^27^ (Supplementary Table 4). This model is generic as it has previously been applied to TCR–pMHC catch-slip bonds without considering the specific conformational changes^6^.

As expected, Eq. 4 also fitted our experimental data with goodness-of-fit measures statistically indistinguishable to Eq. 2 (Supplementary Figs. 1 and 10a-b). However, the fitting parameters correlate with neither the TCR/pMHC-I potency to induce T cell function nor the catch bond intensity (Supplementary Fig. 6, see the negative or zero correlation and the poor *R*^2^ values), hence have no biological relevance. This comparison indicates that the model developed herein is superior to the previous two-pathway model.

### Model for TCR catch bonds with class II pMHC

MHC class II differs from class I in three main aspects (comparing Fig. 2a and Fig. 5a): (1) MHC-I has three α domains and a β_2m_ domain whereas MHC-II has two α and two β domains. (2) MHC-I anchors to the T cell membrane through a single linker to the α_3_ domain. The β_2m_ domain attaches to the α_3_ domain instead of anchoring to the T cell membrane directly. By comparison, MHC-II anchors to the membrane through two linkers, one to the α_2_ domain and the other to the β_2_ domain. (3) The peptide is presented by the α_1_-α_2_ domains of MHC-I but the α_1_-β_1_ domains of MHC-II. These structural differences alter how forces are supported by and transmitted through, and induce conformational changes in, the TCR complexes with pMHC-I *vs* pMHC-II. Thus, it is necessary to modify the previous model in order for it to describe TCR catch and slip bonds with pMHC-II, which is done by using a different *δ*_1_*γ*(*F*) expression than Eq. 3 (Supplementary Model Derivations, Section B). This modification assumes force-induced partial unfolding and stretching of the TCR Vα-Cα joint and the MHC α_1_-α_2_ and β_1_-β_2_ joints during dissociation, which results in tilting of the bonding interface (Fig. 5a, b).

**Figure 5.**
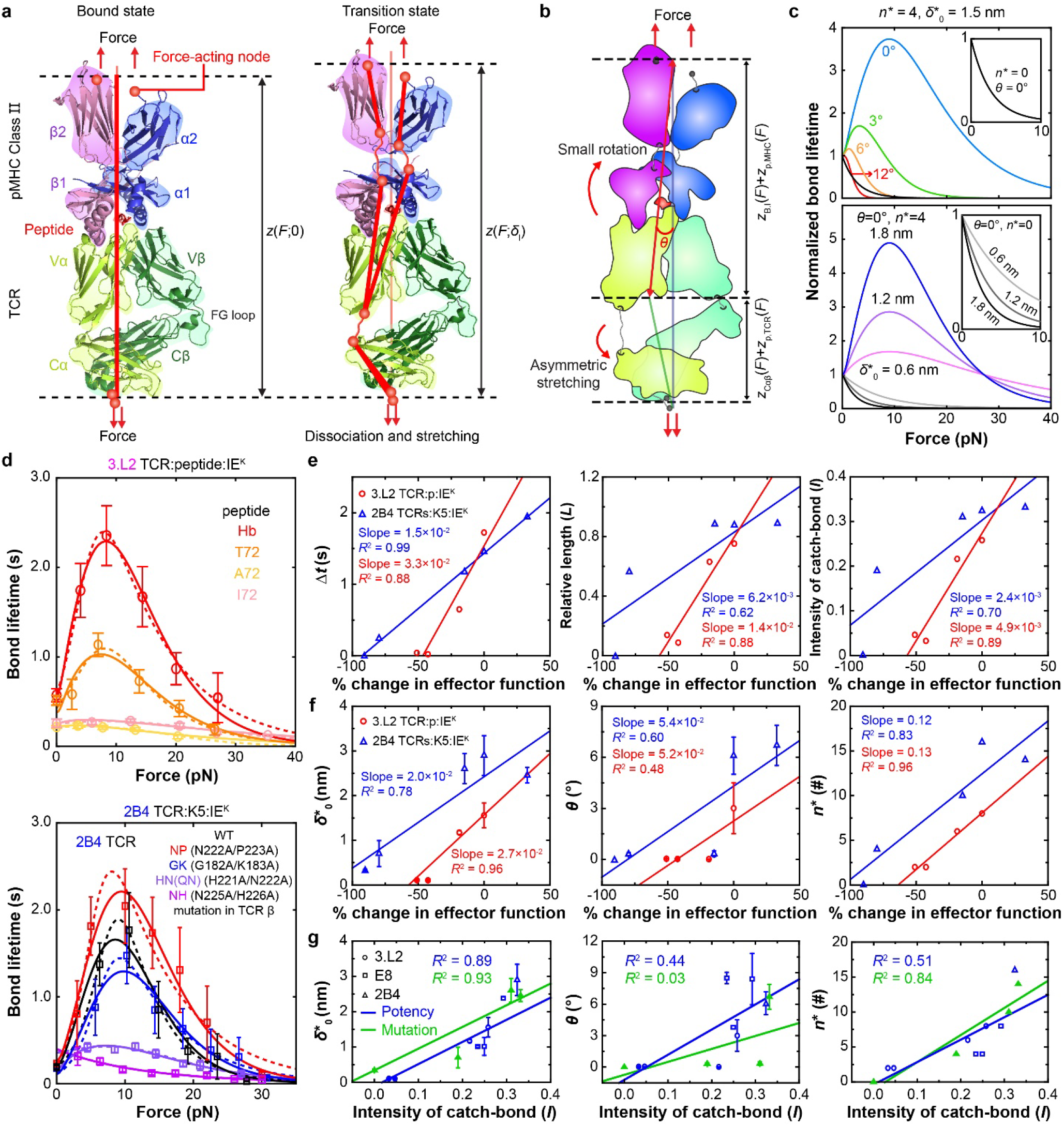
The TCR–pMHC-II model. **a** Force-induced conformational changes of a TCR– pMHC-II complex as it traverses from the bound state (*left*) to the transition state (*right*). The diagrams are based on the published co-crystal structure (2IAM) of the E8 TCR α (*yellow*) β (*green*) subunits and the TPI peptide (*red*) bound to the HLA-DR1 α (*blue*) β (*pink*) subunits with various domains indicated. The force-transmission paths are shown as red lines connecting the force-acting nodes. **b** Various contributions to the total extension projected on the force axis: stretching of the TCR Cα and Cβ domains (*z*_Cαβ_), asymmetric partial unfolding of the TCR Vα-Cα and Vβ-Cβ interdomain hinges (*z*_p,TCR_), asymmetric partial unfolding of the MHC α1-α2 and β1-β2 interdomain hinges (*z*_p,MHC_), and rotation between the α1-β1 and α2-β2 domain hinges and tilting of the bonding interface between the MHC α1-β1 and the TCR Vα-Vβ by an angle *θ* (*z*_B.I_). **c** Theoretical normalized bond lifetime *vs* force curves. The effects of changing *θ* and 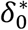 are shown in the upper and lower panels, respectively, for the indicated parameter values. **d** Fitting of predicted 1/*k*(*F*) curves (*dashed lines* by two-pathway model and *solid lines* by TCR–pMHC-II model) to experimental bond lifetime *vs* force data (*points*, mean ± sem) of 3.L2 TCR on CD4^-^CD8^+^ naive T cells interacting with indicated p:I-E^k^’s^5^ (*upper*) or WT and indicated MT 2B4 TCRs on hybridomas interacting with K5:I-E^k^. **e** Dimensional metrics, Δ*t* (*left*), scaled relative length of bond lifetime *L* (*middle*), and intensity of catch bond *I* (*right*) *vs* reciprocal % change (relative to WT) of effector function, i.e., the peptide dose required for 3.L2 T cells to generate 40% B cell apoptosis (1/EC_40_)^36^ (*red*) or the area under the dose response curve (AUC) of the 2B4 hybridoma IL-2 production^37^ (*blue*) plots. **f** Best-fit model parameters 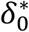 (*left*), *θ* (*middle*), and *n*^*^ (*right*) are plotted *vs* reciprocal relative % change of effector function. **g** The three model parameters in f for both the 3.L2 and 2B4 TCR systems are plotted *vs* the catch-bond intensity *I* and fitted by a straight line. We also added to each panel an additional point obtained from data and model fit of E8 TCR–TPI:HLA-DR1 interactions^13^. All error bars in **e-g** are fitting error (Supplementary Table 6).

In the class II model, the same parameters 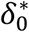, *n*^*^, and *θ* are used but the MHC contribution to *n*^*^, *i. e*., *n*_p,MHC_, represents the average number of amino acids in the polypeptides of the partially unfolded MHC-II α_1_-α_2_ and β_1_-β_2_ joints instead of the MHC-I α_1_α_2_–α_3_ joint, and the relationships between *θ* to other structural parameters are also different from the class I model (Fig. 5b and Supplementary Model Derivation, Section B). Like the class I model, the *k*_0_/*k vs F* plots for a range of *n*^*^, *θ*, and 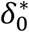 in Fig. 5c show similar features to Fig. 2e and meet our objective of being capable of describing catch-slip bonds if and only if *n*^*^ > 0 and *θ* ≥ 0. Unlike the class I model, a much smaller *θ* value (< 10°) is seen in the class II model (compared Fig. 5c and Supplementary Fig. 7 with Fig. 2e-g), indicating the main conformational change responsible for TCR–pMHC-II catch-slip bond is unfolding rather than tilting. The validity of this model is supported by its excellent fitting to our 6 published datasets of mouse 3.L2 (Fig. 5d, upper panel)^5^ and human E8 (Supplementary Fig. 9i)^13^ TCRs.

In addition, we generated 5 new datasets in this work specifically designed to test our model prediction that (de)stabilizing the TCR-CD3 would alter the TCR–pMHC bond profile (Table 6). Of these, the WT represents a hybrid 2B4 TCR with its mouse Vαβ fused with the Cαβ of the human LC13, expressed on hybridoma cells with human CD3 (see Fig. 7f below) and the four double mutants each replaces two Cβ residues by Ala to respectively decrease (NP) or increase (GK, HN, and NH) Cβ–CD3 interactions under force (see Fig. 7f-i below). Remarkably, interactions of the same K5:I-E^k^ with these five TCRs indeed yielded different bond profiles that were well fitted by our class II model (Fig. 5d, lower panel).

Furthermore, the four metrics of both the 3.L2 and 2B4 TCR–pMHC-II bond lifetime *vs* force curves Δ*t, L, I*, and *t*_peak_ correlate well with the published peptide (for 3.L2) and TCR (for 2B4) potency^36,37^ (Fig. 5e and Supplementary Fig. 8). Moreover, the three model parameters *θ, n*^*^, and 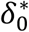 also correlate well with the TCR potency for the 2B4 system^37^ and with the ligand potency for the 3.L2 system^5,36^ (Fig. 5f, see the goodness of fitting, *R*^2^), supporting the ability of the metrics of the bond profile and the model parameters to recognize the change in the TCR-CD3 ECD interaction in addition to the ability to discriminate antigen. These properties are desirable, intuitive, and are consistent with the parallel properties found in the class I model. Similar to the class I model parameters, 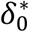 correlates well with the catch bond intensity for the pooled results from all class II data (Fig. 5g, *left*), but *θ* and *n*^*^ correlate less well with *I* (Fig. 5g, *middle and right*). Thus, the validity of the class II model is further supported by the faithful mapping of the relationship between biophysical measurements of catch and slip bonds and biological activities of the TCR–pMHC-II interactions onto a relationship between model parameters and biological function.

As expected, Eq. 4 also fitted our experimental data with goodness-of-fit measures statistically indistinguishable to Eq. 2 (Fig. 5d and Supplementary Fig. 10c-d). Similar to the class I system, the fitting parameters correlate with neither the TCR/pMHC-II potency to induce T cell function nor the catch bond intensity (Supplementary Fig. 6, see the negative or zero correlation and the poor *R*^2^ values), hence have no biological relevance. This comparison again indicates that the model developed herein is superior to the previous two-pathway model.

### Cross-examination of class I model against class II data and *vice versa*

Upon examining the catch-slip and slip-only bond lifetime *vs* force curves in Figs. 3a, 3b, 5d and Supplementary Fig. 1, it became apparent that the data seem very similar regardless of whether they are for class I or class II pMHC. Indeed, applying the class I model to the class II data and *vice versa* reveals that both models are capable of fitting both data well (Supplementary Fig. 9) and produce statistically indistinguishable goodness-of-fit measures (Supplementary Fig. 10). This is not surprising because both models have five fitting parameters and the bond lifetime *vs* force curves have relatively simple shapes. Nevertheless, fitting the same data by different models returns different parameter values depending on the model used, because the two models are constructed based on different structures and force-induced conformational changes of the TCR–pMHC complexes. Therefore, we asked whether the best-fit model parameters were capable of distinguishing data from the two classes of pMHCs and of telling whether a correct model was used to analyze data of matched MHC class. To answer these questions, we plotted 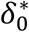 *vs I* (Fig. 6a, b) and 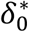 *vs n*^*^ (Fig. 6d, e) using values of the two models that best-fit the data of OT1, 2C, 1G4, P14, N15, and TRBV TCRs interacting with their respective panels pMHC-I ligands (Fig. 6a, d) as well as 3.L2, WT and MT 2B4, and E8 TCRs interacting with their respective panels of pMHC-II ligands (Fig. 6b, e). Surprisingly, the dependency of 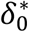 on *I* is 2-5-fold stronger (i.e., steeper slope) (Fig. 6c), indicating a greater discriminative power of receptor/ligand potency, for the matched than the mismatched cases. Furthermore, it is well-known that the average contour length per a single amino acid *l*_c_ is ∼ 0.4 nm^22,38,39^, which sets the biophysical limit for the slope of 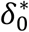 *vs n*^*^ plots. Indeed, we found that the slopes of the 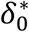 *vs n*^*^ plots are within this limit for both model fits of both class I and class II data (Fig. 6f). Moreover, the goodness-of-fit (*R*^2^) values of the linear fit to the 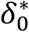 *vs I* (Fig. 6c) and 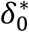 *vs n*^*^ (Fig. 6f) data are much greater for the matched than the mismatched cases, indicating more appropriate models for the data in the matched than the mismatched cases. Indeed, the *R*^2^ value for fitting the class II data by the class I model is too small to be statistically reasonable, therefore telling the mismatch between the model and the data. These results indicate that the model parameters are capable of distinguishing data from the two classes of pMHCs.

**Figure 6.**
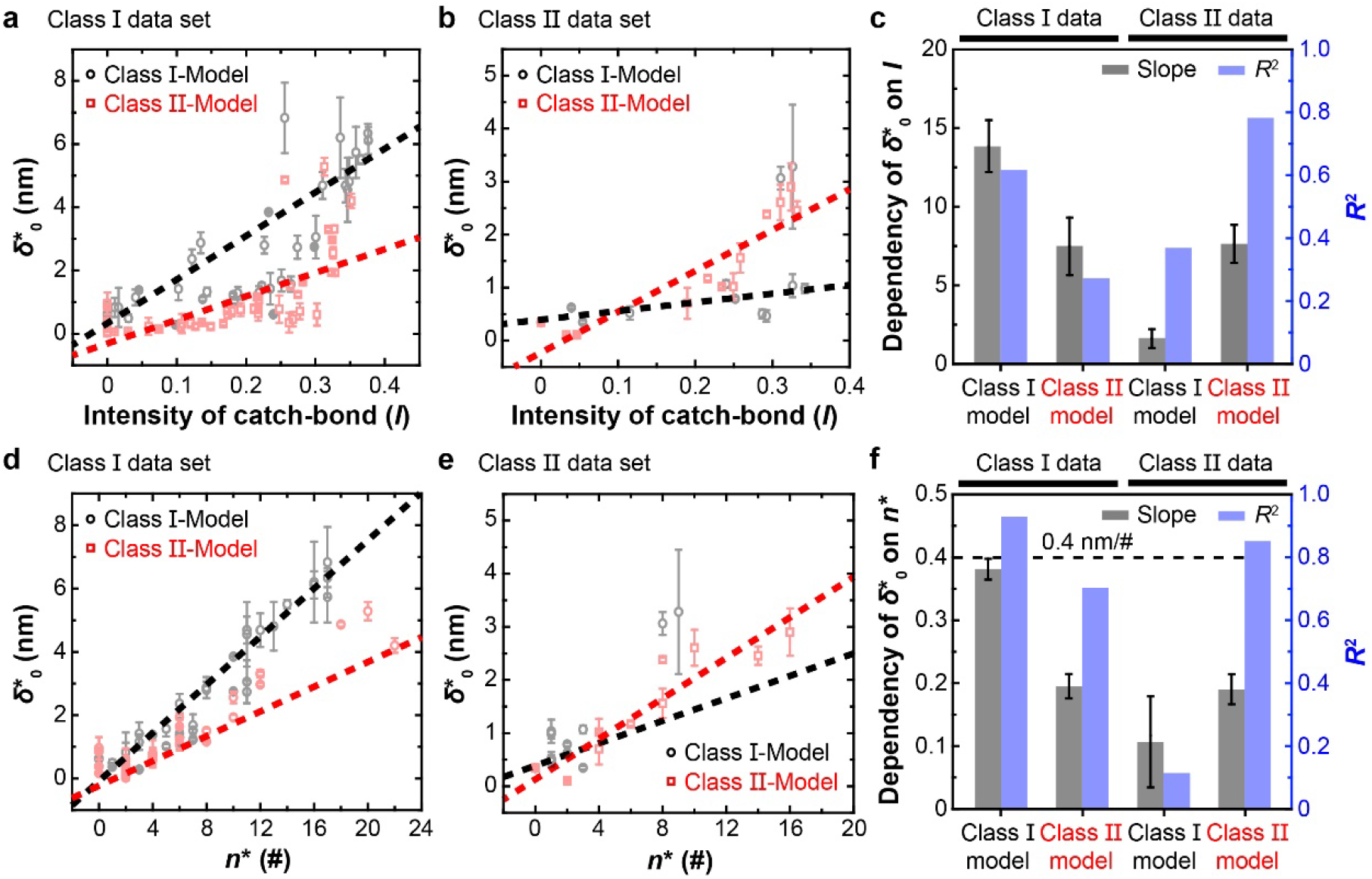
Cross-examination of class I and II models against class I and II data. **a**-**b**, 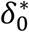 *vs I* plots obtained using class I (*black*) or class II (*red*) model to fit force-lifetime data of TCR interacting with pMHC-I (**a**) or pMHC-II (**b**) molecules. In each panel, two sets of parameter values were returned from fitting depending on whether class I (*black*) or class II (*red*) model was used because they are based on different structures and conformational changes of the TCR–pMHC complexes. **c** The slopes (*gray*, the level of correlation between 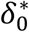 and *I*) and goodness-of-fit (*R*^2^) (*blue*, the degree of appropriateness of the model for the data) of the linear fit in a and b are shown in the matched (1^st^ and 4^th^ groups) and mismatched (2^nd^ and 3^rd^ groups) cases. **d**-**e** 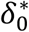 *vs n*^*^ plots obtained using class I (*black*) or class II (*red*) model to fit force-lifetime data of TCR interacting with pMHC-I (**d**) or pMHC-II (**e**) molecules. **f**, The slopes and goodness-of-fit of the linear fit in d and e are shown in the matched (1^st^ and 4^th^ groups) and mismatched (2^nd^ and 3^rd^ groups) cases. The slopes indicate the average unfolding extension per amino acid (nm/a.a.) from each model, which are compared to the maximum average contour length per amino acid of ∼0.4 nm/a.a. (biophysical limit, black dashed line with considerable deviation)^22,38,39^. All error bars are fitting error (Supplementary Table 6).

### Model validation by mutagenesis to alter force-induced conformational changes

The published datasets re-analyzed by our models include TCR interactions with altered peptide ligands, yielding different catch and slip bonds whose profile metrices and model parameters correlate with varied peptide potencies to induce T cell activation (Figs. 3, 5d upper panel, 5e-f red, and 5g blue; Supplementary Figs. 1a, 1b panels 1-2, 1c, 1d, 1e panel 1, 1f-h, 8 red, and 9i). They also include mutations on the TCR or MHC specifically designed to assess how structural change altered bond profile (Supplementary Figs. 1b panels 3-10, and 1e panels 2-4) but do not include functional data^9^. We thus performed two sets of new studies to further validate the class I and II models, respectively, using mutations located away from the TCR and pMHC binding interface but capable of impacting their respective conformational changes under force, which were analyzed by MD simulations, bond lifetime measurements, and functional assays.

The first set of studies compared the WT and a hybrid H2-K^b^α3A2 that swaps the mouse β_2_m with the human β_2_m because the latter binds the mouse class I heavy chain with a higher affinity and better support peptide binding than the former^40^. Since it is easier to make soluble hybrid than complete mouse H2-K^b^ protein, many of our previous studies used the hybrid H2-K^b^ (Supplementary Table 3). Surprisingly, T cells kill less efficiently target cells expressing the hybrid H2-K^b^ than the WT molecule^41^. Our previous study using double-cysteine mutations to lock the α_1_α_2_–β_2_m connection by disulfate bond suppressed both pMHC conformational changes and its catch bond with TCR concurrently^9^ (Supplementary Fig. 1b, compared the R4 curves in panels 2 and 3). Using SMD simulations, we observed force-induced dissociation of the α_1_α_2_–β_2_m interdomain bond (Supplementary Movie 1). We compared MD simulated interactions of H2-K^b^ α chain with mouse β_2_m (using the crystal structure 1G6R) and human β_2_m (using a model built based on 1G6R and 2BNR), finding that Arg14, Glu232, and Gly237 of the H2-K^b^ α chain respectively interacted with three residues – Asp34, Lys6, and Tyr67 – of the human β_2_m but not the corresponding residues of the mouse β_2_m (Fig. 7a-c). This indicates that the hybrid H2-K^b^ has a more stable structure and hence less able to respond to force induction of conformational change than the WT molecule, predicting a less pronounced TCR catch bond with the same peptide presented by the hybrid than the WT H2-K^b^. Remarkably, the newly measured force-dependent bond lifetime indeed showed a much more pronounced catch bond of the 2C TCR with R4 peptide bound to WT than hybrid H2-K^b^ (Fig. 3b, 1^st^ panel), supporting the prediction of our class I model. Consistent with previous report^41^, functional assay also showed that the WT H2-K^b^ was more able to activate T cells than hybrid H2-K^b^ (Fig. 7d-e), further validating the class I model.

**Figure 7.**
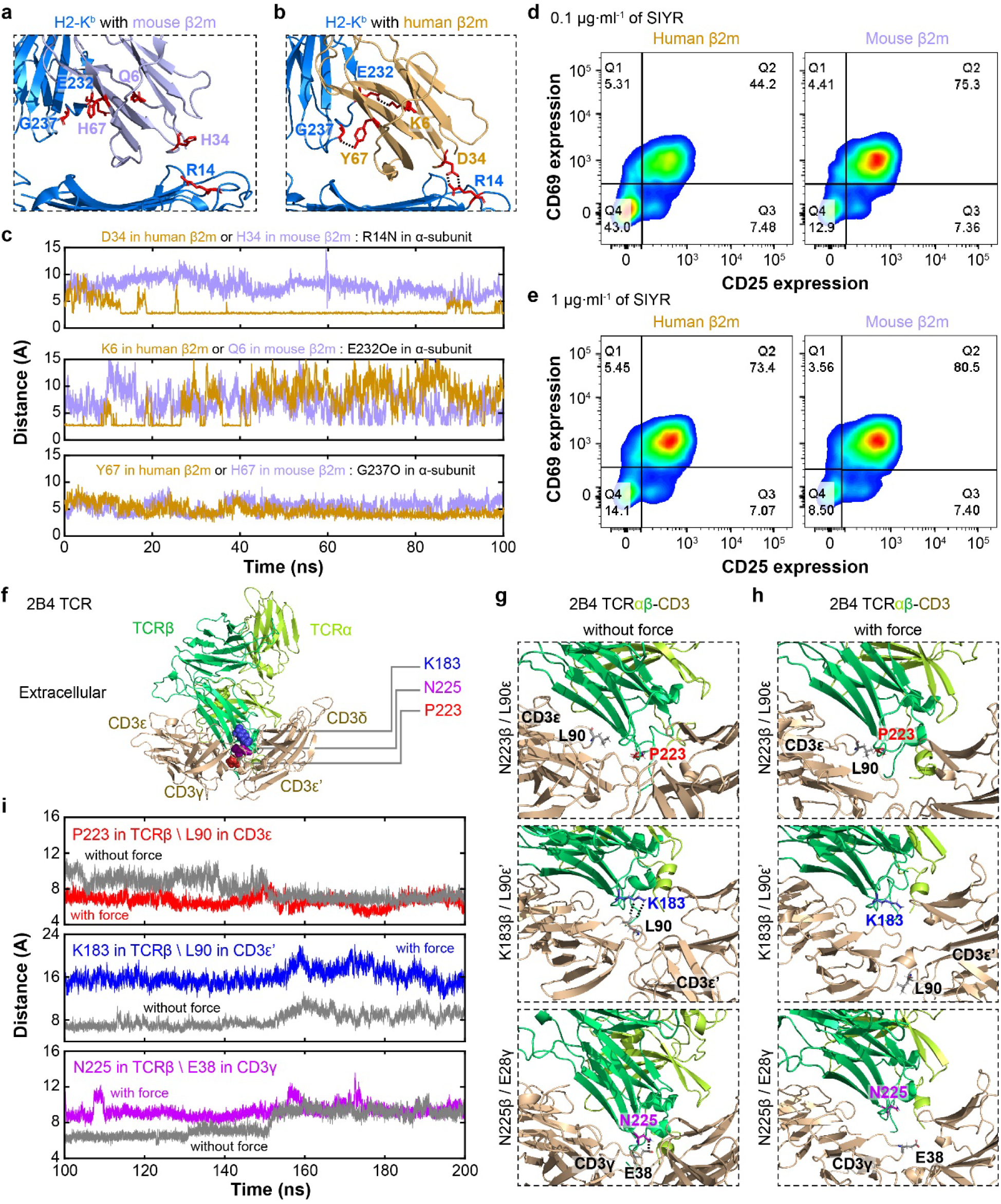
Model validation by mutagenesis. **a**-**c** Comparison of structures (**a, b**) or noncovalent contacts (**c**) of interactions of the H2-K^b^ α_1_α_2_ (*blue*) with mouse (**a, c**) and human (**b, c**) β_2_m (*purple* for mouse and orange for human, respectively). The structures in **a** and **b** are depicted by ribbon diagram using snapshots from SMD stimulations (initials modelled based on 1G6R and 2BNR) with side-chains of the interacting residues shown by sticks (*red*). Simulated time-courses of distances between the interacting H2-K^b^ α chain residues and β_2_m residues are plotted in **c**, showing shorter distances with the human β_2_m and longer distances with the mouse β_2_m. **d**-**e** Comparison of potencies to activate naïve CD8^+^ 2C T cells by hybrid (*left* column) and WT (*right* column) R4:H2-K^b^ at 0.1 μg/ml (**d**) or 1 μg/ml (**e**) concentration for 72 hours. T cell activation was assayed by flow cytometric analysis of upregulation of surface markers CD69 (*y*-axis) and CD25 (*x*-axis) using PE-conjugated anti-CD69 and PE-cy7-conjugated anti-CD25 antibodies. **f** Structure of 2B4 TCRαβ showing the locations of residues N222-P223, G182-K183, and N225-H226 on Cβ domain with CD3 complex (6JXR). **g-h** Comparison of interactions of P223 (1^st^ row), K183 (2^nd^ row), and N225 (3^rd^ row) with the corresponding CD3 residues in the absence (**g**) and presence (**h**) of force using MD simulations (initial build upon 6JXR).. **i** Simulated time-courses of distances between Cβ P223 and CD3ε L90 (1^st^ row), Cβ K183 and CD3ε’ L90 (2^nd^ row), as well as Cβ N225 and CD3γ E38 (3^rd^ row) in the absence (*gray*) and presence (*colored*) of force.

The second set of studies examined a hybrid TCR made of the mouse 2B4 Vαβ fused with the human LC13 Cαβ and 4 double mutations on the Cβ domain, which have been indicated by our previous NMR and chemical shift experiments^37^ and by recently published cryoEM structures^42,43^ to impact its interactions with human CD3 (Fig. 7f). We performed MD simulations to examine the Cβ–CD3 *cis*-interactions in the absence (Fig. 7g) and presence (Fig. 7h) of a force to mimic pulling on the Vαβ by the engaged K5:I-E^k^ (Supplementary Movie 2-4). We found that Cβ Pro223 is force-stabilizing (Fig. 7i, *top*) whereas Cβ Lys183 and Asn225 are force-destabilizing (Fig. 7i, *middle* and *bottom*). These results suggest that the double mutant N222A/P223A (NP) may result in less stable, whereas G182A/K183A (GK) and N225A/H226A (NH) may result in more stable, Cβ–CD3 *cis*-interactions under force, therefore potentially limiting force-induced conformational changes in the TCRαβ less (for NP) and more (for GK and NH), respectively, than the WT molecule. Interestingly, NP was identified as a gain-of-function mutation whereas GK and NN (plus another double mutant H221A/N222A, or HN) were identified as loss-of-function mutations by functional assays^37^. Supporting the prediction of our class II model, force-dependent bond lifetime measurements by BFP indeed showed a more pronounced catch-slip bond of the NP mutant, less pronounced catch-slip bonds of the GK and HN mutants, and slip-only bond of the NH mutant, compared to the WT 2B4 TCR interaction with K5:I-E^k^ (Fig. 5d, *lower panel*). Remarkably, the bond profile metrices (Fig. 5e and Supplementary Fig. 8, *blue*) and best-fit model parameters (Fig. 5f *blue* and 5g *green*) were found to correlate with T cell function, further validating the class II model.

## DISCUSSION

With the exception of a recent paper that observed T cell pulling on TCR by ∼2 pN forces using a spider silk peptide-based force probe^44^, five publications from two laboratories demonstrated that TCR experienced endogenous forces of 12-19 pN using DNA-based force probes^8,15,45-47^. Also, except for a single paper that failed to observe catch bonds in the 1G4 TCRαβ ECD using a flow chamber^48^, extensive data in 9 papers from four laboratories^4-6,8,9,13,16,28,29^ plus new data presented in this study have demonstrated catch bonds in 12 TCRs (including 1G4) using BFP and OT. These experiments prompted us to develop two mathematical models for TCR catch bonds following the 1D formulation of Guo et al.^22^, one with class I and the other with class II pMHC, based on Kramer’s kinetic theory and accounted for the 3D coarse-grained structures, molecular elasticity, and conformational changes of the TCR–pMHC-I/II complexes. Previously, several models have been developed to describe catch-slip bonds of protein–protein interactions, including selectins–ligands^27,49-51^, integrins– ligands^52,53^, platelet glycoprotein Ibα–von Willebrand factor^54^ cadherin–catenin^55^, vinculin– actin^56^, and talin–actin^57^ interactions. Except for the sliding-rebinding model, which is based on force-induced conformational changes in P-selectin–ligand observed from SMD simulations^51^, none of these models have included any specific structural considerations of the interacting molecules. Instead, these models are based on a generic physical picture of dissociation along two pathways from either one or two bound states in a 2D energy landscape that is tilted by force^21,58^. Except for the two-pathway model tested here, which has 4 parameters^27^, all other models have 5-10 parameters; therefore, over-fitting is a concern for applying them to some of the datasets analyzed here. Although the two-pathway model is capable of fitting the TCR–pMHC catch-slip bond data well, as shown previously^6^ and tested more extensively by much larger datasets here (Supplementary Fig. 1 and Fig. 5d), the four best-fit model parameters correlate with neither T cell function nor the catch bond intensity for either class I or class II system (Supplementary Fig. 6), hence informing no biological insights. Consequently, it cannot distinguish the class I and class II systems because the parameters of the generic two-pathway model have nothing to do with the differential structures of the class I and II systems. This comparison highlights the utility and usefulness of our models.

Force-induced conformational changes of TCR–pMHC-I complexes have been observed or suggested by single-molecule experiments and SMD simulations^6,9^. Parameterizing these conformational changes by the number of unfolded amino acids *n*^*^ and the bonding interface titling angle *θ* in the class I model allows us to explain mechanistically and quantitatively the TCR–pMHC-I catch-slip and slip-only bonds. Indeed, the criteria for catch-slip bond are *n*^*^ > 0 and *θ* > 0; the greater their values the more pronounced the catch bond. Importantly, the validity of the class I model has been supported by its capability to fit all force-lifetime datasets published to date plus one dataset presented here, and by the correlation between the best-fit model parameters and the available biological activity data induced by the TCR–pMHC-I interactions.

By comparison, the respective ranges of *n*^*^ and *θ* for the class II model are smaller, consistent with the sturdier structure of the pMHC-II molecule^24^. Mutagenesis studies and MD simulations in the present work have supported the hypothesis of force-induced conformational changes in the TCR structure. Our class II model has also been tested by all published datasets plus four datasets presented here, and their best-fit parameters also correlate well with the biological activities induced by the TCR–pMHC-II interactions. Furthermore, the validity of models of both classes has been supported by the findings that the best-fit model parameter 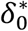 correlates with the catch bond intensity *I* and that the 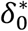 *vs I* and 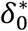 *vs n*^*^ plots have more appropriate slopes and *R*^2^ values when the model matches than mismatches the data.

We should note that since TCR–pMHC unbinding is assumed to be a spatially continuous and temporally instantaneous process, all structural parameters determined here represent mean field values, and they were evaluated by fitting the mean bond lifetime *vs* force data. However, individual bond dissociation events are inherently stochastic; and as such, need not be deterministically mapped onto any specific conformational changes in a one-on-one fashion. Instead, any particular bond type and parameter sets are related on the average sense. Future studies are required to extend the current framework to relate more detailed structural changes and bond lifetime distributions, *e*.*g*., to account for more sequential partial unfolding events prior to transition state as suggested by experiments^6,12,59^.

A strength of our agent-based models lies in their ability to incorporate many different ideas and knowledge into a simple 1D formulation. This simplicity facilitates model application to both class I and II experimental systems, enables quantitative interpretation of TCR–pMHC bond lifetime *vs* force profiles, expresses biological functions by biophysical measurements, and suggests structural mechanisms of how the TCR mechanotransduction machinery might work. However, the 1D simplification is also a weakness because theoretically these models can only describe single-step dissociation by entropic conformational fluctuation in the low-force regime from a single-state along a single-dissociation path, implicitly assuming that there is only a single energy barrier. Although some catch-slip and slip-only bonds can be described by such simple models^5^, more complicated TCR–pMHC bonds has been reported. These are evidenced by the multi-exponential bond lifetime distributions at constant forces, which have been fitted by data-driven multi-state, multi-pathway models^13^. To address this weakness, future studies may extend the present 1D model to 2D, *e*.*g*., by combining Eqs. 2 and 4, to enable proper description of multi-exponential bond survival probabilities.

We introduced the catch-bond intensity *I* as a dimensionless scaled metric for the bond lifetime *vs* force curve and generated four model parameters that describe the curve’s geometric features. Upon analyzing all 49 catch-slip and slip-only bond profiles published to date by four independent laboratories^4-6,8,9,13,16,28,29^ plus 6 new ones reported here, we found that these quantities do a better job to predict TCR function than any other quantities. This finding may explain how force amplifies TCR signaling and antigen discrimination, because *I* is defined by a force curve and *n*^*^ and *θ* only predict signaling when they assume none-zero values at *F* > 0. It should be noted that despite the comparable force ranges, highly variable lifetimes have been observed for different TCR systems interacting with different pMHCs (e.g., WT *vs* hybrid H2-K^b^). Even the same TCR–pMHC interactions could display different bond lifetimes in the absolute scale, depending on the cells on which the TCR is expressed. The power for the catch-bond intensity *I* to predict TCR signaling and discriminate antigen may lie in the ability of this dimensionless number to capture different bond lifetime patterns in a relative scale.

A recent study showed surprising features of reversed-polarity of TRBV TCRs such that interactions of NP_366_:H-2D^bD227K^ to TCRs B13.C1 and B17.C1 induced T cell signaling, whereas interactions of the same pMHC to B17.R1 and B17.R2 TCRs did not^16^. Despite that the former two TCRs formed catch-slip bonds with NP_366_:H-2D^bD227K^ and the latter two TCRs formed slip-only bonds, the authors suggested that the signaling capability of the B13.C1 and B17.C1 TCRs could not be attributed to their force-prolonged bond lifetimes because the B17.C1 TCR–H-2D^bD227K^ bond was shorter-lived than the B17.R2 TCR–NP_366_:H-2D^bD227K^ bond across the entire force range tested. Even at 9.4 pN, which was *F*_opt_ for the former with a *t*_peak_ = 0.61 s, the latter lived 2.48 s on average, and the longest lifetime of the latter was *t*_0_ = 2.83 s occurred at zero force^16^. The authors hypothesized that the TCR–pMHC docking orientation, which was “canonical’ for the B13.C1 and B17.C1 TCRs but “reversed” for the B17.R1 and B17.R2 TCRs, underlain the signaling outcomes by directing the position of Lck relative to the CD3. However, we suggest that even without knowing the docking orientation, our model parameters are capable of determining the signaling outcomes. Indeed, our analysis correctly maps the data of the B13.C1 and B17.C1 TCRs onto the high peptide potency region and the data of the B17.R1 and B17.R2 TCRs onto the low peptide potency region of the 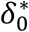 *vs I* (Fig. 4c), 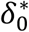 *vs θ* (Fig. 4e), and *θ vs d*_*α*3_ (Fig. 4f) phase diagrams. Thus, by mechanistically modeling the effect of force on bond dissociation, TCR signaling and antigen discrimination can be predicted by the model parameters.

The success in our model applications indicate that the conformational changes assumed in the models may be important to the TCR triggering, thereby suggesting testable hypotheses for future studies designed to investigate the inner workings of the TCR mechanotransduction machinery, *e*.*g*., to extend and/or revise models regarding how TCR signaling is triggered. Some TCR triggering conceptual models involve conformational changes and/or catch bond formation^1,2,60^. Our structure-based biophysical models relate catch and slip bonds to TCR–pMHC conformational changes. For the class I model, the parameterized structural changes include force-induced disruption of the MHC α_1_α_2_–β_2m_ interdomain bond, partial unfolding of the α_1_α_2_–α_3_ joint, tilting of the TCR–pMHC bonding interface, and partial unfolding of the Vα-Cα and Vβ-Cβ joints. For the class II model, these are primarily limited to the force-induced partial unfolding of the MHC-II α_1_-α_2_ and β_1_-β_2_ joints as well as the Vα-Cα and Vβ-Cβ joints. Besides these, one additional conformational change observed in the SMD simulations of TCRαβ–pMHC dissociation is unfolding of the connecting peptides between the TCRαβ ECD and transmembrane domain^9^. We chose not to include this conformational change in our models because such unfolding would likely be prevented by the interaction of the Cαβ with the CD3 subunits. Consistent with this assumption, the experimental data used for model fitting to evaluate conformational change parameters (*n*^*^ and *θ*) are those of pMHC bonds with TCR-CD3 complexes on the cell membrane that includes the TCRαβ ligand binding subunits and the CD3 signaling subunits (except for the N15 TCRαβ case which is soluble ECD only). Indeed, our previous work found that catch bonds of purified TCRαβ were altered from those of cell surface TCR interacting with the same pMHCs^13,28^, which is reflected by their changed model parameters (Supplementary Fig. 11). As such, the TCRαβ conformational changes predicted by our models provide a constraint for possible CD3 conformational changes in the TCR-CD3 complex to be considered in future TCR triggering models. Indeed, our new data on WT and MT 2B4 TCR–K5:I-E^k^ interactions indicate the importance of the TCRαβ–CD3 *cis*-interaction on catch bond formation of the TCR–pMHC *trans*-interaction.

Another constraint to be considered by future studies is that imposed by the coreceptor CD4 and CD8, as co-ligation of the coreceptor prolongs bond lifetimes, amplifies catch bonds, and may even changes slip-only bonds to catch-slip bonds^7,8,13,16,29^. Future studies should also consider how to extend the current models to pre-TCR catch and slip bonds with a broad range of ligands^59,61^. Instead of the TCRα, the pre-TCR uses the TCRβ chain to dimerize with a common pre-Tα chain, which lacks the variable domain (hence no Vα-Cα hinge). Without extension, even if our models are still able to fit the data of pre-TCR–ligand bonds or data of TCR–pMHC bonds where the TCRβ F-G loop was deleted or bound by an anti-TCR antibody^6^, the best-fit model parameters may not correspond to the conformational changes of these molecular complexes which are likely different from the conformational changes in the TCR– pMHC bonds with intact F-G loop.

An objective of the present work is to explore the extent to which 1D models can describe experimental data with a minimal set of meaningful parameters. Our parameters consider coarse-grained structural features and relate catch and slip bonds to specific force-induced conformational changes of the TCR–pMHC complex. This approach should be extendable to the modeling of other receptor–ligand systems of different structural features yet also form catch and slip bonds, such as selectins^18,62,63^, integrins^52,53,64-67^, cadherin^68^, Fcγ receptor^69^, notch receptor^70^, platelet glycoprotein Ibα^54,71^, FimH^72^, actin with myosin^73^, actin with actin^74,75^, cadherin-catenin complex with actin^55^, vinculin with actin^56^, talin with actin^57^, and microtubule with kinetochore particle^76^.

Our models allow us to develop working hypotheses regarding how T cell function is regulated through structural modulations of catch and slip bonds. For example, in this study we validated a prediction of the class I model that strengthening of the α_1_α_2_–β_2m_ interdomain bond would weaken the TCR–pMHC catch bond, which would in turn reduce T cell activation. This prediction has also been supported by our published data that somatic mutations in HLA-A2 found in some cancer patients impair TCR–pHLA-A2 catch bonds, which may explain the suppressed anti-tumor T cell immunity^9^. More interestingly, our models pave the way for engineering of TCR function for tumor immunotherapy by modulating the TCR catch and slip bonds through alteration of its structures. For example, we have shown a mutation that weaken the TCRαβ–CD3 ECD *cis*-interaction under force amplifies TCR catch bond and enhances T cell effector function, which suggests a strategy that may be more advantageous compared to mutations at the pMHC docking interface because mutations at the Cβ–CD3 ECD binding interface are not expected to alter the TCR specificity but the same mutation may be effective to different TCRs specific for different tumor antigens. By comparison, mutations at the TCR binding interface may be applicable to a specific pMHC only, and may be riskier in terms of cross-reactivity to self-pMHCs. Thus, rational design guided by catch bond models may provide new TCR engineering strategies that warrant future studies.

## Methods

### Cells and proteins

Naïve CD8^+^ T cells were purified by negative selection from spleens of 2C transgenic mice housed in the animal facility of Emory University following a protocol approved by the Institute Animal Care and Use Committee of Emory University as described^4^. Human red blood cells (RBCs) for BFP experiments were isolated from the whole blood of healthy volunteers according to a protocol approved by the Institutional Review Board of Georgia Institute of Technology as described^4^. Mouse 58^-/-^ T cell hybridoma cells^77^ expressing mouse CD3 but not TCRαβ were a generous gift from Dr. Bernard Malissen (Centre d’immunologie de Marseille-Luminy, France). WT or MT mouse 2B4 TCR were re-expressed on 58^-/-^ cells through retroviral transduction, which were cultured as described^37^. The transduced cells were stained with PE anti-mouse CD3ε (clone 145-2C11 or 2C11, eBioscience) and allophycocyanin (APC)-conjugated anti-TCRβ (clone H57-597 or H57, eBioscience) mAbs and sorted for dual expression of CD3 and TCR. The sorted cells were expanded for 6 days and quantified for TCR and CD3ε expression.

C-terminally biotinylated WT and β_2_m swapping hybrid H2-K^b^ presenting the R4 peptide (SIYRYYGL) were from the National Institutes of Health Tetramer Core Facility at Emory University. To prevent CD8 binding, the MT H2-K^b^α3A2 (with the mouse α3 domain swapped to that of the HLA-A2) was used. Inclusion bodies for I-E^k^ α (with C-terminal biotinylation sequence) and β chains were produced in One Shot™ *E. coli* BL21 (DE3), refolded with K5 peptide (ANERADLIAYFKAATKF), and purified as described previously^78^.

### BFP bond lifetime measurement

A previously described BFP force-clamp assay was used to measure TCR–pMHC bond lifetimes in a range of constant forces at room temperature^4^. Briefly, pMHC was coated onto streptavidin-conjugated glass beads *via* biotin-streptavidin coupling. A pMHC-coupled bead was attached to a biotinylated RBC aspirated on a glass micropipette to form a force probe to test binding with a primary T cell or hybridoma expressing the specific WT or MT TCR in repetitive cycles. In each cycle, the cell was driven by a piezo translator controlled by a computer program to approach and briefly (∼0.1 s) contact the probe bead with a small impingement force (∼10 pN) to allow bond formation, followed by retraction of the cell at a force loading rate of 1000 pN/s. If a bond was detected at a preset tension level, the force was clamped until spontaneous bond dissociation. Bond lifetime was measured as the duration of force clamp. To ensure most adhesion events were mediated by single molecular bonds, the adhesion was controlled to be infrequent (≤20%)^79^. Bond lifetimes were measured at forces ranging from 2 – 30 pN, pooled, and binned into >7 force bins (>50 measurements per bin) to reduce system errors and presented as average lifetime and standard error of the mean (s.e.m.). A previously described thermal fluctuation assay was used to measure bond lifetime at zero force, ^80^. Here, instead of retracting the T cell to apply a tensile force as in the force-clamp assay, the retraction stopped when the contact force disappears and the TCR and the pMHC were then allowed to interact *via* thermal fluctuation of the probe bead. Bond association and dissociation were identified from reduction and resumption of thermal fluctuation of the bead position. Individual lifetimes were measured as the duration from fluctuation reduction to resumption.

### *In vitro* T cell activation

Upregulation of CD25 and CD69 on naïve 2C T cells were assayed using 96-well plates pre-coated with WT or hybrid SIYR:H2-K^b^ at 0.1 ug/mL or 1 ug/mL concentrations for 1 hour at 37 °C. Upon addition of naive 2C T cells at 1 million per well, the plates were incubated at 37 °C for 72 hours. Cells were harvested and analyzed for fluorescence staining using PE-conjugated anti-CD69 (clone H1.2F3, BD Biosciences) and PE-cy7-conjugated anti-CD25 (clone PC61, BD Biosciences) by flow cytometry (BD FACSAria).

### Molecular Dynamics simulations

#### Molecular modeling of the hybrid H2-K^b^

Two complex models of human β_2_m and H2-K^b^ were built based on the crystal structure of mouse β_2_m and H2-K^b^. Because of the high sequence identity (68%) and high structural similarity (backbone RMSD <1 Å) between the human and mouse β_2_m, we made *in silico* mutation to replace mouse β_2_m residues by those of human β_2_m to the WT H2-K^b^ (PDB ID: 1G6R), or replace the entire mouse β_2_m by the human β_2_m in the HLA-A2 (PDB ID: 2BNR).

#### Stability comparison between hybrid and WT H2-K^b^ by conventional MD

Upon adding hydrogen atoms and counter ions (∼150 mM NaCl), and solvating the structures in rectangular water boxes (>16 Å from the box edges and protein) by the VMD software package, we obtained two solvated systems – one for the hybrid and the other for WT H2-K^b^ – with dimension of ∼92×82×97 Å^3^. Both systems were first equilibrated with three steps: (1) 10,000 steps energy minimization and 4-ns equilibration simulations under 1-fs timestep with heavy atoms constrained (except difference residues between mouse β_2_m and human β_2_m); (2) 4-ns equilibration simulations under 1-fs timestep with backbone atoms of proteins constrained; (3) 10-ns equilibration simulation under 1-fs timestep without constrains. Subsequently, the production simulations last ∼100 ns with 2-fs timesteps under rigid bond algorithms to relax the models. Energy minimizations and MD simulations were performed with NAMD2 using the CHARM36m force field for proteins under periodic boundary conditions. Temperature was maintained at 310 K with Langevin dynamics and pressure was controlled at 1 atm with the Nosé-Hoover Langevin piston method. Particle Ewald Mesh summation was used for electrostatic calculation and a 12-Å cutoff was used for short-range non-bounded interactions.

#### Modeling and simulation of the TCR-CD3 ectodomain interaction

All simulations were based on the recently published cryoEM structure of a human TCR-CD3 complex (PDB ID: 6JXR)^42^, which shares the same Cαβ, CD3γε, and CD3δε’ with the mouse 2B4 Vαβ and human LC13 Cαβ hybrid TCR used in our experiments. The structure was transmembrane domain truncated, end ACE/NME capped, and missing residues repaired^81^ to form a complete CD3δε’–TCR–CD3γε trimeric ECD complex. Unit cells were built to enclose the molecular systems to be simulated in a physiologically-appropriate and thermodynamically-favorable state. The initial structures were oriented and centered within optimized orthorhombic cells, which were subsequently solvated using the TIP3P water model^82^, counter-balanced using sodium ions, and ionic strength tuned to ∼150 mM with sodium chloride. To achieve a thermodynamically favorable initial state, the unit cell was energy minimized, followed by two equilibration cycles under NTV, then NPT ensembles with the heavy-atom restraints. The systems were then ready for subsequent equilibration and production runs with/without external force applied using GROMACS (version 2019.6)^83-85^ under the AMBER99SB*-ILDNP force field^86^.

Conventional molecular dynamics (CMD, without force) simulations were performed by letting initials freely evolve without any constraints. While in steered molecular dynamics (SMD, with force) simulations, the external constant forces with constant magnitudes (175 pN) along a fixed direction (*z*-axis) were added to the ECD initials. The N-terminus of the TCR α chain was pulled; in the meantime, the C-termini of CD3ε chains were fixed to mimic the anchor effect of their transmembrane domains. Every simulated trajectory consists of a 100-ns equilibration and a 100-ns production stage. The conformations per 2 ps during the production stage were analyzed to obtain the center-of-mass distances between interested residues. The snapshots per 400 ps during the same period were extracted for visual comparisons.

### Modeling

The model developments, characterization, and validation are described in the main text with more details in the Supplementary Model Derivations. Model fitting to experimental data was done by nonlinear curve-fitting in the least-squares sense using the Levenberg-Marquardt algorithm, as detailed in Supplementary Model Derivation, A.4. Model applications, curve-fitting strategies, and biological relevance. Statistical and clustering analyses were done using Bayesian statistics and Lloyd’s algorithm, respectively. All analyses were done using MATLAB. All published experimental data of force *vs* bond lifetime were measured at room temperature as reported in Refs. ^4-6,8,9,13,16,28,29^ and the TCR–pMHC constructs used were described in the footnotes of Supplementary Tables 3, 4 and 6.

## Data availability

All data are included in the article’s main body and Supplementary Information. Previously published bond lifetime data^4-6,8,9,13,16,28^ re-analyzed for model fitting are summarized and deposited in Github (https://github.com/Chengzhulab/NCOMMS-22-20167).

## Code availability

All code for three models used in this study are summarized and deposited in Github (https://github.com/Chengzhulab/NCOMMS-22-20167).

## Acknowledgments

We thank Jinsung Hong, Peng Wu, and Tongtong Zhang for sharing their published BFP experimental data for re-analysis and model fitting. We also thank Brian Evavold (Emory University) for providing 2C T cells and Bernard Malissen (Centre d’immunologie de Marseille-Luminy, France) for sharing 58^-/-^ hybridoma cells. We acknowledge the National Institutes of Health Tetramer Core Facility at Emory University for providing the pMHC molecules. The MD simulations were supported by an NSF award (MCA08X014) in advanced computing infrastructure for U.S. and performed in the Extreme Science and Engineering Discovery Environment (XSEDE). This work was supported by a Postdoctoral Fellowship from the National Research Foundation of South Korea (2021R1A6A3A03038382 to H.-K.C.), by a National Natural Science Foundation of China grant (31971237 to W.C.), and by a National Natural Science Foundation of China grant (32090044 to J.L.). This work was also supported by National Institutes of Health grants (U01CA250040 to C.Z., U01CA214354 and R01CA243486 to C.Z. and M.K., and R01GM124489 to M.K. and C.Z.).

## Author contributions

H.-K.C. and C.Z. conceived the project and developed the models. P.C., C.G., B.L., and A.N. performed experiments. P.C., Y.Z., and J.L. performed MD simulations. H.-K.C., K.L., and M.N.R. provided data for re-analysis prior to their publication. H.-K.C. performed model analysis and data fitting. H.-K.C., K.L. and C.Z. made key observations and conclusions from analyses. W.C., M.K., J.L., and C.Z. supported the project. H.-K.C., K.L., and C.Z. prepared the manuscript with input from other authors.

## Ethics declarations

No authors have competing interests to declare.

## Supplementary Information

### Supplementary Model Derivations

#### A. Kinetic model for TCR–pMHC-I bonds

##### A.1. Derivation of the force-induced energy change δ_1_γ(F)

The purpose of this subsection is to provide detailed derivation of the force-induced change in the energy landscape, or the work done by force as the TCR–pMHC complex is stretched at the two ends from the bound state to the transition state until dissociation (Fig. 2a, b). Mathematically, *δ*_1_*γ*(*F*) is an integral over a force range 0 ≤ *f* ≤ *F*, where *F* is the level of force under which the kinetic rate is being evaluated (Eq. 2). The integrand *δ*_z_(*f*) is the projection on the force direction of the length change of the TCR–pMHC structure relative to its bound state during dissociation. Subtraction of this work from the interaction free energy tilts the energy landscape that governs the off-rate (Eq. 1, Fig. 1b). To calculate molecular stretch, we assume the TCR–pMHC complex to behave as a system of semi-rigid bodies of globular domains connected by semi-flexible polymers (Fig. 2b). As such, the total length change includes three components: 1) extension of individual globular domains, 2) various domain rotations about hinges, and 3) unfolding of secondary structures at specific regions.

For a globular domain without unfolding, the force-extension relationship is described by the three-dimensional freely-jointed chain model^1^:

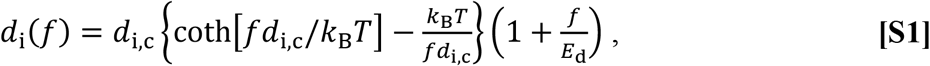

where *d*_i_ is end-to-end distance of the *i*th-domain, *k*_B_ is the Boltzmann constant, *T* is absolute temperature, *f* is force, and *E*_d_ ∼ 100 pN is the elastic modulus of the folded globular domain^2^. The present work considers domain extension of the whole TCR–pMHC-I complex (*d*_N_) and three of its parts: the MHC α_3_ domain (*d*_α3_), bonding interface that includes the MHC α_1_α_2_ domains bound to the TCR Vαβ domains (*d*_B.I_), and the TCR Cαβ domains (*d*_Cαβ_). Their contour lengths, *d*_i,c_, have well-defined values depending on the TCR–pMHC complex in question, which are summarized in Supplementary Table 1. For example, *d*_N,c_ = 12.7 and 10.9 nm for 2C TCR complexed with H2-K^b^ (PDB codes 2CKB, 1MWA, and 1G6R) and H2-L^d^m31 (2E7L), respectively, and 12.3 nm for 1G4 TCR complexed with HLA-A2 (2BNR and 2BNQ). 12.7 nm and 12.5 nm were used for P14 TCR (5M00) and NP1-B17 TCR complexed with H2-D^b^ (5SWZ), respectively. We also choose 11.6 nm as a reasonable guess value for the N15 TCR/OT1 TCR–p:H2-K^b^ complex (because average *d*_N,c_ across 33 structures of TCR-pMHC class I complexes is 11.6 ± 1 nm (mean ± sd)^3^).

To calculate the work *δ*_1_*γ*(*F*), we project the above domain extensions onto the direction of force, which is taken as the *z* direction (Fig. 2b) using two angles between the *z* axis and: 1) the normal direction of the bonding interface (*θ*) and 2) the line connecting the C- and N-termini of the MHC-I α_3_ domain excluding any unfolded residues (*φ*):

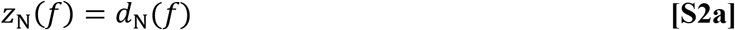

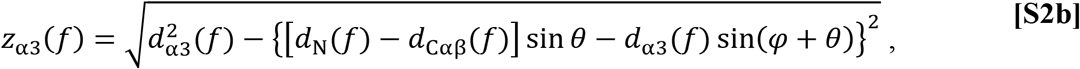

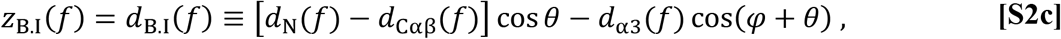

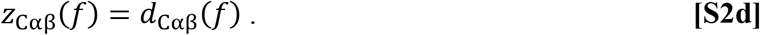

Note that *θ* is a model parameter as it describes the tilting of the bonding interface as part of the force-induced conformational change, whereas *φ* is a model constant is measured from the crystal structure (Supplementary Table 1, see A.2 below).

We assume that partial unfolding in the molecular complex may occur at connecting regions of globular domains, in particular, the α_1_α_2_–α_3_ joint of the MHC-I and the Vα-Cα interdomain joint of the TCR (Fig. 2a). The former may be caused by dissociation of the noncovalent α_1_α_2_–β_2m_ interdomain bond, which shifts the mechanical load originally borne by this bond to the α_1_α_2_–α_3_ hinge, resulting in its partial unfolding, as observed in SMD simulations^4^ (Fig. 2a). Similarly, α_1_α_2_–β_2m_ dissociation results in tilting of the bonding interface and load shifting from the Vβ-Cβ joint to the Vα–Cα joint, leading to partial unfolding of the latter joint (Fig. 2a). The unfolded polypeptides are flexible and can bear only tension but not moment, ensuring that their extension is along the direction of force, i.e., the *z* axis.

The force-extension relationship of the unfolded polypeptides can be described by an extensible worm-like chain (eWLC) model^5^:

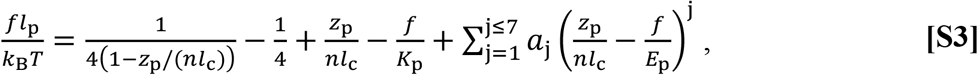

where *z*_p_ is the extension of the unfolded coil under force with the subscript *p* indicating unstructured polypeptide, *l*_c_ = 0.36 nm and *l*_p_ = 0.39 nm are the average contour length and persistence length per unfolded amino acid, respectively^6-8^, *E*_p_∼ 50 μN is the elastic modulus of polypeptides^8^, *a*_j_ are polynomial coefficients for the improved approximation, and *n* is the number of constituent amino acids in the unfolded polypeptide. In particular, we denote the respective numbers of amino acids in the unfolded MHC-I α_1_α_2_–α_3_ and TCR Vα-Cα joints to be *n*_p,MHC_ and *n*_p,TCR_. Eq. S3 defines *z*_p_ as a function of *f*, which can be solved numerically to express in an explicit form: *z*_p_/*nl*_c_ = *z*_u,p_(*f*) = the extension per unit contour length for the polypeptide under force *f*.

Thus, the length of the TCR–pMHC complex at the transition state is (Fig. 2b):

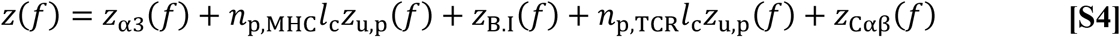

Since we do not have prior knowledge about either number of unfolded amino acids, we will evaluate their sum *n*^*^ = *n*_p,TCR_ + *n*_p,MHC_ from curve-fitting of our model to the experimental data (see below). Since *d*_N_(*f*) is the length of the TCR–pMHC complex at the bound state (Fig. 2b), we have *z*(*f*; 0) = *d*_N_(*f*). Finally, the integrand on the right-hand side of Eq. 3 can be written as

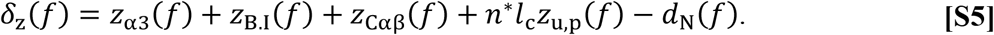

##### A.2. Simplifying assumptions and reducing parameters

The purpose of this subsection is to provide the details omitted in the main text of how the model parameters are reduced to the smallest possible set and the underlying simplifying assumptions. To begin, we take the *φ* values from PDB for the specific TCR–pMHC interactions in question (Supplementary Table 1). We assume that the *φ* value remains constant during forced dissociation due to their small range, *e.g*., *φ* = 12.7° and 5° for 2C TCR complexed with H2-K^b^ (PDB codes 2CKB, 1MWA, and 1G6R) and H2-L^d^m31 (2E7L), respectively, and 13.3° for 1G4 TCR complexed with HLA-A2 (2BNR and 2BNQ), 15° for both P14 TCR (5M00) and NP1-B17 TCR complexed with H2-D^b^ (5SWZ). We also chose 23.5° as a reasonable guess value for the N15 TCR/OT1 TCR–p:H2-K^b^ complex. Additionally, due to the semi-rigid approximation, the thickness of the constant domain of TCR α- and β-subunit (*d*_Cαβ_) is also treated as constant for each construct (Supplementary Table 1). This leaves only two structural parameters (*d*_α3_, *θ*) in our model to be evaluated by curve-fitting to data. Noting that the structure-based force function *γ*(*F*; *δ*_1_) scales with the characteristic extension change per unit change of molecular length such that 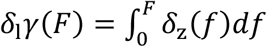, the dissociation rate of TCR–pMHC-I bond can be written as:

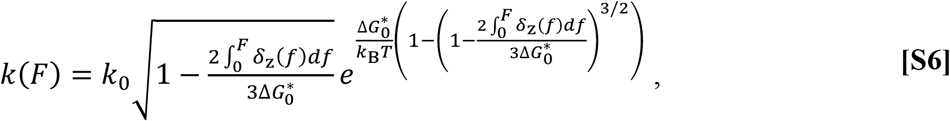

where *δ*_z_(*f*) is given by Eq. S5.

To further reduce model parameters, we make additional assumptions as discussed in the main text. It is well-known that the fractions of free-energy change in biological interactions in liquid, such as unfolding and refolding of proteins, unbinding and rebinding of receptor– ligand bonds, and unzipping and rezipping of RNA or DNA, are small because of their limited dynamic transition time^9-12^. Such a rate limit results from the nature of biological interactions, *e.g*., polar/non-polar interactions, hydrophobic interactions, and charged interactions, which typically yield finite range of transition kinetics. This enables to roughly estimate the free-energy barrier as 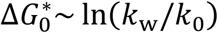 where *k*_w_∼10^6^ s^-1^ known as the prefactor^9-12^.

##### A.3. Defining the dissociation coordinate

The purpose of this subsection is to derive an operational way to determine the dissociation coordinate variable *δ*_1_ used in the main text. We note that the total number of unfolded amino acids *n*^*^ is zero at the bound state before unfolding occurs, increases monotonically during progressive unfolding along the dissociation path, and reaches maximum at the dissociation point. Because *n*^*^ is not known *a priori*, it must be treated as a fitting parameter similar to *d*_α3_, *θ*, and 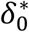. Since *δ*_1_ is the contour length change along the dissociation path, we wish that *δ*_1_ approaches its upper bound 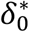 and depends on force implicitly through the model parameters *n*^*^, *d*_α3_, and *θ*, as given below:

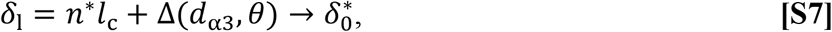

where Δ(*d*_α3_, *θ*) is the difference of the contour lengths except for the partially unfolded regions. Thus, Eq. S7 provides a constraint for *δ*_1_ instead of introducing another model parameter. *d*_α3_ and *θ* are determined for each *n*^*^ during the model fitting that searches for parameters to enable 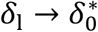. Small errors, which may occur for the various contributions to Δ (red lines in Fig. 2a labeled as force transmission lines), can be identified using a pair of (*d*_α3_, *θ*) values and the crystal structure for each complex. Specifically, average differences ⟨Δ⟩ are - 0.8 nm for the strong catch bond (SC) group (*d*_α3_ > 3 nm, *θ* > 30°), -0.4 nm for the weak catch bond (WC) group (1 nm < *d*_α3_ < 3 nm, 8°< *θ* < 30°), and 0 nm for slip-only bond (SO) group (*d*_α3_ < 1 nm, *θ* < 8°), respectively (see Fig. 4e-i and associated text for the definitions of SC, WC, and SO groups). It has been well established that the contour length of a single amino acid is ∼0.4 ± 0.02 nm/a.a^7,13,14^, implying that the model has a resolution of 2 amino acids. We further note that, even without conformational change, it is possible for the slip-only bond group to have 3 unfolded amino acids due to the limited resolution of the model. Finally, the best-fit model parameters can be determined by finding the subset of best-fit parameters among possible *n*^*^ values that match the contour length change *δ*_1_ to the free-energy well width at zero-force 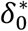, *i.e*., finding *n*^*^ such that 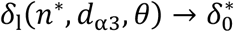 (see Supplementary Table 2). Under this condition, 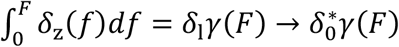 and Eq. S6 becomes identical to Eq. 2.

##### A.4. Model applications, curve-fitting strategies, and biological relevance

The purpose of this subsection is to outline the procedures of applying our model to experiments. Our procedures include four steps: 1) examine how the model parameters control the model behaviors, 2) fit the model-predicted reciprocal off-rate (Eq. S6) to the experimental bond lifetime *vs* force data, 3) construct the energy landscape and investigate its properties based on the parameters evaluated in part 2, 4) elucidate the biological relevance of the model parameters.

To examine how the model parameters control the model behaviors, we varied one parameter while keeping others constant. For example, to investigate the effect of varying the titling angle *θ*, we kept the other parameters constants (*i.e*., *n*^*^ = 7, and 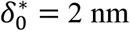, *k*_0_ = 3 s^-1^ and 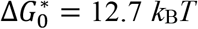), Also kept constant were several structural constants (*φ* = 15°, *d*_*N*_ = 12 nm and *d*_Cαβ_ = 3.5 nm) introduced in section A.1. Of note, *d*_α3_ varies as *θ* changes because of a pulling constraint (see section B.1). With a fixed *θ* (*i.e*., 30°) the effect of molecular extension at zero force was investigated by varying extension from 0.5-3.5 nm. These were selected by the average values of actual fitting results, which had been tested and confirmed as reasonable.

The free-energy landscape can be constructed by substituting the best-fit model parameters into the following equations^14^:

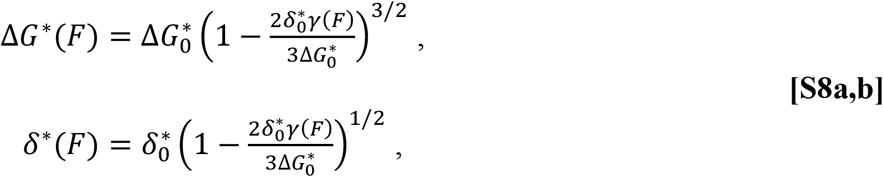

Thus, by using model parameters, the dissociation state coordinates relative to the bound state coordinates in the free-energy *vs* dissociation coordinate space can be defined as functions of force. Note that this force-induced change of the barrier height should be under the condition of small perturbation such that 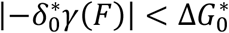. Since our fitting results show that the average free-energy barrier height at zero force is ∼12 *k*_B_*T* 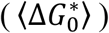, the force range corresponding to a change of the barrier height of <10 *k*_B_*T* is reasonable for each dataset, *i.e*., force range corresponding to energy barrier heights in the range of 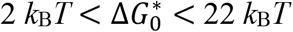.

Firstly, to fit the model-predicted reciprocal off-rate (Eq. S6) to the experimental bond lifetime *vs* force data, all four parameters were changed simultaneously to search for the minimum of the chi squared error. Second, every fitting curve was then determined by the parameter set that has the closest *δ*_1_ value to 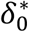 (described as Eq. S7 in section A.3 and Supplementary Table 2). The error bars of the best-fit parameters (or their ranges) were calculated from the standard errors of mean of bond lifetimes and the residual matrices. Of the 55 datasets analyzed, only one has 4 data points and this dataset shows slip bond; as such, is governed by two fitting parameters because the other two parameters are nearly zero. All other datasets have 6-10 data points; therefore, over-fitting is not a problem.

To elucidate the biological relevance of the model parameters *θ, n*^*^, *d*_Cαβ_, 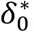, and 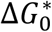, we examine their changes with varying bond lifetime *vs* force data obtained from different TCR and pMHC interactions that induce a wide range of biological responses. Finding correspondence between a group of model parameters individually and/or collectively with the biological response would be considered as support for the biological relevance of the model, because such correspondence suggests that the model can discriminate different TCR–pMHC interactions that result in differential T cell functions.

##### A.5. Class I model constraints

The purpose of this subsection is to check whether the model parameters obtained from data fitting are consistent with the constraints to which the model is subjected. Our experiments applied tensile force through the two ends of the TCR–pMHC complex, such that the force direction would always align with the line connecting the C-termini of the respective TCR and MHC molecules during dissociation, giving rise to the so-called pulling constraint. To formulate this pulling constraint in our model, we note that the pulling line is maintained so that the coordinate perpendicular to the force direction is invariant. As depicted in Supplementary Fig. 4a, several angle and length variables can be related using model parameters and structural constants by:

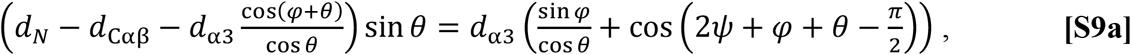

where *ψ* is the angle in an isosceles triangle constructed by rotating the α3 domain. By assuming that the α3 domain would be aligned with force, we estimate that the angle in the last term is near 90°, *i.e*., 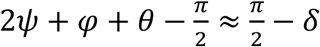 where 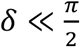. Under this small angle assumption, Eq. S9a can be approximated by:

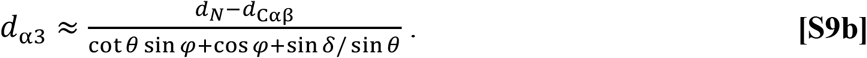

Upon inversing Eq. S9b, we found that the tilting angle *θ* is a function of the end-to-end distance of the α3 domain, *i.e*., *θ* = *f*(*d*_α3_). Setting *δ* = 25°, which seems reasonable as it approximates the maximum value of *φ*, the structural parameters obtained by fitting are scattered in-between two black curves on the *d*_α3_ − *θ* plane marked as pulling constraint: [*d*_*N*_ = 13.5 nm, *d*_Cαβ_ = 2.75 nm, *φ* = 0°] and [*d*_*N*_ = 10.5 nm, *d*_Cαβ_ = 4.25 nm, *φ* = 25°] (Fig. 4f).

Another constraint is the tilting constraint resulted from the asymmetric unfolding and stretching of the interdomain links between the TCR constant and variable domains. This constraint is introduced in the model to account for the potential regulatory effect of the FG-loop on catch bond. Notwithstanding the total number of unfolded amino acids *n*^*^ can be determined by the validation procedure demonstrated in A.2, its breakdown into the number of unfolded amino acids for MHC (*n*_p,MHC_) and TCR (*n*_p,TCR_) remains undetermined. To do this, known structures from *PDB* were used to determine *n*_p,MHC_ first. Briefly, by matching *d*_α3_ (Supplementary Fig. 4b, c) with the end-to-end distance between C-terminal end of α3 domain and certain point following known *PDB* structure, the exact starting position of partial unfolding in MHC can be found following additional assumption that unfolding starts from C-terminal end of α2 domain towards α3 domain (Supplementary Fig. 4b). Thus, *n*_p,TCR_ can be simply calculated as *n*_p,TCR_ = *n*^*^ − *n*_p,MHC_. Upon combining all information, the tilting angle (*θ*_TCR_) of variable domains of TCR can be described by simple trigonometrical function:

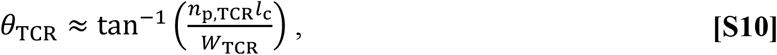

where *W*_TCR_ is width of two interdomain hinges of the TCR α- and β-subunits measured from the crystal structures (Supplementary Fig. 4a, tilting constraint). In this work we use *W*_TCR_ = 3.7 ± 0.3 nm as a representative width due to structure-to-structure variations. Thus, by comparing the tilting angle of the bonding interface *θ* (model parameter) to the tilting angle of the TCR *θ*_TCR_ (derived from another model parameter *n*_p,TCR_ and structural constants *W*_TCR_ and *l*_c_), we can check the validity of tilting constraint using linear regression in the *θ vs θ*_TCR_ p1ot (Supplementary Fig. 4d).

#### B. Kinetic model for TCR–pMHC-II bonds

##### B.1. Development, validation, and characterization

The purpose of this subsection is to present details of the development, validation, and characterization of the TCR–pMHC-II catch bond model omitted in the main text for simplicity, in a similar fashion as the TCR–pMHC-I catch bond model described in Section A. The two models share exact the same framework but have different detailed form of the characteristic extension change *δ*_z_(*f*) (Eq. S5). Comparing to the TCR–pMHC-I complex, the TCR–pMHC-II complex has different docking domains and pulling geometries (one *vs* two transmembrane domains on both TCR and MHC). For this reason, we assume that the force-induced bonding interface tilting angle (*θ*) would be much smaller in the TCR–pMHC-II than TCR–pMHC-I complex. The extension at the bound state can be defined as the end-to-end distance between both end-points identified by crystal structures (E8: 2IAM, 2IAN and 2B4 (as the substitution of 3.L2): 6BGA, 3QIB):

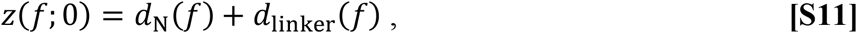

where *d*_N_ is set to be 12.3 nm based on the crystal structures and *d*_linker_ ≈ 9.4 nm represents the linker (*e.g*., a leucine zipper) engineered at the C-termini of soluble pMHC-II constructs to stabilize both the MHC α- and β-subunits, which is often used in experiments for measuring TCR–pMHC-II catch bonds. To account for domain rotation resulted from partial unfolding inside the TCR–pMHC-II complex, we introduce one more variable, *φ*, as the tilting angle of the TCR constant domains. Using the tilting constraint similarly to that used in the class I model, this angle can be approximately described by structural parameters (*d*_B.I_, *θ*) and model constants (see subsection B.2). In short, each component in the right-hand side of Eq. S11 can be expressed by the model parameters (*d*_B.I_, *θ, n*^*^), and model constants as follows.

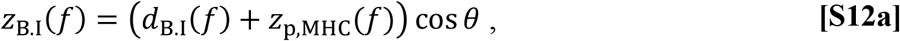

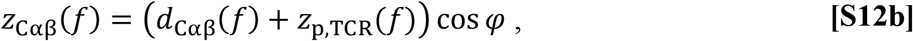

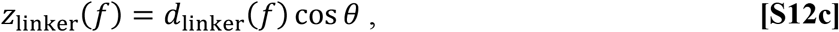

where *z*_p,MHC_(*f*) and *z*_p,TCR_(*f*) are respective extensions of unfolded polypeptides given by *n*_p,MHC_*l*_c_*z*_*u,p*_(*f*) and *n*_p,TCR_*l*_c_*z*_*u,p*_(*f*), respectively, *d*_B.I_ is the length of structure consisting of the MHC and the TCR variable domains, and other parameters defined previously. Finally, the rate coefficient of the TCR–pMHC-II dissociation can be developed by employing the same framework (see Eq. S6). However, when applying the model to experimental data, we can use the constraints of the TCR–pMHC-II interaction to make the model much simpler (see section B.2).

Validation of the class II model follows exactly the same procedure as that used in the validation of the pMHC I model, which is done by checking self-consistent through the definition of the reaction coordinate. By varying the *n*_p,MHC_ from 0 to 10 (see details explained in section B.2), the molecular extension at zero force 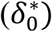 can be estimated by using the contour length-change (*δ*_1_) along the dissociation coordinate:

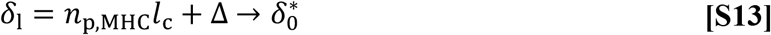

where Δ(*d*_B.I_, *θ*) < 0.2 nm because *θ* < 10°. The best-fit model parameters can be determined by finding a subset of best-fit parameters among possible *n*_p,MHC_ values that match the contour length-change along the dissociation coordinate (*δ*_1_) to the width of the free-energy well at zero force 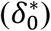, *i.e*., finding *n*_p,MHC_ such that 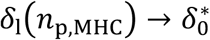 (see Supplementary Table 5).

To characterize the class II model, we examine the model predictions by varying the model parameters one by one while fixing the others as constants. For example, to investigate force-induced bonding interface titling, we fixed the other model parameters (*i.e*., *n*^*^ = 4, 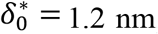, *k*_0_ = 10 s^-1^, and 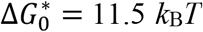) and structural constants (*d*_N_ = 12 nm, *d*_B.I_ = 12.3 nm, and *d*_1inker_ = 9.4 nm for pMHC-II constructs that have linkers). As another example, we fixed *θ* = 0° or 3° and examined the effect of the molecular extension at zero force by varying 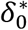 from 0.3-1.8 nm. The constants used for model characterization were selected by their averages from the corresponding values used to fit actual experiments. The parameters for free-energy landscape construction, the energy barrier height (Δ*G*^*^) and energy well width (*δ*\*) as functions of force, are given by Eq. S8, exactly the same as the pMHC I model.

Fitting of class II model to experimental data and examination of the biological relevance of the best-fit parameters were done the same way as the class I model.

##### B.2. Class II model constraints

The purpose of this subsection is to describe the constraints of the class II model, which share similar ideas to those of the class I model (*e.g*., pulling and tilting constraints) but differ in their specific expressions. To formulate the pulling constraint, we again used the fact that the pulling force direction must aligns with the line connecting to the C-termini of the TCR and pMHC molecules so that the coordinate perpendicular to the force direction is invariant. Using model parameters and structural constants, this pulling constraint can be written as:

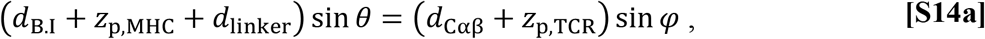

which can be solved for *z*_p,TCR_ explicitly:

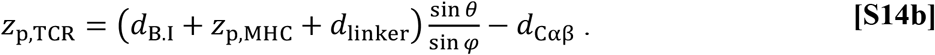

By combining Eq. S12 and S14, the total extension (*z*) can be calculated as the sum of all component extensions (∑_*i*_ *z*_*i*_) at dissociation (*δ*_*l*_ > 0):

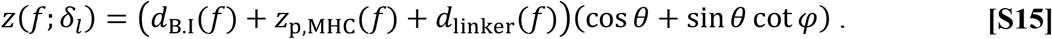

This equation states that only the number of unfolded amino acids in MHC (*n*_p,MHC_) affects extension change during transition. The total number of unfolded amino acids from both TCR and MHC can be estimated by Eq. S14b:

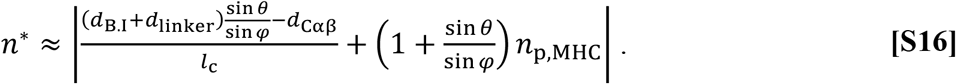

which can be approximately calculated using contour lengths of length components at force-free state.

By assuming small angle perturbation, which seems reasonable, we further reduce the number of model parameters after relating the tilting angle of the TCR constant domain (*φ*) and the titling angle of the bonding interface by the following equation:

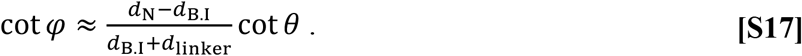

Thus, all terms including *φ* can be re-expressed by using Eq. S17.

Additionally, the tilting constraint can be expressed as follows:

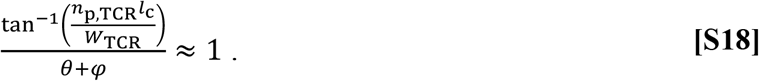

Thus, by using Eq. S15 and S17, only 4 fitting parameters, two structural parameters (*d*_B.I_, *θ*) and two biophysical parameters 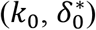, were used to fit the class II model to data.

#### C. A general biophysical limit of model parameters

The purpose of this section is to describe a general biophysical limit that constrains the fitted model parameters, which is used in Fig. 6 to accept the correct model application to data of matched MHC class and reject incorrect model application to data of mismatched MHC class. The idea is that, even if the model is capable of fitting experimental data and the parameters are self-consistent with one another within the model, their values should be within known limits. A prototypical example of such a biophysical limit involves the molecular extension per unfolded amino acids. It follows from Eqs. S7 and S13 that the average molecular extension at zero force over all data (⟨*δ*_0_⟩) should be a linear function of *n*^*^ such that ⟨*δ*_0_⟩ = *an*^*^ + *b*. The *y*-axis intercept *b* can be determined from the slip bond data because slip bonds are not expected to have unfolded amino acids (i.e., *n*^*^ = 0) but still have a nonzero extension. The slope *a* is constrained by the fact that the contour length of a single amino acid has a small range (∼0.4 ± 0.02 nm/a.a.)^7,13,14^; thus, the average molecular extension (⟨*δ*_0_⟩) per unfolded amino acid should be bounded by:

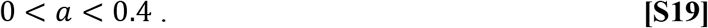

Imposing this range limit of ⟨*δ*_0_⟩ /a.a. would enable us to rule out inappropriate application of the model even if such application could achieve reasonable level of goodness-of-fit. Indeed, all results were below the biophysical limitation.

Conversely, a nearly zero estimate of *a* would indicate that the model is inappropriate for catch-slip bond data because, for the model to fit catch-slip bonds, it requires *n*^*^ > 0 (see Figs. 2c-g and 3e, 3^rd^ row). A parameter estimation of *a* ≈ 0 indicates the lack of dependence of the model behavior on *n*^*^, which abolishes the model’s ability to predict TCR signaling and antigen discrimination, making the model irrelevant to biology.

The sturdier class II than class I pMHC structure also precludes large rotation during conformational change at transition state, leaving only unfolding and stretching along the force. Thus, the average molecular extension per amino acid should be close to 0.4 nm/a.a. On the other hand, *a* ≈ 0 is expected from fitting the slip-only data because such data correspond to the *n*^*^ = 0 case, which makes it difficult to robustly estimate the correct *a* value. Thus, the average molecular extension per amino acid should be well-correlated with each other (*i.e*., high level of goodness-of-fit as measured by *R*^2^) and in the range between 0 to 0.4 nm/a.a. In Fig. 6, we used these criteria to test the appropriateness of cross-applying the class I model to class II data and *vice versa*, showing that it is appropriate to use either model to fit matched data but inappropriate to use either model to fit mismatch data.

## Supplementary Figures

**Supplementary Figure 1.**
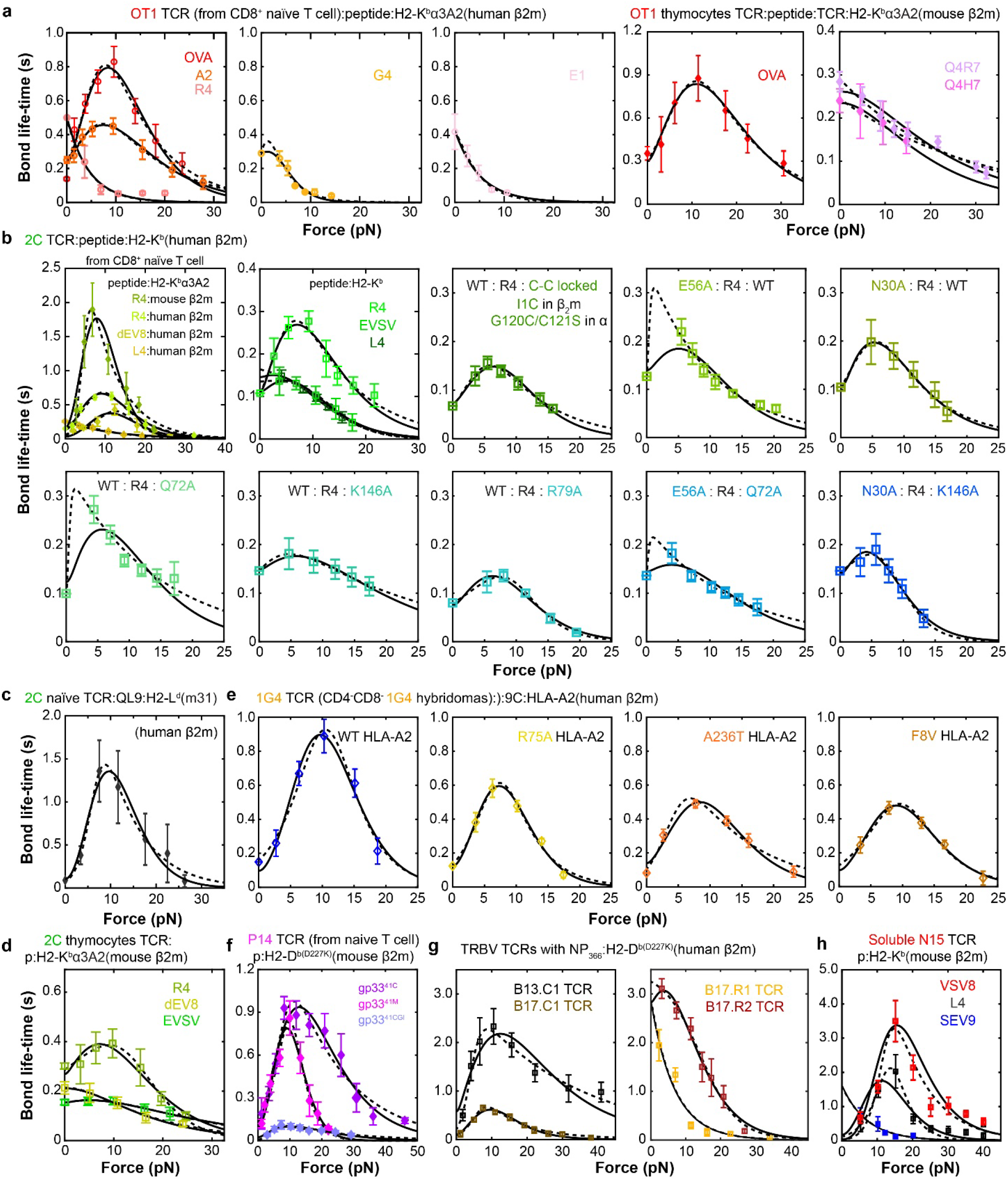
Fitting class I model and two-pathway model to data of TCR– pMHC-I bond lifetime *vs* force. Fitting of theoretical 1/*k*(*F*) curves of the class I model (solid curves) and the two-pathway model (dashed curves) to experimental bond lifetime *vs* force data (*points*, mean ± sem) of 9 TCRs and their mutants interacting with different pMHCs as described below (re-analyzed from Refs. ^4,15-19^). **a** OT1 TCR expressed on CD8^+^ naïve T cells (first three panels) or CD4^+^CD8^+^ thymocytes (last two panels) interacting with indicated p:H2-K^b^α3A2. **b** 2C TCR expressed on CD8^+^ naïve T cells interacting with indicated p:H2-K^b^α3A2 (*top*, 1^st^ panel) or on CD8^-^ hybridomas interacting with indicated peptides presented by WT (*top*, 2^nd^ panel) or disulfate-locked MT (*top*, 3^rd^ panel) H2-K^b^, 2C TCR with indicated point mutations expressed on CD8^-^ hybridomas interacting with R4:H2-K^b^ (*top*, 4^th^ and 5^th^ panels), 2C TCR expressed on CD8^-^ hybridomas interacting with R4 peptide presented by H2-K^b^ with indicated point mutations (*bottom*, 1^st^-3^rd^ panels), or 2C TCR with indicated point mutations expressed on CD8^-^ hybridomas interacting with R4 peptide presented by H2-K^b^ with indicated point mutations (*bottom*, 4^th^ and 5^th^ panels). **c** 2C TCR expressed on CD8^+^ naïve T cells interacting with indicated p:H2-L^d^(m31) with truncated α3 domain. **d** 2C TCR expressed on CD4^+^CD8^+^ thymocytes interacting with indicated p:H2-K^b^α3A2. **e** 1G4 TCR expressed on CD8^-^ hybridomas interacting with indicated p:HLA-A2. **f** P14 TCR expressed on CD8^+^ naïve T cells interacting with indicated p:H2-D^bD227K^. **g** TRBV TCRs of canonical (B13.C1 and B17.C1) and reverse (B17.R1 and B17.R2) pMHC-I docking orientation expressed on hybridomas interacting with NP_366_:H2-D^bD227K^. **h** Purified N15 TCRαβ interacting with indicated p:H2-K^b^. The first panel of a and the first two top panels of b are replotted from **Fig. 3a** and **b** for completeness. See Supplementary Table 3 for a list of the interacting molecules.

**Supplementary Figure 2.**
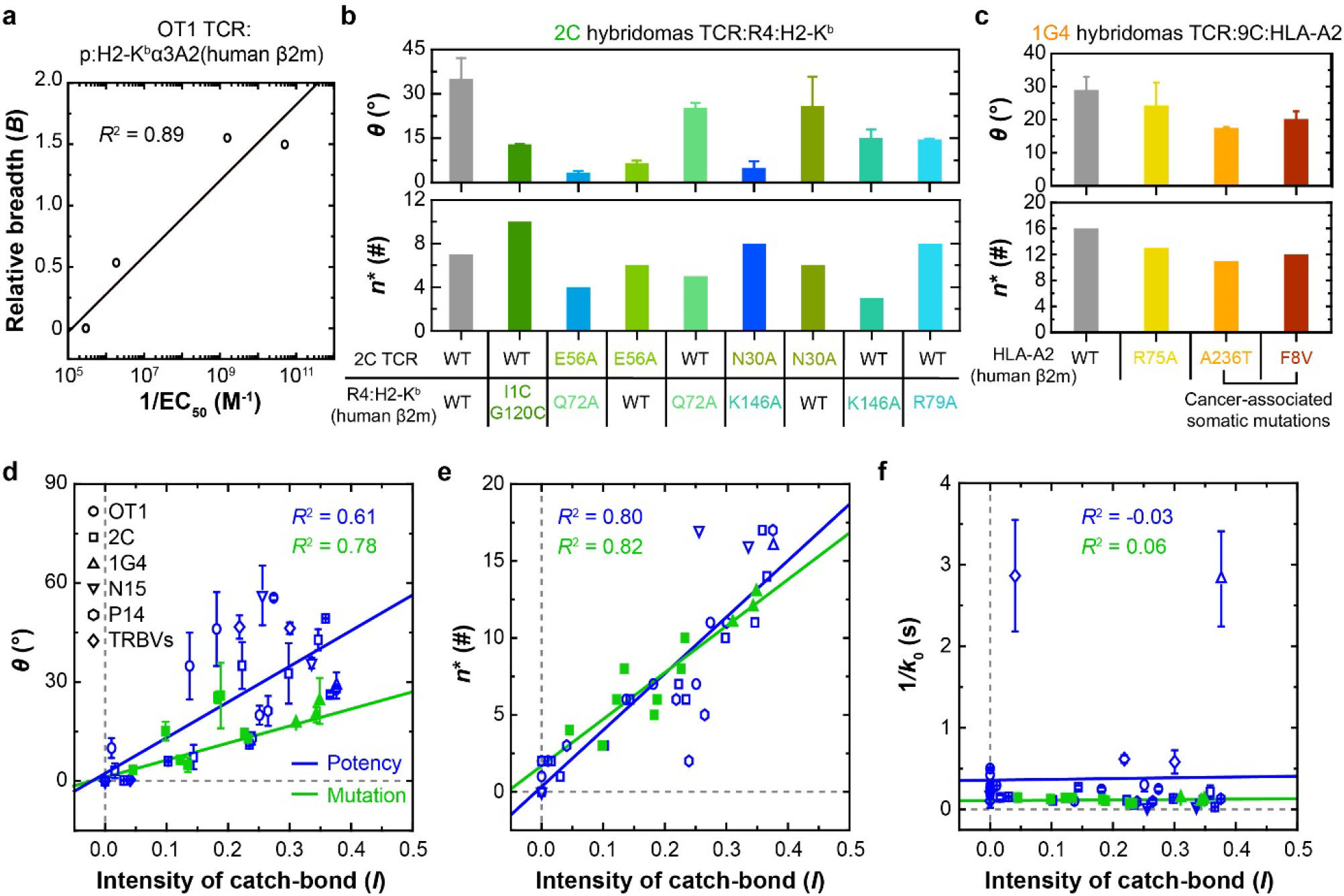
Correlation of model parameters with pMHC-I biological activity and TCR bond type. **a** Relative breadth *vs* the reciprocal peptide dose required to stimulate half maximal of OT1 T cell proliferation. **b**-**c** Model parameters ***θ*** (the tilted angle of the bonding interface, *upper*) and ***n***^*^ (the number of unfolded amino acids, *lower*) that best-fit the data in Supplementary **Fig. 1b** and **d** are plotted *vs* the indicated WT and mutant 2C (**b**) and 1G4 (**c**) TCRs with their indicated pMHCs. **d-f** Scattergrams of ***θ*** (**d**), ***n***^*^ (**e**), and **1**/***k***_**0**_ (**f**) *vs* ***I*** = ***L***/(**1** + ***B***) (catch bond intensity) are plotted using the data from **Figs. 3e, 4b**, and **Supplementary Figs. 2b & 2c** to examine correlation. Blue-open symbols indicate data of known T cell biological activities (ligand potencies) induced by the corresponding TCR– pMHC-I interactions that correlate to catch bond intensity. Green*-*closed symbols indicate data of known effects on catch bond metrics by targeted mutations on the TCR, MHC, or both. See Supplementary Table 3 for a list of the interacting molecules.

**Supplementary Figure 3.**
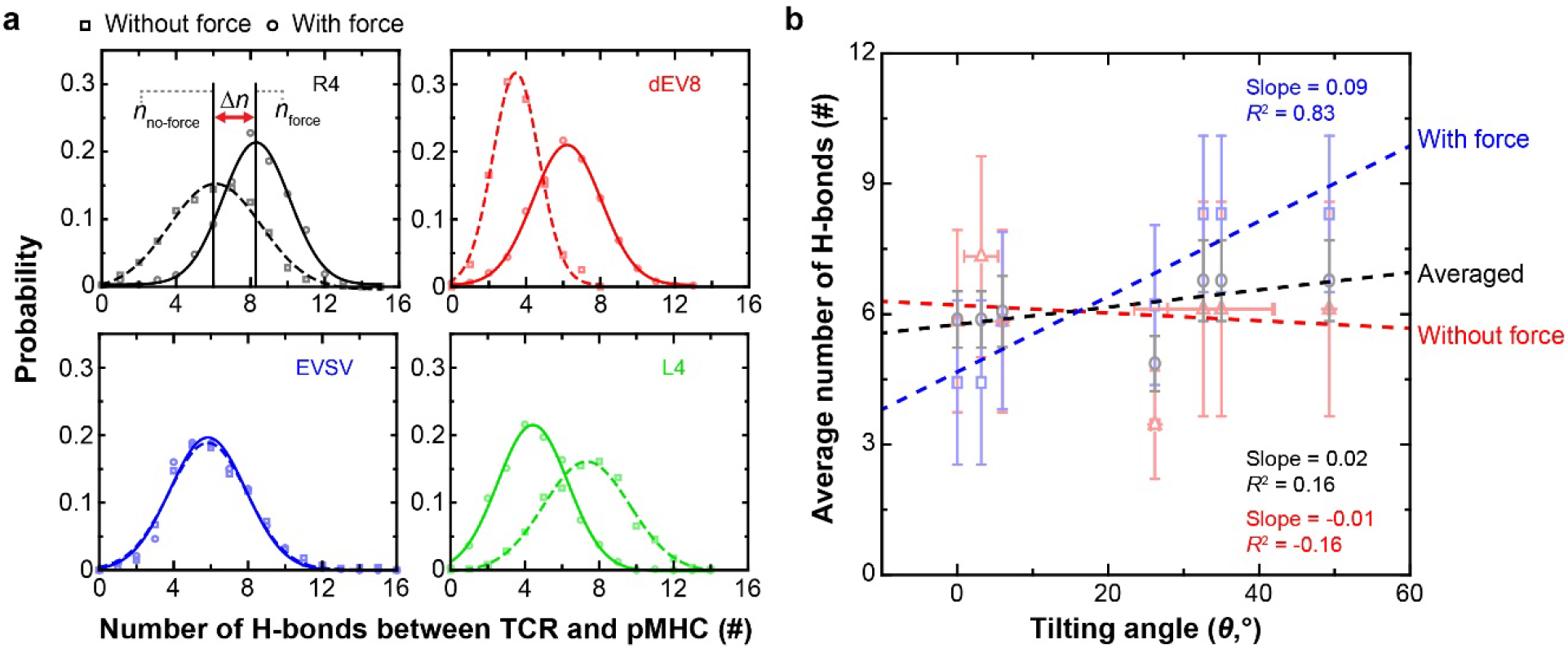
Correlation of force-induced TCR–pMHC-I hydrogen (H) bonds with bonding interface tilting angle. **a** H-bond distributions at bonding interface between 2C TCR and the indicated pMHCs (R4, dEV8, EVSV, or L4) in the presence (*solid curves*) or absence (*dotted curves*) of the force (obtained by re-analysis of SMD simulation results from Ref. ^**4**^ and fitted by Gaussian functions). **b** Plots of average number of H-bonds (mean ± sd) with respect to fitted tilting angle (with fitting error) of bonding interface in the absence (*red*) and presence (*blue*) of force as well as their average (*gray*). Average numbers of H-bonds were determined as mean value for each Gaussian distribution at each titling angle. All colored dashed lines are linear fits.

**Supplementary Figure 4.**
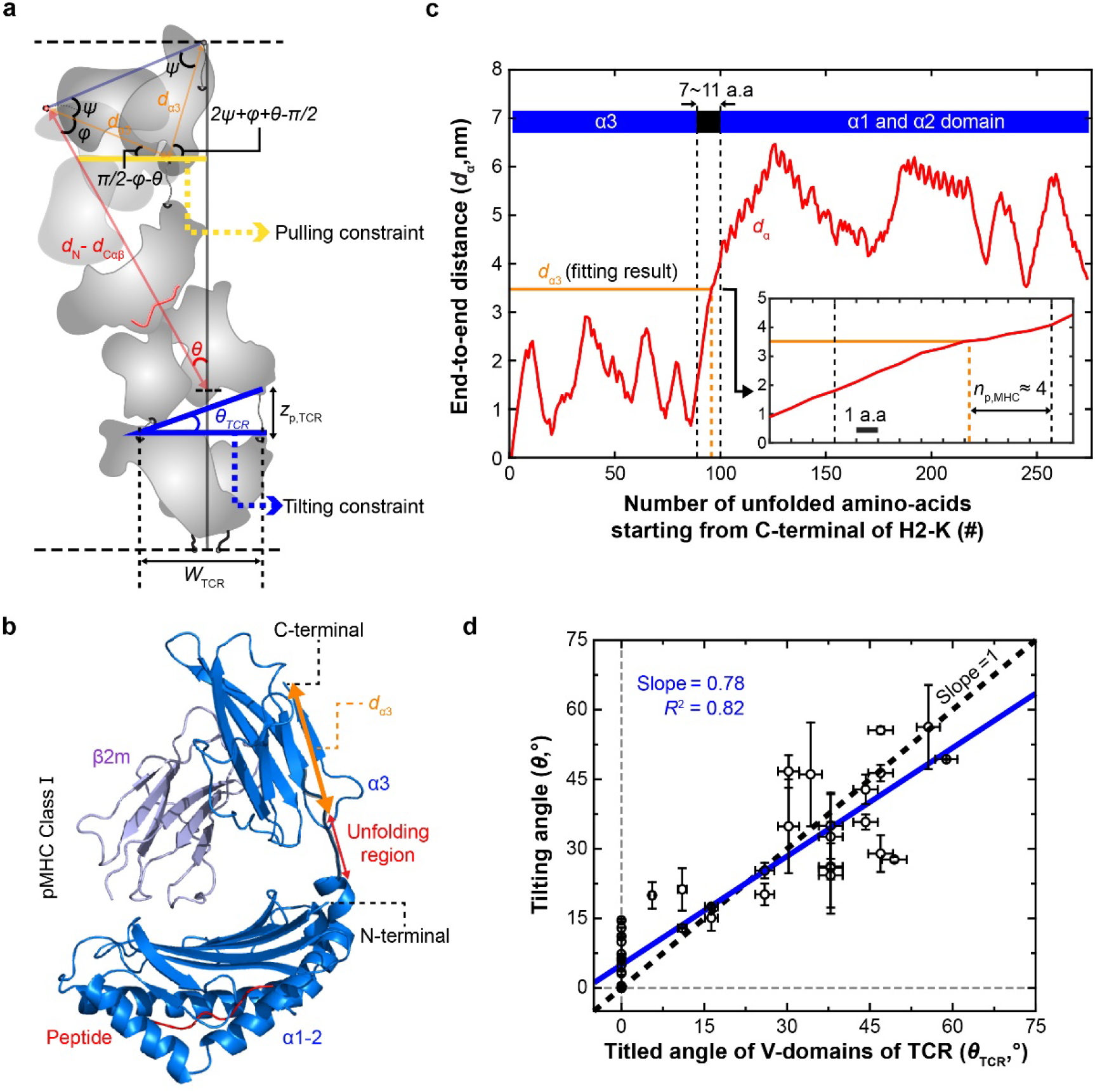
Pulling and tilting constraints of class I model. **a** An illustration of both pulling and tilting constraints. Length of yellow line was used to set up the equation of the pulling constraint between ***d***_**α3**_ and ***θ***. The tilted angle from the blue lines was used to check asymmetric stretching of TCR. **b** A pMHC-I structure based on PDB code 2CKB. Partial unfolding of MHC is assumed to start from the end of the α1-α2 domain towards the α3 domain as suggested by SMD simulation^**4**^. **c** Representative end-to-end distance of ***d***_**α3**_ *vs* the number of unfolded amino acids. The distance is calculated from C-terminal end of the **α3** domain based on the structure shown in (**b**). Distance is calculated using the PBD structure for each construct. ***d***_**α3**_ distance obtained from fitting the model to data is converted to ***n***_**p**,**MHC**_ (*inset*). **d** Tilted angle of the TCR variable domains (***θ***_**TCR**_) *vs* the titled angle of bonding interface (***θ***) as one of the fitting parameters. The validity of asymmetric stretching of TCR was checked by the linear relationship between the two angles.

**Supplementary Figure 5.**
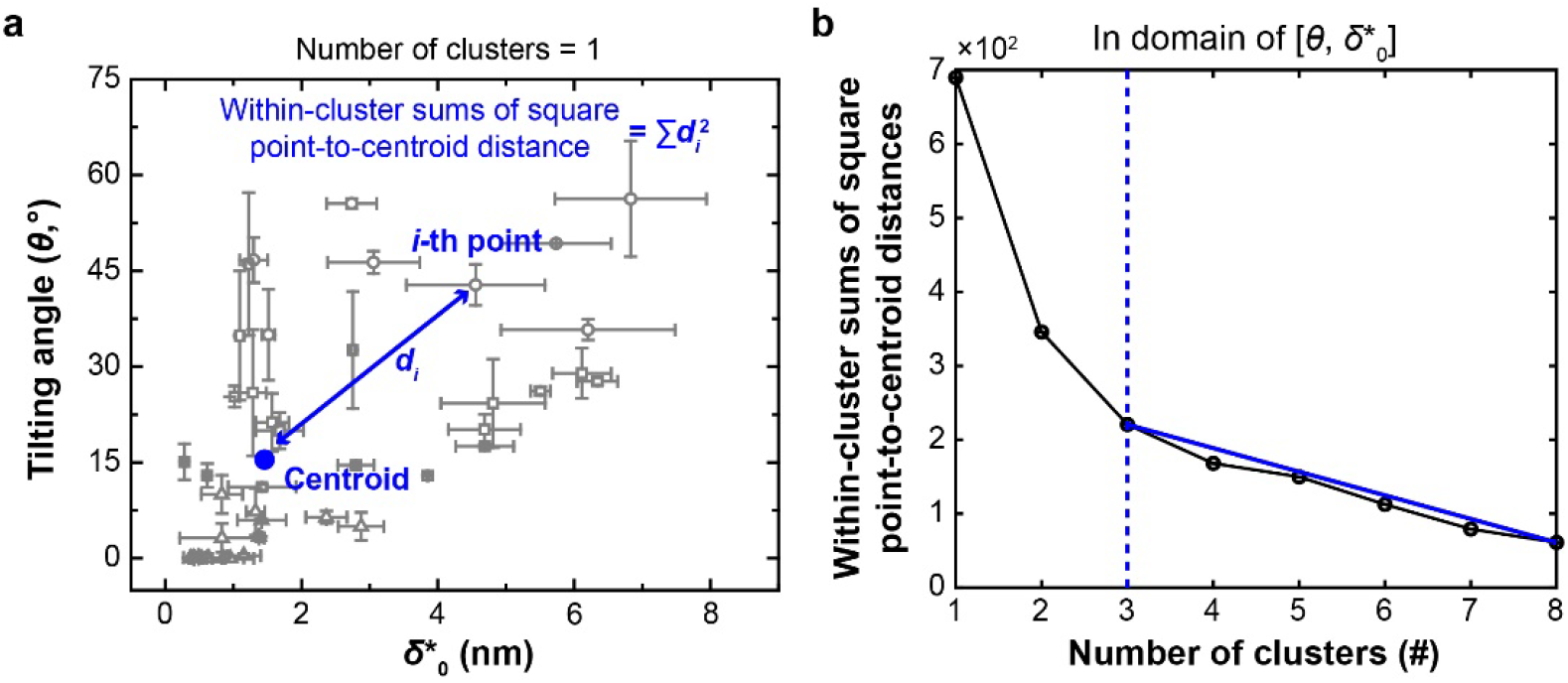
Determining the number of clusters in parameter space. **a** Definitions of the *i-*th point-to-centroid Euclidean distance (blue line) and of the within-cluster sums of square point-to-centroid distance when the number of clusters is 1. **b** Sum of all squared Euclidean distances *vs* number of possible clusters. Point of abrupt reduction of steepness of slope leading to a straight line (*blue solid-line*) was identified a number of clusters (*blue dotted-line*).

**Supplementary Figure 6.**
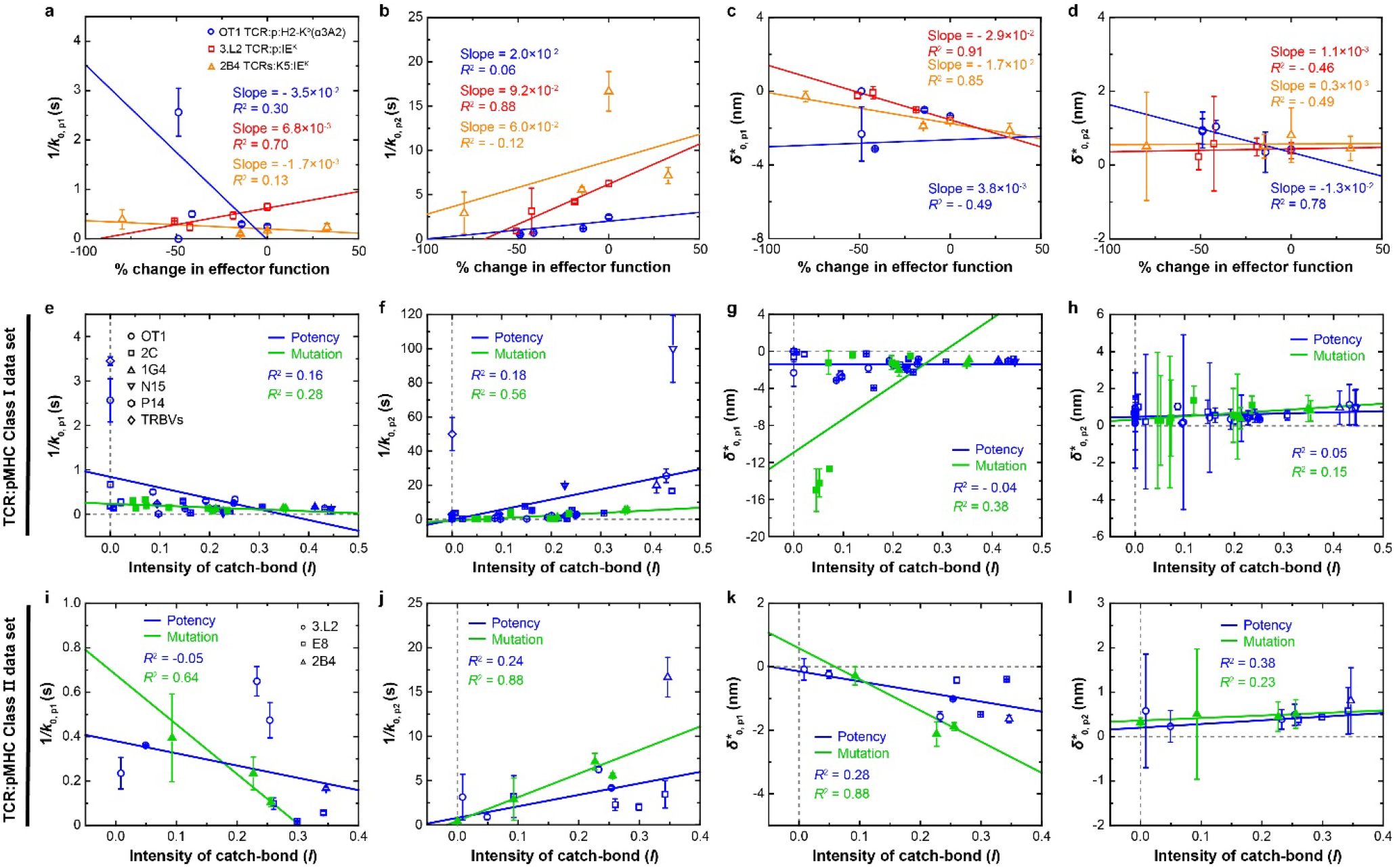
Lack of correlations between best-fit parameters of the two-pathway model and T cell function. **a-d** Correlation plots of **1**/***k***_**0**,***p*1**_ (**a**), **1**/***k***_**0**,***p*2**_ (**b**), 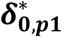 (**c**), and 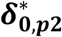 (**d**) *vs* % changes in effector function. T cell effector function was defined by the peptide dose required to achieve half-maximal proliferation (1/EC^50^) of OT1 T cells (*blue*) or to generate 40% B cell apoptosis (1/EC^40^)^**20**^ for 3.L2 T cells (*red*), or the area under the dose response curve (AUC) of 2B4 hybridoma IL-2 production^**21**^ (*orange*). Color-matched solid lines are linear fits for the indicated TCR–pMHC systems. Also indicated are the slopes and *R*^2^ values. **e-l** Correlation plots of **1**/***k***_**0**,***p*1**_ (**e, i**) **1**/***k***_**0**,***p*2**_ (**f, j**), 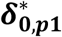 (**g, k**), and 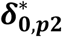 (**h, l**) *vs* the intensity of catch bond evaluated from the indicated class I (**e-h**) or class II (**i-l**) datasets. Solid lines are linear fits, which are color-matched to indicate the ‘potency-’ or ‘mutation-’ related datasets for the class I system. All error bars are fitting error (Supplementary Table 4).

**Supplementary Figure 7.**
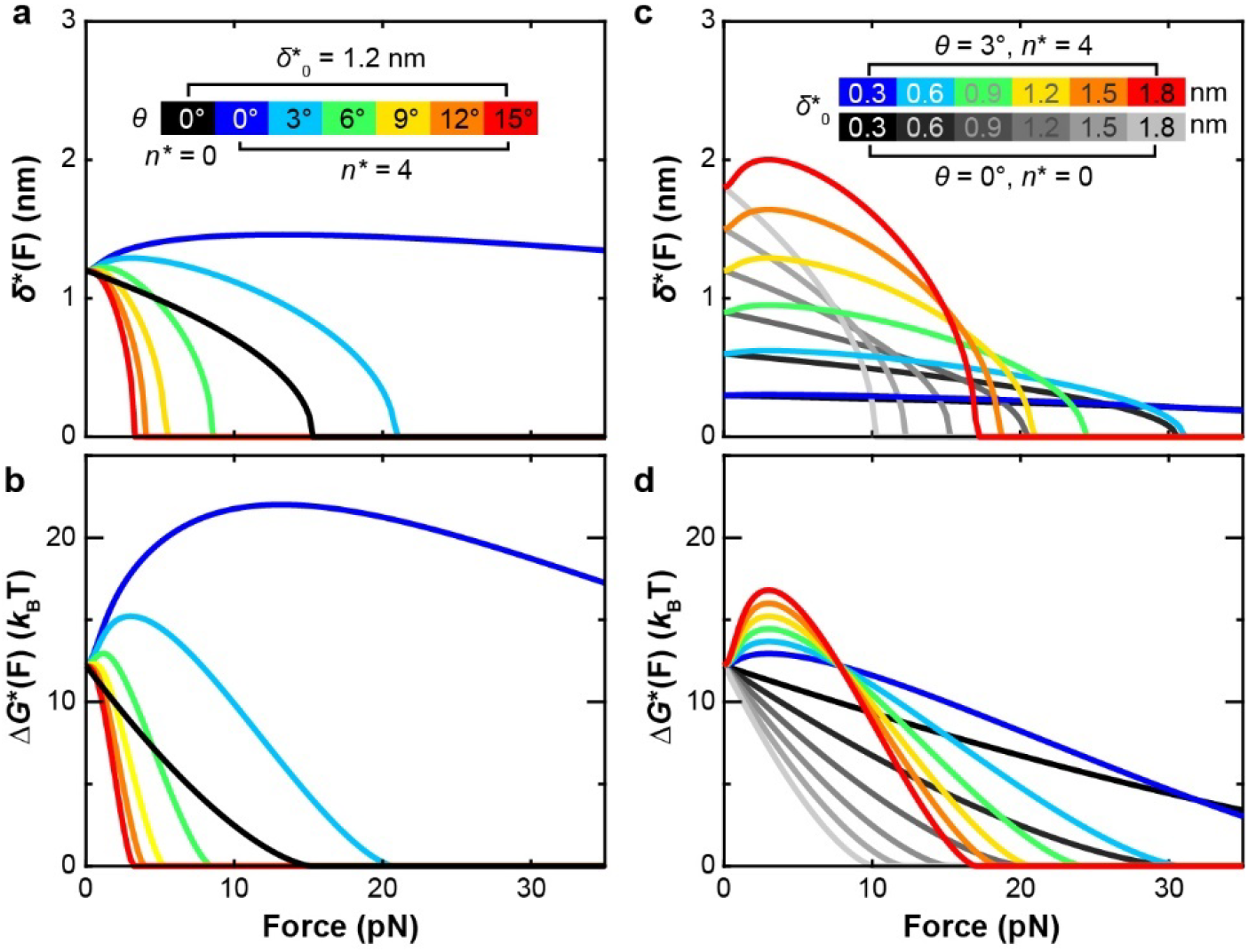
Characterization of the energy landscape of TCR–pMHC-II dissociation. **a-d** Plots of transition state location ***δ***^*^ (**a, c**) and height of energy barrier Δ***G***^*^ (**b, d**) *vs* force ***F*** for changing ***θ*** and ***n***^*^ (**a, b**) or 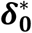 and ***n***^*^ (**c, d**) while keeping 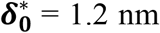 (**a, b**) or ***θ*** = 0, 3° (**c, d**). Inset color bars indicate parameter values used to plot the color-match theoretical curves.

**Supplementary Figure 8.**
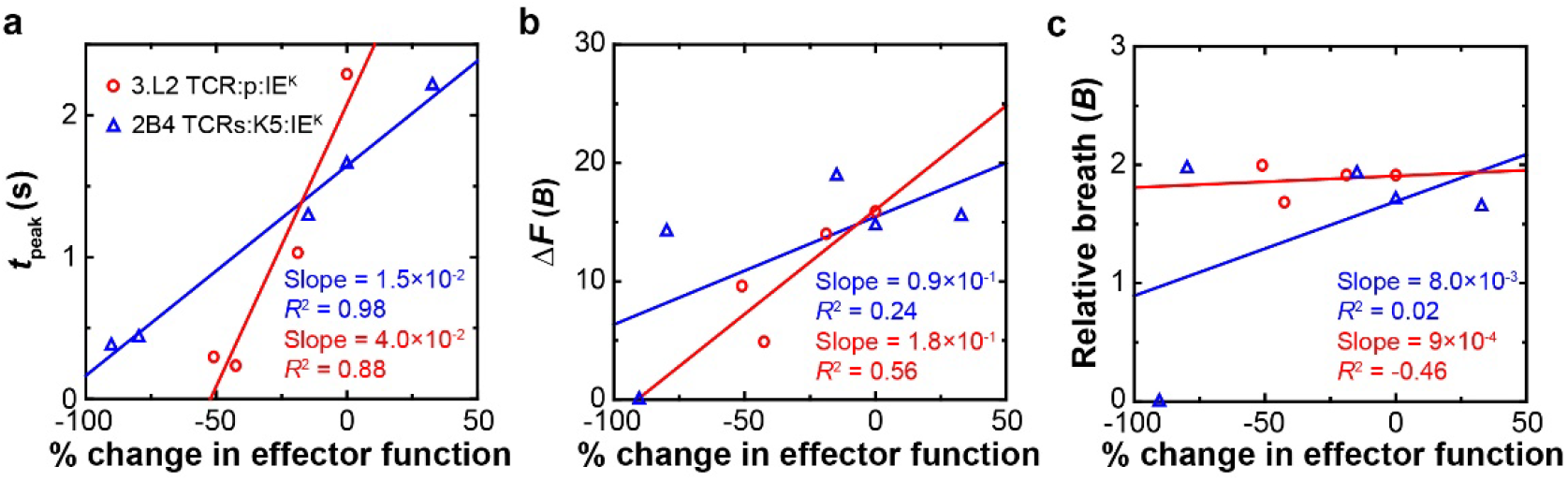
Correlation of metrics of TCR–pMHC-II bond lifetime *vs* force curves with T cell biological activity. **a-c** Dimensional metrices, ***t***_**peak**_ (**a**), Δ***F*** (**b**) and relative breadth of catch-slip bond ***B*** (**c**) *vs* reciprocal % change (relative to WT) in T cell effector function, i.e., the peptide dose required for 3.L2 T cells to generate 40% B cell apoptosis (1/EC_40_)^**20**^ (*red*) or the area under the dose response curve (AUC) of 2B4 hybridoma IL-2 production^**21**^ (*blue*) plots.

**Supplementary Figure 9.**
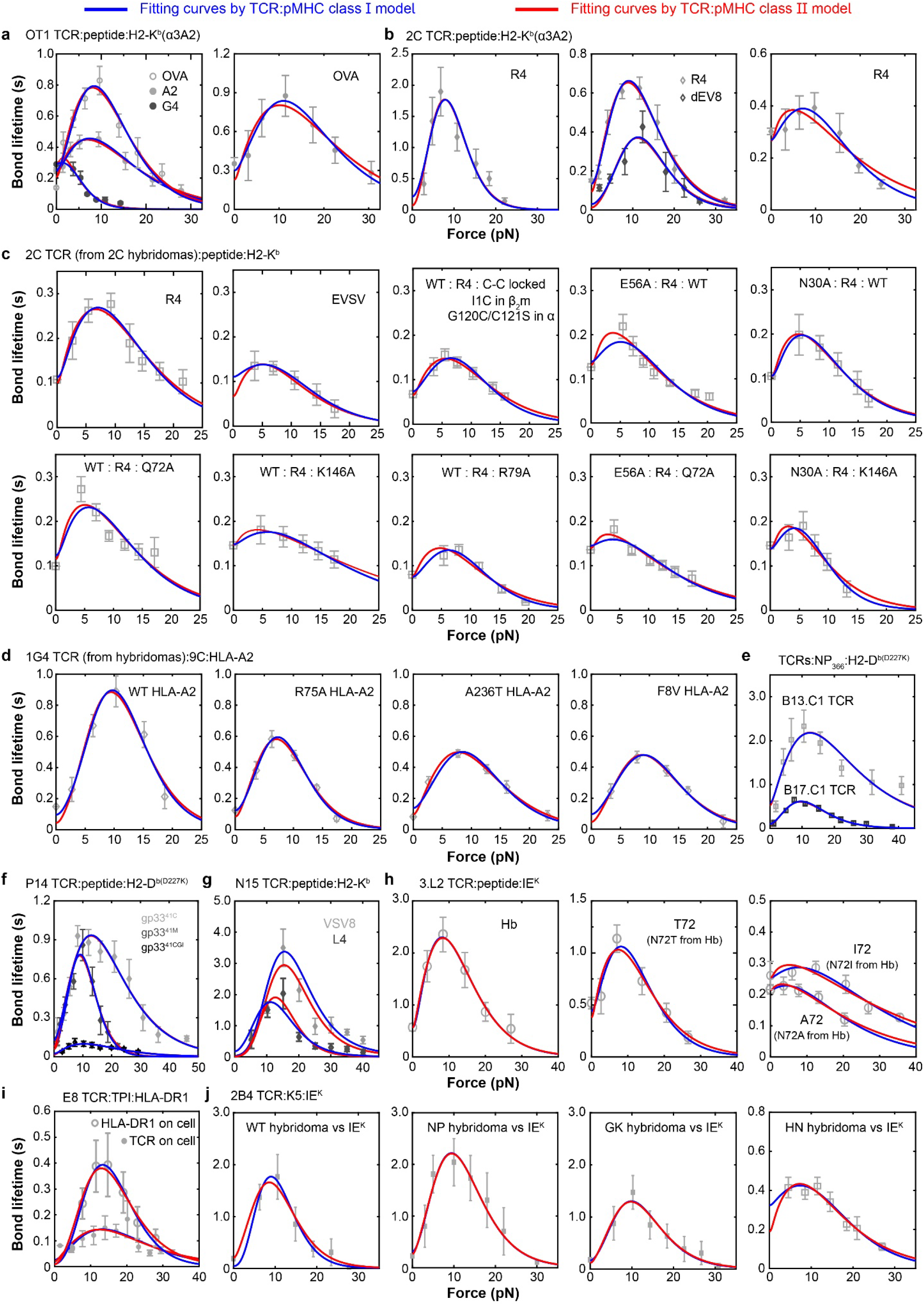
Comparing fitting curves of class I and class II models to the same data. **a-j** Fitting of theoretical 1/*k*(*F*) curves predicted by the class I (blue) or class II (*red*) model to experimental lifetime (*points*, mean ± sem) *vs* force data of TCR bonds with pMHC-I (**a-g**) or pMHC-II (**h-j**) ligands, as described below (partly presented in Supplementary Fig. 1, re-analyzed from Refs. ^4,15-19,22-24^): OT1 (**a**) or 2C (**b**) TCR expressed on either CD8^+^ naïve T cells or CD4^+^CD8^+^ thymocytes interacting with indicated p:H2-K^b^α3A2; WT or mutant 2C (**c**) or 1G4 (**d**) TCR expressed on hybridomas interacting with indicated peptides presented by WT or MT H2-K^b^ or HLA-A2. B13.C1 and B17.C1 TCR expressed on hybridomas interacted with NP_366_ bound to the D227K MT of H-2D^b^ to prevent CD8 binding (**e**). P14 TCR expressed on CD8^+^ naïve T cells interacting with indicated the D227K MT of H-2D^b^ to prevent CD8 binding (**f**). Soluble N15 TCRαβ interacting with indicated p:H2-K^b^ (**g**). 3.L2 TCR expressed on CD4^-^CD8^+^ T cells interacting with indicated p:I-E^k^ (**h**). E8 TCR expressed on CD4^-^ Jurkat cells interacting with TPI:HLA-DR1 or TPI:HLA-DR1 expressed on THP-1 cells interacting with E8 TCR (**i**). WT or MT 2B4 TCRs expressed on CD4^-^ hybridomas interacting with K5:I-E^k^ (**j**). See Supplementary Table 3 and 6 for lists of the interacting molecules.

**Supplementary Figure 10.**
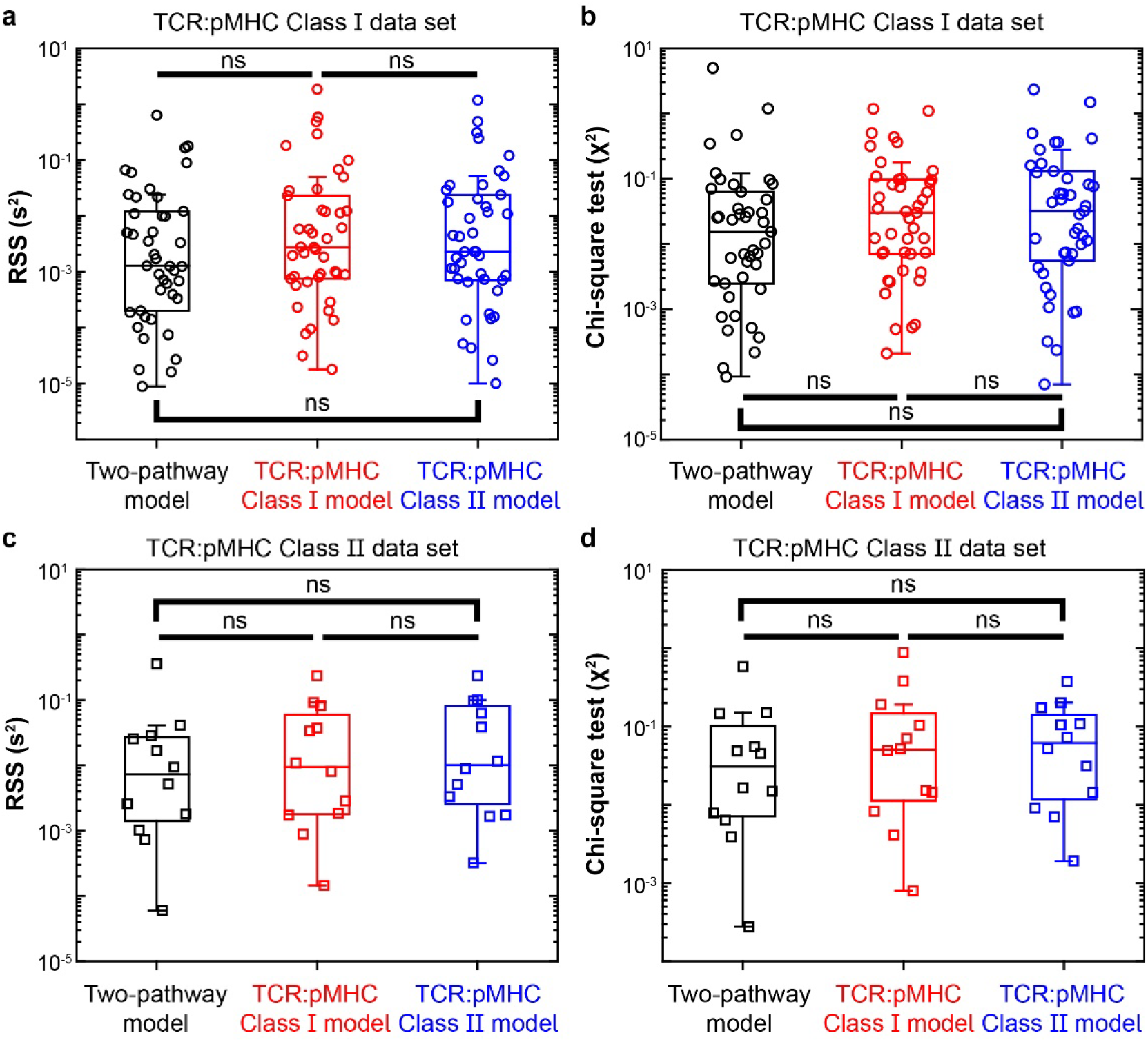
Comparison of goodness-of-fit measures among the class I and II models and the two-pathway model to class I and class II data. **a-b** Residual sum of squares (RSS, **a**) and Chi-square values (***χ***^**2**^, **b**) obtained using three models – two-pathway model (*black*), TCR–pMHC-I model (red), and TCR–pMHC-II model (*blue*) – to fit pMHC-I data. **c-d** Residual sum of squares (RSS, **c**) and Chi-square values (***χ***^**2**^, **d**) obtained using three models – two-pathway model (*black*), TCR-pMHC-I model (red), and TCR-pMHC-II model (*blue*) – to fit pMHC-II data. Individual scattered points, median, 25%, 75% and 5%, 95% whiskers are shown. Statistical tests were done using two-tailed paired *t* test.

**Supplementary Figure 11.**
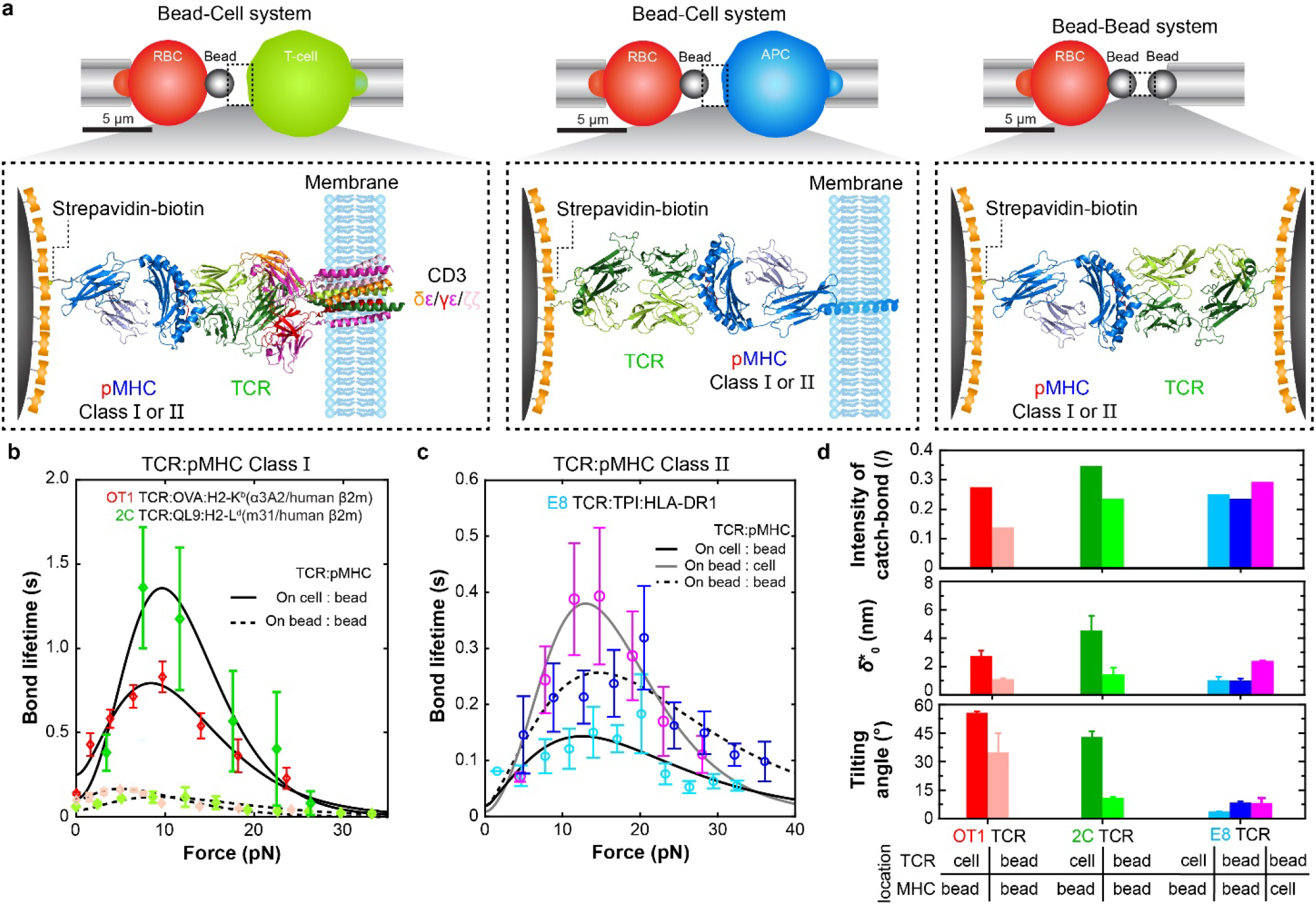
Comparison of pMHC interactions with cell surface and soluble TCRs. **a** Schematics of three biomembrane force probe (BFP) experiments. The BFP were set up in three configurations (*upper row*) depending on the molecular systems tested (*lower row*). In the first configuration (*left column*), the pMHC was coated on a BFP bead (*left*) and the TCR-CD3 complex was expressed on a T cell (*right*). In the second configuration (*middle column*), the purified TCRαβ was coated on a BFP bead (*left*) and the pMHC was expressed on an APC (*right*). In the third configuration, the pMHC was coated on a BFP bead (*left*) and the purified TCRαβ was coated on a glass bead (*right*). The TCR–pMHC complexes were drawn based on the cryoEM structure with CD3 (6JXR) or crystal structure without CD3 (2CKB). **b-c** Fitting of theoretical 1/*k*(*F*) (*curves*) to experimental bond lifetime *vs* force data (*points*, mean ± sem) of the following interactions: OT1 or 2C TCR respectively expressed on CD8^+^ naïve T cells (*red*) or CD8^-^ hybridomas (*green*) or coated on beads (*pink* for OT1 and *yellow-green* for 2C) respectively interacting with OVA:H2-K^b^α3A2 or QL9:H2-L^d^(m3), both MHC class I molecules^**22**^ (**b**) or of E8 TCR expressed on CD8^-^ Jurkat (*sky-blue*) or coated on beads interacting with TPI:HLA-DR1, a MHC class II, coated on beads (*blue*) or expressed on THP-1 cells (*purple*)^**24**^ (**c**). **d** Comparison of the catch bond intensity (*top*) and best-fit parameters 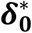 (*middle*) and ***θ*** (*bottom*) evaluated from data in b and c of TCR–pMHC interactions measured using the three configurations depicted in a. All error bars indicate fitting errors. See Supplementary Table 3 and 6 for lists of the interacting molecules.

**Supplementary Table 1.** Summary of model constants.

| Parameters | Symbol | Value | Reference |
| --- | --- | --- | --- |
| Elastic modulus of the folded globular domain | $E_d$ | 100 pN | <sup>2</sup> |
| Average contour length | $l_c$ | 0.36 nm | <sup>6-8</sup> |
| Persistence length | $l_p$ | 0.39 nm | <sup>6-8</sup> |
| Elastic modulus of polypeptides | $E_p$ | 50 μN | <sup>8</sup> |

| TCR : peptide : MHC | $d_{N,c}$ (nm)<sup>‡</sup> | $d_{C\alpha\beta,c}$ (nm)<sup>*</sup> | $\varphi$ (°)<sup>‡</sup> |
| --- | --- | --- | --- |
| OT1:peptide:H2-K<sup>b</sup> | 11.6 | 3.4 | 23.6 |
| 2C:peptide:H2-K<sup>b</sup> | 12.7 | 3.7 | 13.8 |
| 2C:peptide:H2-L<sup>d</sup>(m3) | 10.9 | 3.7 | 5 |
| 1G4:peptide:HLA-A2 | 12.3 | 3.7 | 13.3 |
| P14:peptide:H2-D<sup>b</sup> | 12.7 | 3.4 | 15 |
| B13.C1, B17.C1, B17.R1, B17.R2:peptide:H2-D<sup>b</sup> | 12.5 | 3.4 | 15 |
| N15:peptide:H2-K<sup>b</sup> | 11.6 | 3.4 | 23.6 |
| 3.L2:peptide:I-E<sup>k</sup> | 12.3 | 3.4 | N.A. |
| E8:peptide:HLA-DR1 | 12.3 | 3.4 | N.A. |
| 2B4:peptide:I-E<sup>k</sup> | 12.3 | 3.4 | N.A. |

<sup>‡</sup> Force-free extension of bound state
<sup>\*</sup> Force-free length of TCR $\alpha\beta$ constant domain
<sup>‡</sup> Angle shown in Fig. 2b and Supplementary Figure 4
N.A. indicates not applicable.

**Supplementary Table 2.**
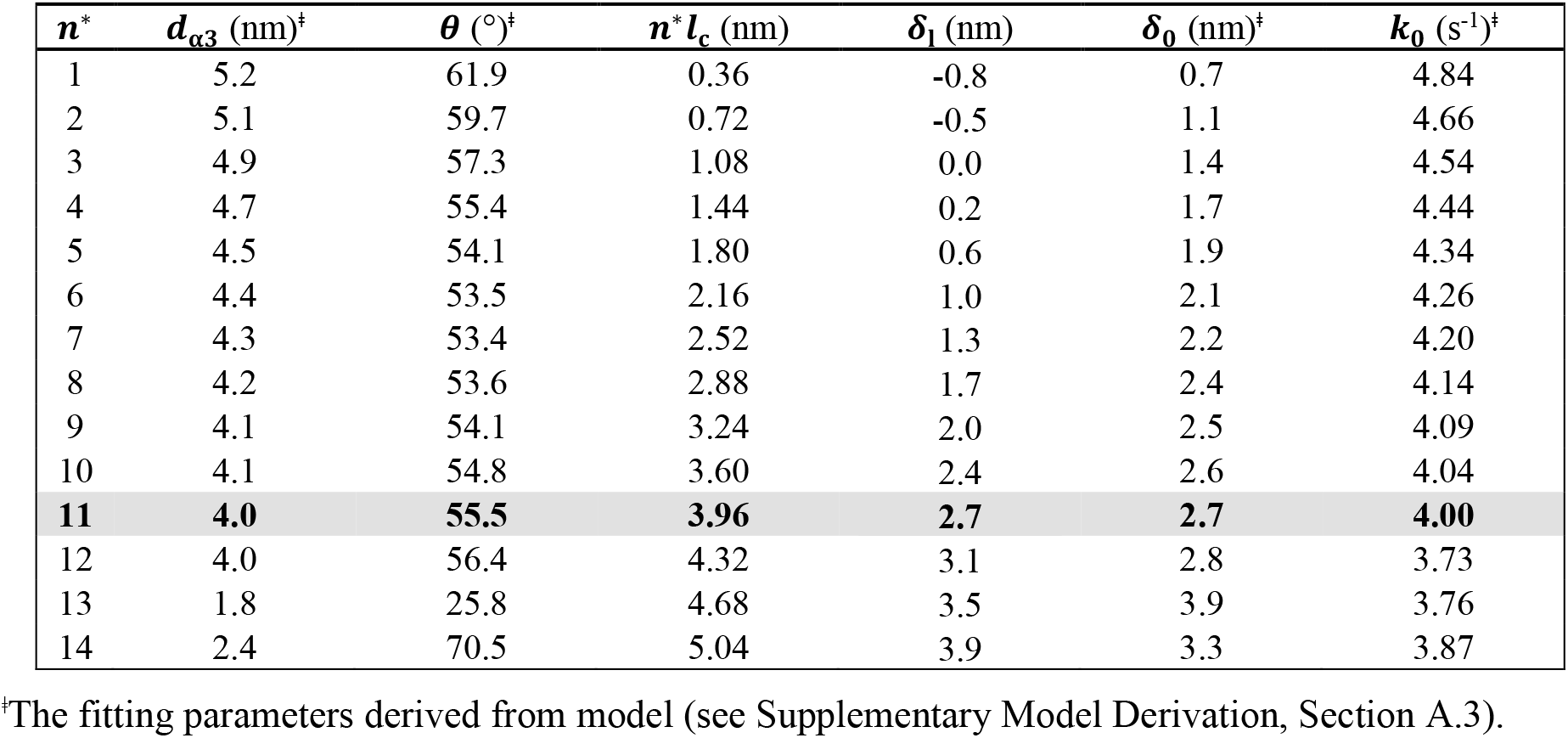
An example of finding the best-fit parameters for OT1 TCR– OVA:H2-K^b^a3A2 bond.

**Supplementary Table 3.**
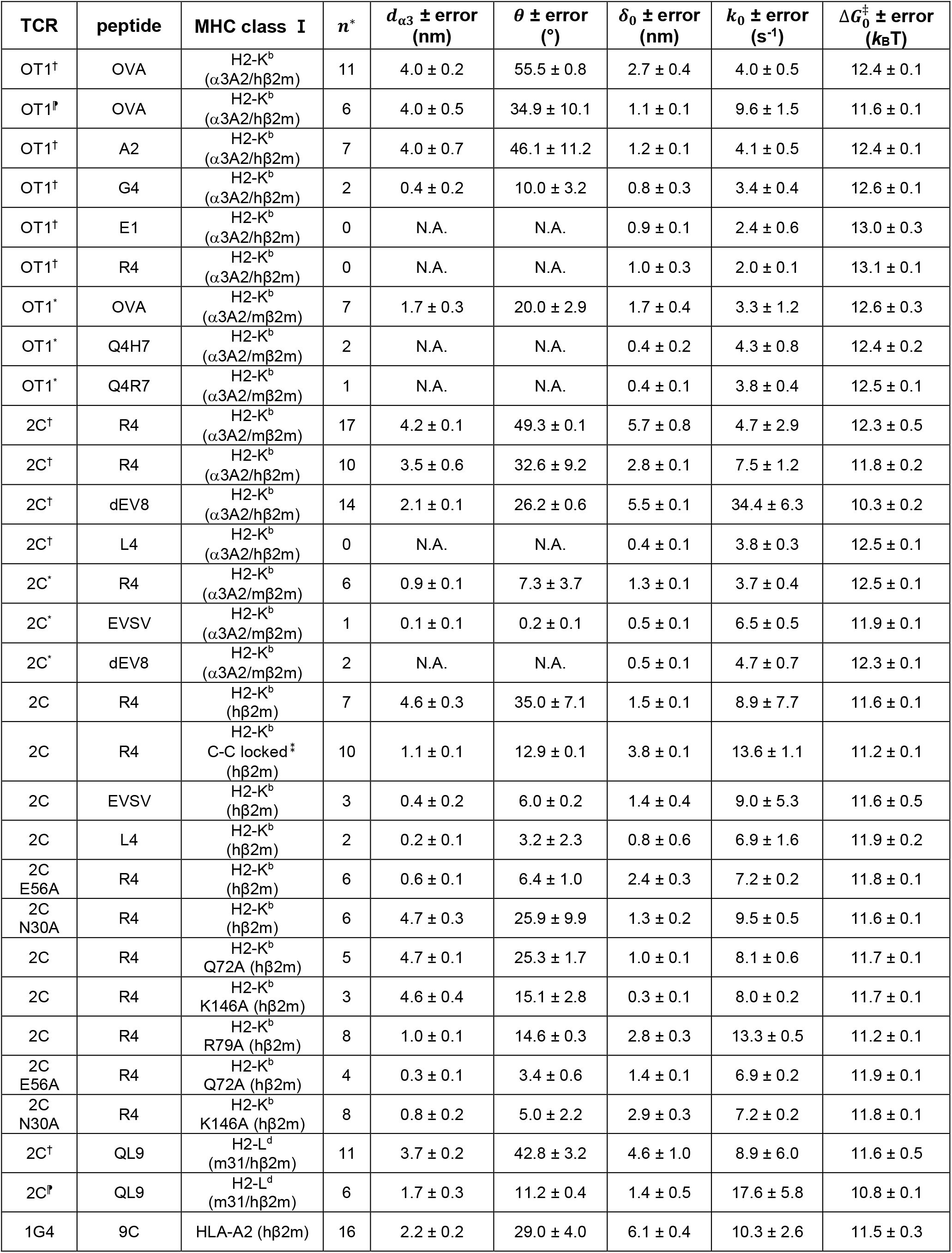
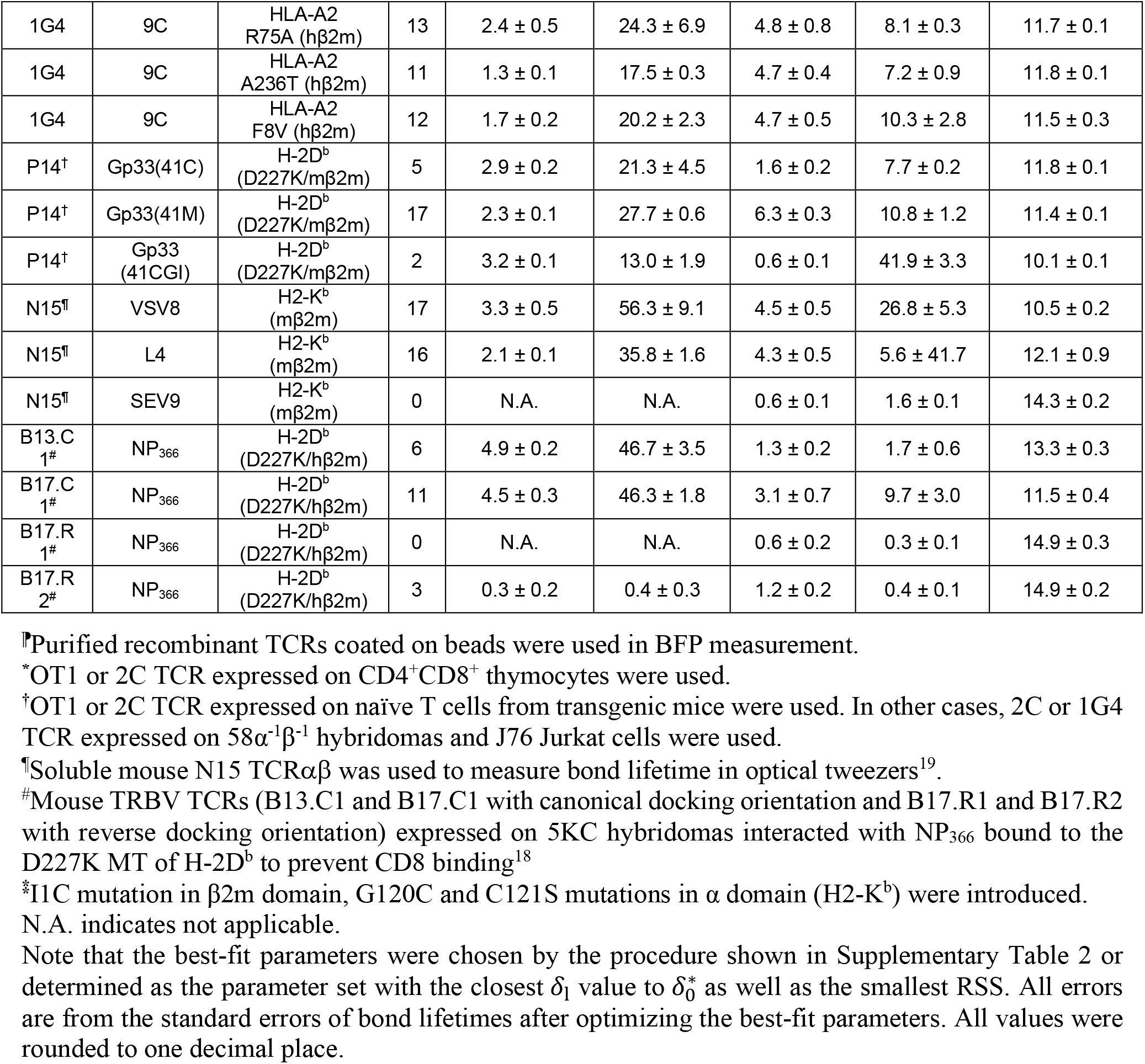
TCR–pMHC-I bond summary and their best-fitting parameters.

**Supplementary Table 4.**
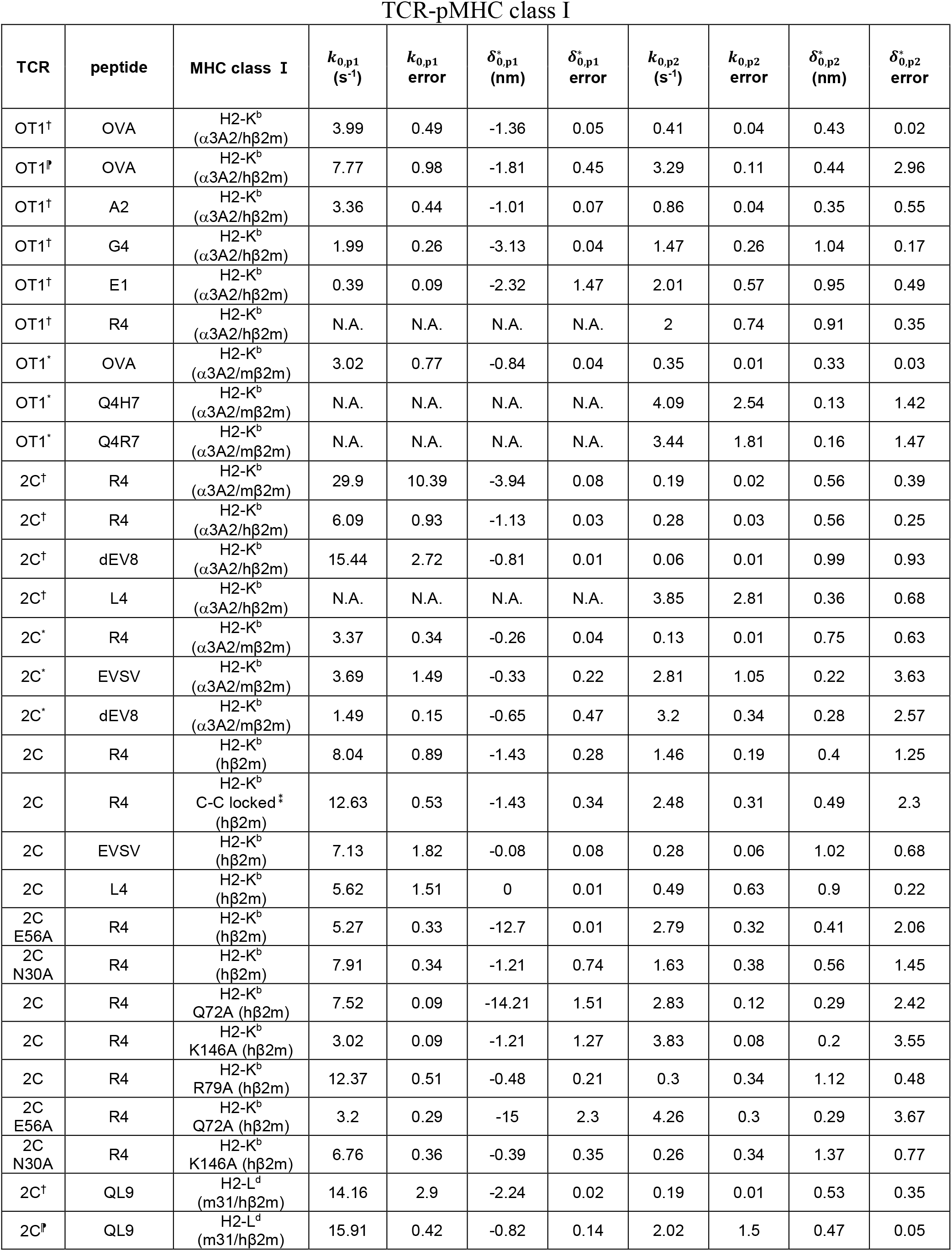
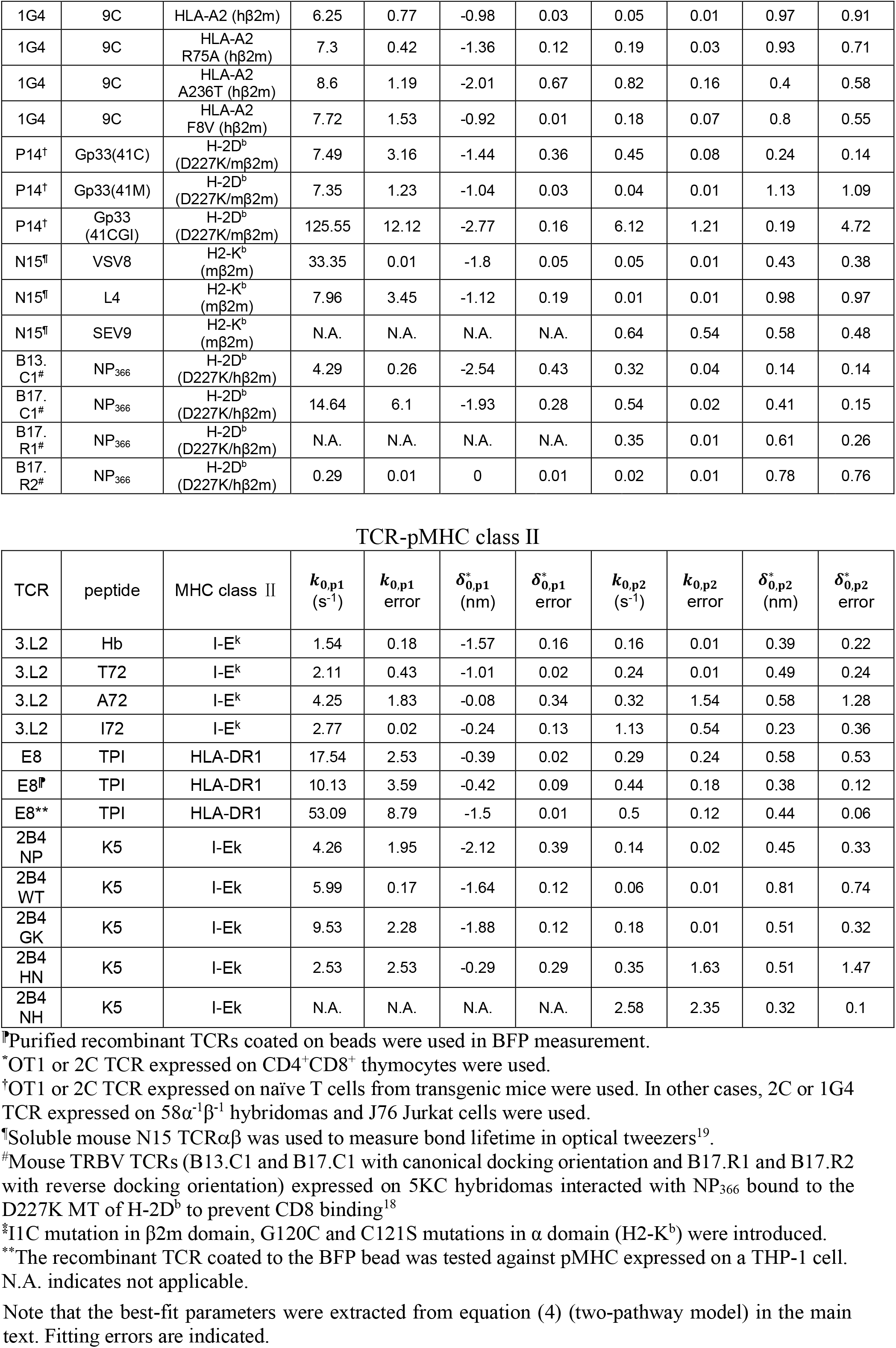
Two-pathway model summary and their best-fitting parameters.

**Supplementary Table 5.** An example of finding the best-fit parameters for the 3.L2 TCR–Hb:I-E^k^ bond.

TCR–Hb:I-E<sup>k</sup> bond
| $n_{\text{MHC}}^*$<sup>†</sup> | $d_{\text{B.I}}$ (nm)<sup>‡</sup> | $\theta$ (°)<sup>‡</sup> | $n^*l_c$ (nm) | $\delta_1$ (nm) | $\delta_0$ (nm)<sup>‡</sup> | $k_0$ (s<sup>-1</sup>)<sup>‡</sup> |
| --- | --- | --- | --- | --- | --- | --- |
| 1 | 8.9 | 0.0 | 0.4 | 0.6 | 0.7 | 2.0 |
| 2 | 8.8 | 0.0 | 0.7 | 0.9 | 1.2 | 1.9 |
| 3 | 8.6 | 0.5 | 1.1 | 1.3 | 1.5 | 1.8 |
| **4** | **8.0** | **3.2** | **1.4** | **1.6** | **1.6** | **1.8** |
| 5 | 7.9 | 2.6 | 1.8 | 2.0 | 1.7 | 1.7 |
| 6 | 7.8 | 2.1 | 2.2 | 2.4 | 1.9 | 1.7 |
| 7 | 8.0 | 1.5 | 2.5 | 2.7 | 2.1 | 1.6 |
| 8 | 7.8 | 0.6 | 2.9 | 3.1 | 2.2 | 1.6 |

<sup>†</sup> $n^*$ can be estimated from $n_{\text{pMHC}}^*$ .
<sup>‡</sup>See Supplementary Model Derivations, Section B.

**Supplementary Table 6.** TCR–pMHC-II bond summary and their best-fit parameters.

| TCR | peptide | MHC class II | $n^*$ | $d_{B,I} \pm \text{error}$<br>(nm) | $\theta \pm \text{error}$<br>(°) | $\delta_0 \pm \text{error}$<br>(nm) | $k_0 \pm \text{error}$<br>(s<sup>-1</sup>) | $\Delta G_0^\ddagger \pm \text{error}$<br>(k<sub>B</sub>T) |
| --- | --- | --- | --- | --- | --- | --- | --- | --- |
| 3.L2 | Hb | I-E<sup>k</sup> | 8 | 8.0 ± 0.2 | 3.2 ± 1.5 | 1.6 ± 0.3 | 1.8 ± 0.2 | 13.2 ± 0.1 |
| 3.L2 | T72 | I-E<sup>k</sup> | 6 | 8.7 ± 0.1 | N.A. | 1.2 ± 0.1 | 2.6 ± 0.3 | 12.9 ± 0.1 |
| 3.L2 | A72 | I-E<sup>k</sup> | 2 | 9.3 ± 0.2 | N.A. | 0.2 ± 0.1 | 4.7 ± 0.2 | 12.3 ± 0.1 |
| 3.L2 | I72 | I-E<sup>k</sup> | 2 | 9.1 ± 0.1 | N.A. | 0.1 ± 0.0 | 3.9 ± 0.3 | 12.4 ± 0.1 |
| E8 | TPI | HLA-DR1 | 4 | 8.5 ± 0.1 | 3.8 ± 0.4 | 1.0 ± 0.3 | 50.2 ± 4.8 | 9.9 ± 0.1 |
| E8<sup>†</sup> | TPI | HLA-DR1 | 4 | 7.9 ± 0.1 | 8.5 ± 0.5 | 1.1 ± 0.3 | 51.0 ± 10.5 | 9.9 ± 0.3 |
| E8<sup>*</sup> | TPI | HLA-DR1 | 8 | 7.6 ± 0.2 | 8.4 ± 2.4 | 2.4 ± 0.4 | 114.0 ± 64 | 9.1 ± 0.1 |
| 2B4 (NP) | K5 | I-E<sup>k</sup> | 14 | 7.5 ± 0.3 | 6.7 ± 1.2 | 2.5 ± 0.2 | 3.7 ± 0.2 | 12.5 ± 0.1 |
| 2B4 (WT) | K5 | I-E<sup>k</sup> | 16 | 7.5 ± 0.4 | 6.1 ± 1.1 | 2.9 ± 0.4 | 4.9 ± 6.7 | 12.2 ± 0.9 |
| 2B4 (GK) | K5 | I-E<sup>k</sup> | 10 | 8.3 ± 0.5 | 0.3 ± 0.2 | 2.6 ± 0.3 | 8.4 ± 1.2 | 11.7 ± 0.2 |
| 2B4 (NH) | K5 | I-E<sup>k</sup> | 0 | N.A. | N.A. | 0.3 ± 0.1 | 2.6 ± 0.6 | 12.9 ± 0.3 |
| 2B4 (HN) | K5 | I-E<sup>k</sup> | 4 | 8.9 ± 0.1 | 0.3 ± 0.1 | 0.7 ± 0.3 | 5.3 ± 0.5 | 12.2 ± 0.1 |

<sup>†</sup>The recombinant TCR coated to the bead was tested against pMHC on the BFP bead.
<sup>\*</sup>The recombinant TCR coated to the BFP bead was tested against pMHC expressed on a THP-1 cell.
N.A. indicates not applicable.
Note that the best-fit parameters were chosen using the procedure shown in Supplementary Table 5 or determined as the parameter set with the $\delta_z(n^*)$ value closest to $\delta_0^*$ as well as the smallest RSS. All errors are standard error of mean after optimizing the best-fit parameters. All values were rounded to one decimal place.

## Supplementary Movie

**Supplementary Movie. 1** | **SMD simulated trajectories for pulling the 2C TCR–SIYR:H2-K**_**b**_ **complex**. The C-terminal residue of H-2K^b^ was harmonically constrained and the C terminus of TCR was pulled by a dummy spring moving at ∼0.1 nm/ns with a spring constant of ∼70 pN/nm. The color codes were the same as that in Figure 7a,b and 2C TCRα (blue), 2C TCRβ (red), MHC H-2K^b^ α (orange), MHC H-2K^b^ β2m (cyan), and R4 peptide (green). Three different pathways of conformational changes were observed in the pMHC.

**Supplementary Movie. 2** | **CMD and SMD simulated trajectories side-by-side comparison of pulling effect on the human TCR-CD3 complex focusing on interaction of P223 (TCRβ) and L90 (CD3ε)**. The left panel shows the freely evolved fluctuations without force from CMD, and the right panel presents the constrained dynamics with force from SMD. For SMD trajectory, the external force of 175pN was exerted on the N-terminal of TCRα, pulling upwards while keeping the C-termini of CD3ε anchored. The color codes were the same as that in Fig. 7g: TCRα (yellow-green), TCRβ (green), and CD3 subunits (golden). The focused interacting residues were highlighted using spheres, where P223β (red) was pulled closer to L90ε (gray) in the presence of force.

**Supplementary Movie. 3** | **CMD and SMD simulated trajectories side-by-side comparison of pulling effect on the human TCR-CD3 complex focusing on interaction of K183 (TCRβ) and L90 (CD3ε’)**. The left panel shows the freely evolved fluctuations without force from CMD, and the right panel presents the constrained dynamics with force from SMD. For SMD trajectory, the external force of 175pN was exerted on the N-terminal of TCRα, pulling upwards while keeping the C-termini of CD3ε anchored. The color codes were the same as that in Fig. 7g: TCRα (yellow-green), TCRβ (green), and CD3 subunits (golden). The focused interacting residues were highlighted using spheres, where K183β (blue) was pulled away from L90ε’ (gray) in the presence of force.

**Supplementary Movie. 4** | **CMD and SMD simulated trajectories side-by-side comparison of pulling effect on the human TCR-CD3 complex focusing on interaction of N225 (TCRβ) and E38 (CD3γ)**. The left panel shows the freely evolved fluctuations without force from CMD, and the right panel presents the constrained dynamics with force from SMD. For SMD trajectory, the external force of 175pN was exerted on the N-terminal of TCRα, pulling upwards while keeping the C-termini of CD3ε anchored. The color codes were the same as that in Fig. 7g: TCRα (yellow-green), TCRβ (green), and CD3 subunits (golden). The focused interacting residues were highlighted using spheres, where N225β (purple) was pulled away from E38γ (gray) in the presence of force.

## Notes

### Competing Interest Statement

The authors have declared no competing interest.

### Summary of Updates

This version includes additional experimental/simulation data as well as additional analysis to improve the impact of the study.

